# Evolutionary convergence of sensory circuits in the pallium of amniotes

**DOI:** 10.1101/2024.04.30.591819

**Authors:** Eneritz Rueda-Alaña, Rodrigo Senovilla-Ganzo, Marco Grillo, Enrique Vázquez, Sergio Marco-Salas, Tatiana Gallego-Flores, Artemis Ftara, Laura Escobar, Alberto Benguría, Ana Quintas-Gorozarri, Ana Dopazo, Miriam Rábano, María dM Vivanco, Ana María Aransay, Daniel Garrigos, Ángel Toval, José Luis Ferrán, Mats Nilsson, Juan Manuel Encinas, Maurizio De Pitta, Fernando García-Moreno

## Abstract

The amniote pallium contains sensory circuits structurally and functionally equivalent, yet their evolutionary relationship remains unresolved. Our study employs birthdating analysis, single-cell RNA and spatial transcriptomics, and mathematical modeling to compare the development and evolution of known pallial circuits across birds (chick), lizards (gecko) and mammals (mouse). We reveal that neurons within these circuits’ stations are generated at varying developmental times and brain regions across species, and found an early developmental divergence in the transcriptomic progression of glutamatergic neurons. Together, we show divergent developmental and evolutionary trajectories in the pallial cell types of sauropsids and mammals. Our research highlights significant differences in circuit construction rules among species and pallial regions. Interestingly, despite these developmental distinctions, the sensory circuits in birds and mammals appear functionally similar, which suggest the convergence of high-order sensory processing across amniote lineages.

## Introduction

The neocortex is the biological substrate underlying the sophisticated and complex functions performed by mammalian species including humans. Neocortical neurons connect to each other following a stereotyped pattern (*1*, *2*). This canonical circuit is comprised of three types of pallial glutamatergic neurons, each fulfilling a specific role in the circuitry (**Fig. 1A**). Electrophysiological evidence supports that these circuits account for major neocortical functions (*3*) and could be a major breakthrough of mammalian evolution. However, other vertebrate species that do not display a neocortex have been shown to present corresponding circuits. In both birds and reptiles, sensory information is also processed within circuits comprised of three glutamatergic neurons, contained within the pallium. The evolutionary origin of these amniote circuits is a matter of intense debate. These could be homologues to neocortical circuits -and so derived from the same ancestral circuit- or analogues, having convergently evolved separately (*4–6*) .

**Fig. 1.**
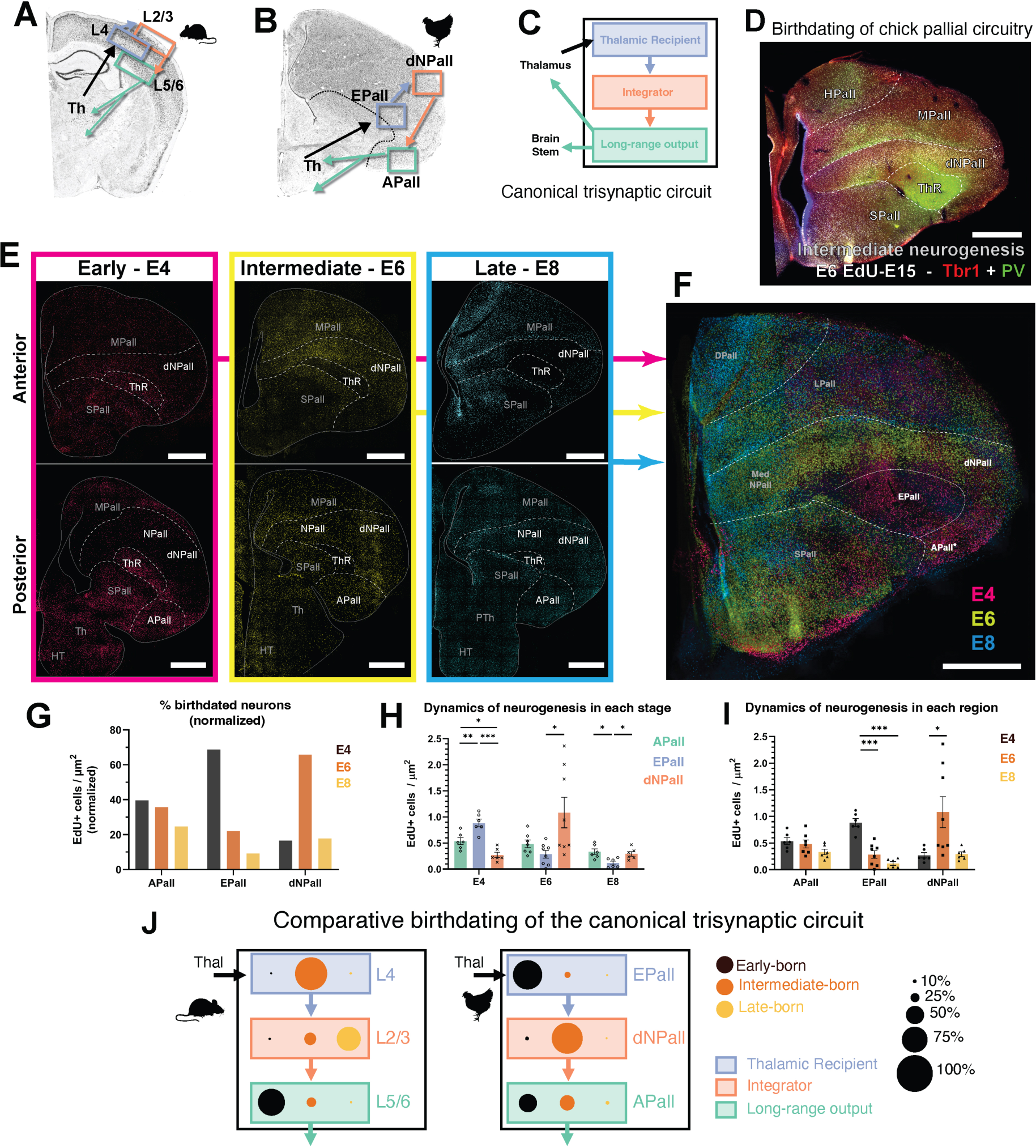
Neuronal birthdating of the chick DVR sensory circuit. (**A**) Schematic highlighting the connections of the mammalian cortical canonical circuit. Th: thalamus. (**B**) Equivalent schematic depicting the connections of the chick DVR canonical circuit. (**C**) Scheme of the basic cell type organization in the trisynaptic circuit of amniotes. MPall: mesopallium; SPall: subpallium; ThR: thalamic recipient area. (**D**) Example of coronal section of the chick telencephalon in a typical experiment, where EdU is labeled (white) along with two markers by immunohistochemistry, PV (green) and Tbr1 (red). (**E**) Summary of the birthdating performed at early (E4, magenta), intermediate (E6, gold) or late (E8, cyan) neurogenic timepoints. Two coronal sections are shown, one anterior (top row) and one at posterior levels (bottom row). HT: hypothalamus; PTh: prethalamus. (**F**) Pseudoimage made by the merge of images coming from three different animals, showing the distribution of pallial cells according to birth time. (**G-I**) Quantitative analysis of the neurogenesis of DVR circuit cells. (**G**) Graphic representation of the temporal neurogenic construction of the different DVR circuit roles. The graph illustrates the normalized percentage of cells of each circuit role generated at early, intermediate or late neurogenic timepoints. (**H**) Graphic representation of the distribution of DVR circuit neurons generated at each analyzed neurogenic timepoint. (**I**) Graphic representation of the effect of age in the generation of the cells of each DVR circuit roles. Statistical tests for (**I, J**) are described in **Supplementary Materials.** (**J**) Summary of the comparative birthdating of mammalian cortical vs. avian DVR sensory circuits. The graph depicts the percentage of cells of each circuit role generated at early, intermediate or late neurogenic timepoints. Scale bars, 1 mm.

The relevant circuits in the three vertebrate groups - mammals (*1*), birds (*7*, *8*) and non-avian reptiles (*9*)- are functionally similar. Briefly, in all three groups there is a specific neuronal population which receives thalamic inputs. These neurons connect to an integrator neuron which, in turn, sends its input to a third pallial neuron. This last neuron of the circuit projects the information out of the cortex, to either the thalamus, the brainstem or other subcortical structures. Beyond similarities in their location, type of information processed and neurotransmitters, the neuronal populations of these circuits share the expression of a handful transcription factors -such as Tbr1, RorB, Satb2 or Ctip2 (*9–12*)- and electrophysiological properties (*13*). Due to these functional and structural similarities (*14*), it has been suggested and maintained that all these amniote circuits are homologous (*15*), implying that an equivalent ancestral circuit was already present in the brain of the last common ancestor of amniotes around 320mya. This view contrasts with evidence from developmental biology which propose a divergent scenario for the evolution of the pallium (*16*, *17*) as these circuits appear in developmentally non-comparable brain regions.

Development and evolution are tightly linked: variations in one of them dictate trends in the other one (*18*, *19*). Because evolutionary homology implies inheritance from a common ancestor, the developmental trajectory must be conserved for arguing the conservation of homologous neurons and circuits. Here, we argue that understanding the developmental trajectory of a neuron or a circuit may unravel their evolutionary history (*20*, *21*). Conserved homologous cell populations are those who share an equivalent developmental program. Therefore, the long-standing debate over the homology of the sensory circuit can only be resolved by comparing the developmental trajectories of neurons in these amniote circuits. To investigate this evolutionary relationship and potential homology, we compared the developmental and transcriptional trajectories over the course of the construction of pallial sensory circuits in three amniote species.

## Results

### Developmental construction of avian high-order sensory circuits

Conserved homologous cells share their developmental history. To test whether mammalian cortical circuits have homologous circuits in other amniote species we determined the developmental trajectory of pallial sensory circuits in sauropsids. The temporal neurogenic construction of the mammalian cortical circuit is well known. By means of EdU injections at three different time points during the neurogenic period of the chick pallium (E4 to E9) we revealed the temporal construction of the avian sensory circuits (**Figs. 1,2, S01-S10**). We researched on the two pallial structures that have been proposed as neocortical homologues, the dorsal ventricular ridge (DVR; **Figs. 1, S01-S04, 09, 10**) and the hyperpallium (HypPall; **Figs. 2, S05-S08**). We found that none of these structures develops in an equivalent temporal sequence as that known for the mammalian cortical circuit.

**Fig. 2.**
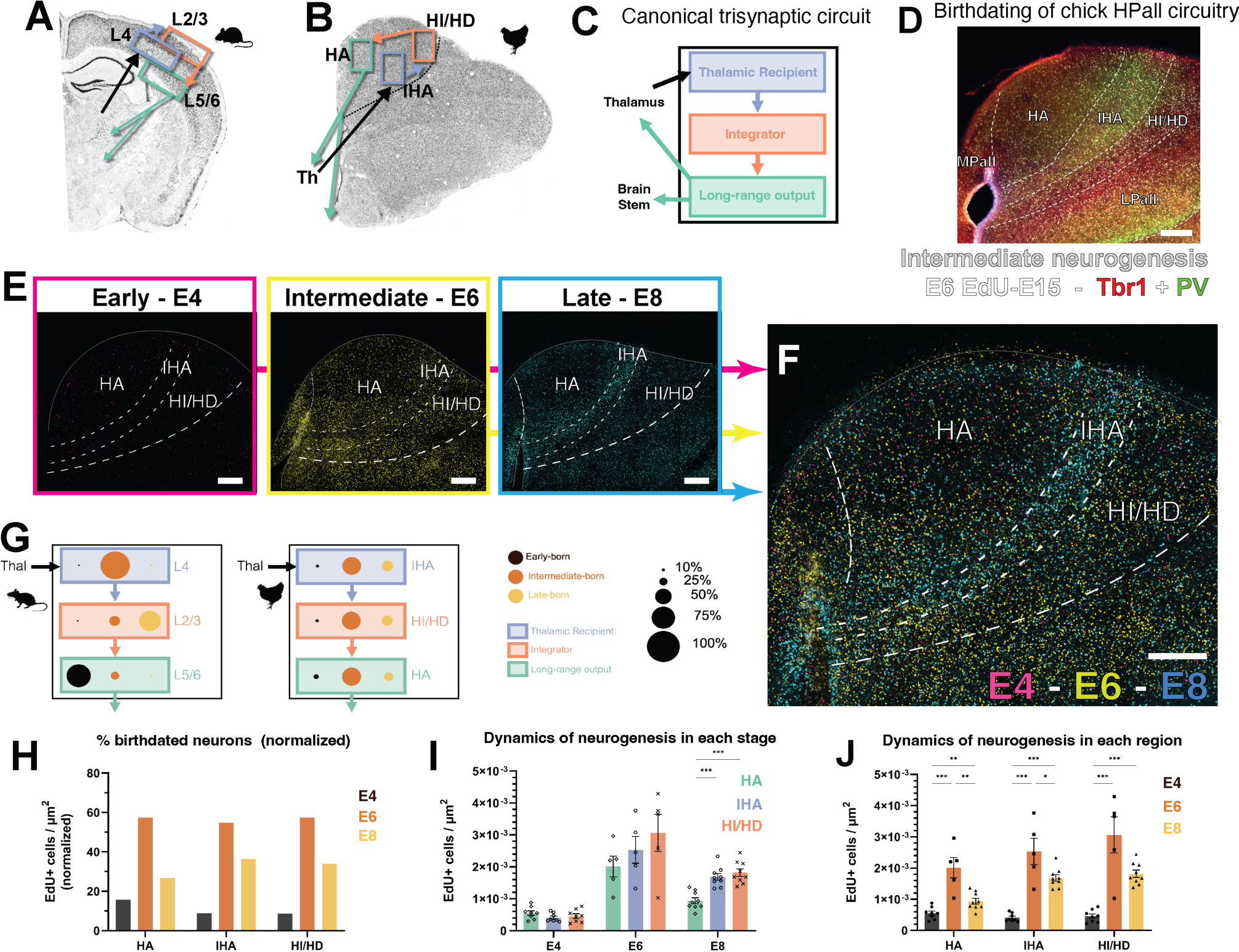
Neuronal birthdating of the chick hyperpallial sensory circuit. (**A**) Schematic highlighting the connections of the mammalian cortical canonical circuit. (**B**) Equivalent schematic depicting the connections of the chick hyperpallial canonical circuit. HA: hyperpallium apicale; HI/HD: intermediate and densocellular hyperpallium; IHA: intercalated hyperpallial area. (**C**) Scheme of the basic cell type organization in the trisynaptic circuit of amniotes. (**D**) Example of coronal section of the chick telencephalon in a typical experiment, where EdU is labeled (white) along with two markers by immunohistochemistry, PV (green) and Tbr1 (red). (**E**) Summary of the birthdating performed at early (E4, magenta), intermediate (E6, gold) or late (E8, cyan) neurogenic timepoints. (**F**) At the right, pseudoimage made by the merge of images coming from three different animals, showing the distribution of pallial cells according to birth time. (**G**) Summary of the comparative birthdating of mammalian cortical vs. avian hyperpallial sensory circuits. The graph depicts the percentage of cells of each circuit station generated at early, intermediate or late neurogenic timepoints. (**H-J**) Quantitative analysis of the hyperpallial circuit neurogenesis. (**H**) Graphic representations of the temporal neurogenic construction of the neurons of the different hyperpallial circuit roles. The graph illustrates the normalized percentage of cells of each circuit role generated at early, intermediate or late neurogenic timepoints. (**I**) Graphic representation of the distribution of hyperpallial circuit neurons generated at each analyzed neurogenic timepoint. (**J**) Graphic representation of the effect of age in the generation of the cells of each hyperpallial circuit roles. Statistical tests for (**I, J**) are described in **Supplementary Materials.** Scale bars, 250 µm.

First, we focused on the avian DVR, which was primarily considered the homologous of the neocortex according to its internal circuitry. A similarity with mammalian corticogenesis was that both the glutamatergic neurons and the GABAergic interneurons of each circuit relay were generated mostly synchronously (**Figs. S02-04**). However, DVR developed following different temporal neurogenic instructions to those present in the developing cortex. Two crucial differences were: First, the first-generated neurons act as thalamic recipient neurons within the entopallium (EPall) and the BSS (**Figs. 1E-J and S01**), whereas in mouse other cortical populations are generated earlier to thalamic recipient neurons of layer 4; and second, latest-generated neurons of the DVR do not contribute to sensory processing, as they remain in the most medial nidopallium (**Figs. 1E-J and S01**). In the case of mammals, all cortical neurons contribute to the circuitry, and the latest generated neurons act as integrators of the circuit in supragranular layers 2 and 3.

Beyond timing differences in the formation of the sensory neurons, we also found other dissimilarities between these two circuits. GABAergic interneurons, widely conserved amongst vertebrate pallia, distribute differently in these circuits. For instance, parvalbumin-expressing interneurons of the DVR are nearly exclusively located in the thalamic recipient areas (**Figs. 1 and S02-03**), whereas in mammals these neurons are widely distributed across all cortical layers. Somatostatin-expressing neurons, on the contrary, are dispersed through all regions of the avian circuit (**Figs. 1 and S04**), whereas are preferentially confined to deep cortical layers in mammals.

In addition, other developmental features in charge of building these circuits were different between mammalian cortex and avian DVR. While the neocortex derives from the dorsal pallial (DPall) germinative zone, the DVR derives from the ventral pallium (VPall; **Fig. 1**). This is a different, discrete region of the embryonic pallium that differs from DPall in the expression of key transcription factors, cell lineage and mature structures derived (in mammals, VPall mostly generates the pallial amygdala and the pyriform cortex)(*22*). These ancestral early developmental regions are so deeply conserved that only cell types within the same territory are considered homologous. Thus, neocortex and DVR derive from non-homologous progenitor populations.

The trisynaptic circuit in the DVR represents the building block of its internal circuitry but it oversimplifies the true connections, as it also occurs in neocortical circuits. Importantly, the ventral mesopallium plays a significant role as integrator in the circuitry, by receiving both EPall and dorsal nidopallium (dNPall) connections and due to its efferences towards the Arcopallium (APall)(*14*). We researched its main neurogenic waves to evaluate its similarities to integrator neurons of the neocortex (**Fig. S05**). The main neurogenic timepoint of the ventral mesopallium was the intermediate stages, similarly to the other integrator neurons of the dNPall. However, this is different to the late neurogenic timepoint of mammalian integrator neurons. And also, the ventral mesopallium is generated in the lateral pallium, a different developmental territory to the rest of the DVR circuit. These are all substantial developmental differences to the neocortical canonical circuit.

The DVR circuits described above mainly represent the visual and somatosensorial circuits present in the VPall. There is another sensorial circuit in the same developmental territory, the auditory Field L at the posterior nidopallium (*23*). We also assessed the sequence of neurogenesis of this circuit and found that most of its neurons were generated at late neurogenic timepoints (**Fig. S06**). This implies that this auditory circuits generates differently to other DVR circuits, and also different to the mammalian neocortical one.

We next examined the program of development of the HypPall. The mammalian neocortex and the avian HypPall are both DPall derivatives, for which the two structures are considered field homologous (*16,17*). We researched the same birthdated brains as for the DVR, but focusing on the neurogenic program of the HypPall. We also found key differences in the temporal generation of HypPall neurons (**Figs. 2 and S07-10**). HypPall neurogenesis happened much quicker than neocortical neurogenesis. Most HypPall neurons were generated during intermediate- and late-neurogenic stages (E6 and E8 stages in chick), and only a minor fraction of them were early generated. Both E6 and E8-generated HypPall neurons settled in all three HypPall functional columns (**Figs. 2 and S07-10**). Thus, differently from the mammalian cortex, HypPall neurons are not cell fate-determined by their time of generation. We did not identify any cell fate preference correlated to the neurogenic time of HypPall neurons (**Figs. 2G-J and S07-10**). This is a clear-cut difference to the neocortex, which shows a wide correlation between the time when a neocortical neuron is generated and its final function within the circuit. Interestingly, glutamatergic and GABAergic neurons of each HypPall column were generated synchronously at the same embryonic stage (**Figs. S07-10**), a shared feature with the DVR and the mammalian cortex.

In addition, the distribution of interneuron types was also different to that of mammalian cortical layers. The majority of parvalbumin-expressing neurons settled in the HI/HD (**Fig. S09**), a suggested homologous area of the integrator layers 2 and 3, whereas PV-expressing interneurons widely distribute through all cortical layers in mammals.

These data show evidence that the two main high order-sensory circuits in birds, DVR and HypPall, develop following a different program to that known in mammals ((**Figs. 1J and 2G**). That is, these circuits were not homologous and evolved exploiting different developmental instructions.

### Neurogenic time of gecko pallial sensory circuits

Because birds possess a highly specialized brain in the diapsid taxon (*24*), we further characterized the developmental construction of sensory circuits in a non-avian diapsid species such as the ground Madagascar gecko *Paroedura picta*. This species displays a pallium with fewer specializations, that better resembled the ancestral condition of the amniote group (*25*). The gecko pallium receives and processes thalamic sensory information in two main structures, the DVR and the dorsal cortex (DC). We identified these structures by markers such as Tbr1 or Satb2 (**Fig. 3A**; Satb2 antibody recognizes both Satb2- and Satb1- expressing neurons). We performed in situ hybridization to detect the expression of known markers for long-range output neurons (*Etv1* and *Sulf2,* **Fig. 3B,C**) thalamic recipient neurons (*Satb1*, **Fig. 3B,E**) (*26*, *27*), and so describe the location of the neurons executing these two functions. We also analyzed PV immunohistochemistry patterns, commonly expressed at the Rotundus nucleus in sauropsids, which is the main visual thalamic input to the DVR (**Fig. 3B,D**). Altogether, we identified the location of main types of gecko pallial neurons (**Fig. 3F**) and focus our attention on the crucial populations for the evolutionary comparison: in DC, long-range output cells settle layer 2, in line with available literature. Precisely, it has been shown than turtle DC neurons with higher similarity to cortical long-range output neurons locate the upper half of layer 2 (*11*). In addition, *Satb1* expression is nearly absent in layer 2, suggesting thalamic recipient neurons locate layer 3 (**Fig. 3E**). As for the DVR circuitry, we identified that the thalamic recipient neurons were located at the outer region of the DVR, according to *Satb1* expression, and to neuropil staining of PV from rotundal axons.

**Fig. 3.**
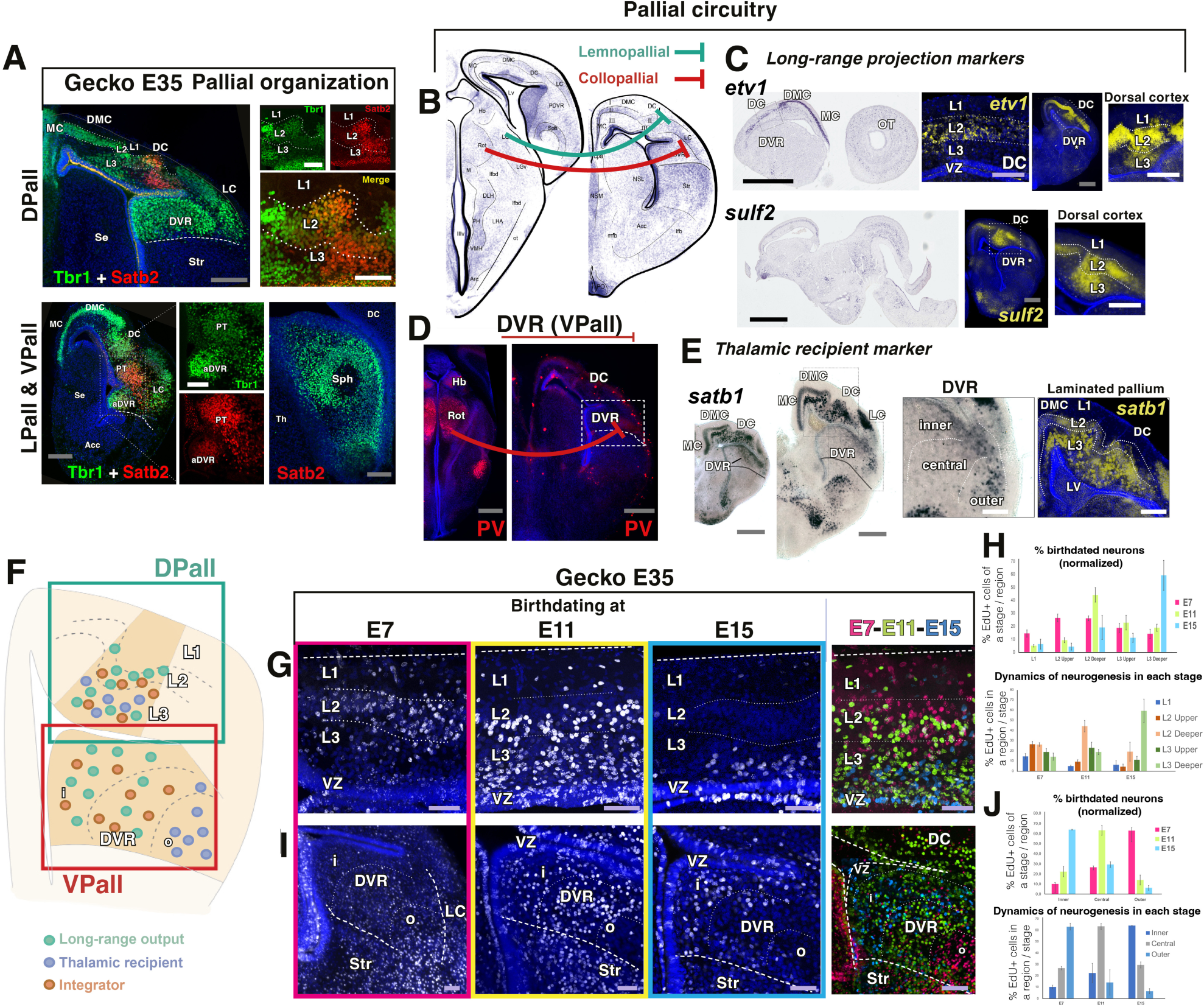
Neuronal birthdating of the gecko pallial sensory circuit. (**A**) Examples of immunostaining for general pallial markers (Tbr1 in green; SatB2 and SatB1 in red) in gecko coronal sections. DMC: dorsomedial cortex; MC: medial cortex; Str: striatum; aDVR: anterior DVR; Sph: spherical nucleus. (**B**) Scheme of two coronal sections of the gecko brain depicting the two main communications of the thalamo-pallial pathway: the lemnothalamic pathway reaching DC (turquoise green) and the collopallial pathway reaching the aDVR (magenta). Abbreviations in **Supplementary Materials**. **(C**) In situ hybridization (ISH) against etv1 and sulf2 mRNAs in the E35 gecko brain, markers of long-range output neurons in mice. Sagittal and coronal sections. Colorimetric staining, pseudotransformed in fluorescent gold staining on the right images. VZ: ventricular zone. (**D**) PV immunostaining in red, two coronal sections at the level of the thalamus (left) and of the telencephalon (right). PV stains rotundal neurons at the thalamus, part of the collothalamic pathway, and their projections reaching the lateral portion of the DVR. (**E**) Colorimetric ISH against satb1, marker of thalamic recipient neurons in mouse cortex, in coronal sections of the E35 gecko telencephalon, pesudotransformed in fluorescent gold at the right image. Satb1 expression is highest at the outer portion of the DVR in the collothalamic pathway, and in the deeper layers of DC (L3 and deep L2). (**F**) Scheme depicting a coronal section of the chick telencephalon with the location of neurons of pallial circuits (**G,H**) Birthdating of the DC neurons at three neurogenic stages. (**G**) Images of the EdU labeling. At the right, pseudoimage made by the merge of images coming from three different animals, showing the distribution of DC cells according to birth time. (**H**) Quantification analysis of the birthdating in DC. (**I,J**) Birthdating of the DVR neurons at three neurogenic stages. (**I**) Images of the EdU labeling. At the right, pseudoimage made by the merge of images coming from three different animals, showing the distribution of DVR cells according to birth time. (**J**) Quantification analysis of the birthdating in DC. DAPI counterstain in blue in A, C, D, E, G and I. Colored scale bars: black, 1 mm; grey, 250 µm; white 100 µm; light purple, 50 µm.

We then performed equivalent birthdating experiments to those in chick, EdU injections at three different time points during the neurogenic period of the gecko pallium (E7 to E15) in order to reveal the temporal construction of the reptilian sensory circuits (**Fig. 3G,J**). The developmental program giving rise to the DVR, in the VPall, followed a clear outside-first inside-last gradient of neurogenesis (**Fig. 3I,J**). Early born (E7) DVR neurons of the gecko located the outer-most region of the DVR, deep beneath the lateral cortex. At intermediate stages (E11), newborn neurons tended to locate the central portions of the DVR. Finally, latest-generated neurons (E15) hardly migrated away from the ventricular surroundings and located in the inner-most DVR region. An equivalent outside-in gradient of neurogenesis also led the formation of the DC, in the DPall, although not as sharp clear as in the DVR (**Fig. 3G,J**). Early-generated neurons (E7) in the dorsal cortex mostly locate the compact, cell-dense layer 2 (L2). At intermediate stages (E11), newborn neurons settle L3 and the deep stratum of L2, suggesting that the earliest neurons locate the upper half of layer 2. The latest generated neurons barely separate away from their original site at the ventricular zone, and locate the deepest stratum of L3.

This sequential neurogenesis verified that, akin to chicks but in stark contrast to the mouse neocortex, the initial neurons generated in the VPall sensory circuit in geckos were thalamic recipient neurons. The neurogenic progression observed in the DVR appears to be conserved in sauropsids, deviating from the pattern observed in mammals. Regarding the DPall sensory circuit, it is probable that geckos and mammals exhibit shared neurogenic characteristics. In both taxa, the long-range output neurons emerge as the first neurons in the circuit, diverging from the pattern observed in any avian pallial circuit. These data support the notion that pallial circuits develop differently across amniotes, revealing an unexpected evolutionary diversity.

### Single RNA sequencing of early-born pallial populations

To confirm that the glutamatergic neurons of these amniote circuits are not conserved homologous cells, we proceeded to compare their transcriptional signatures and trajectories during development. We used our original *BirthSeq* method (*28*) to isolate and analyze birthdated cells days after their generation time. This method allowed us to compare those chick and mouse pallial neurons that were generated at equivalent neurogenic timepoints. By applying BirthSeq, we performed single cell RNA sequencing (scRNAseq) on cells generated early during the formation of the pallial circuits, both in chick (cells born at E4-E6 period, **Fig. 4A**) and in mouse (cells born at E12-E14; **Fig. 4D**). We focused on the earliest-born pallial neurons because these were shown to play divergent roles in the sensory circuit of the two species (**Figs. 1,2**).

**Figure 4.**
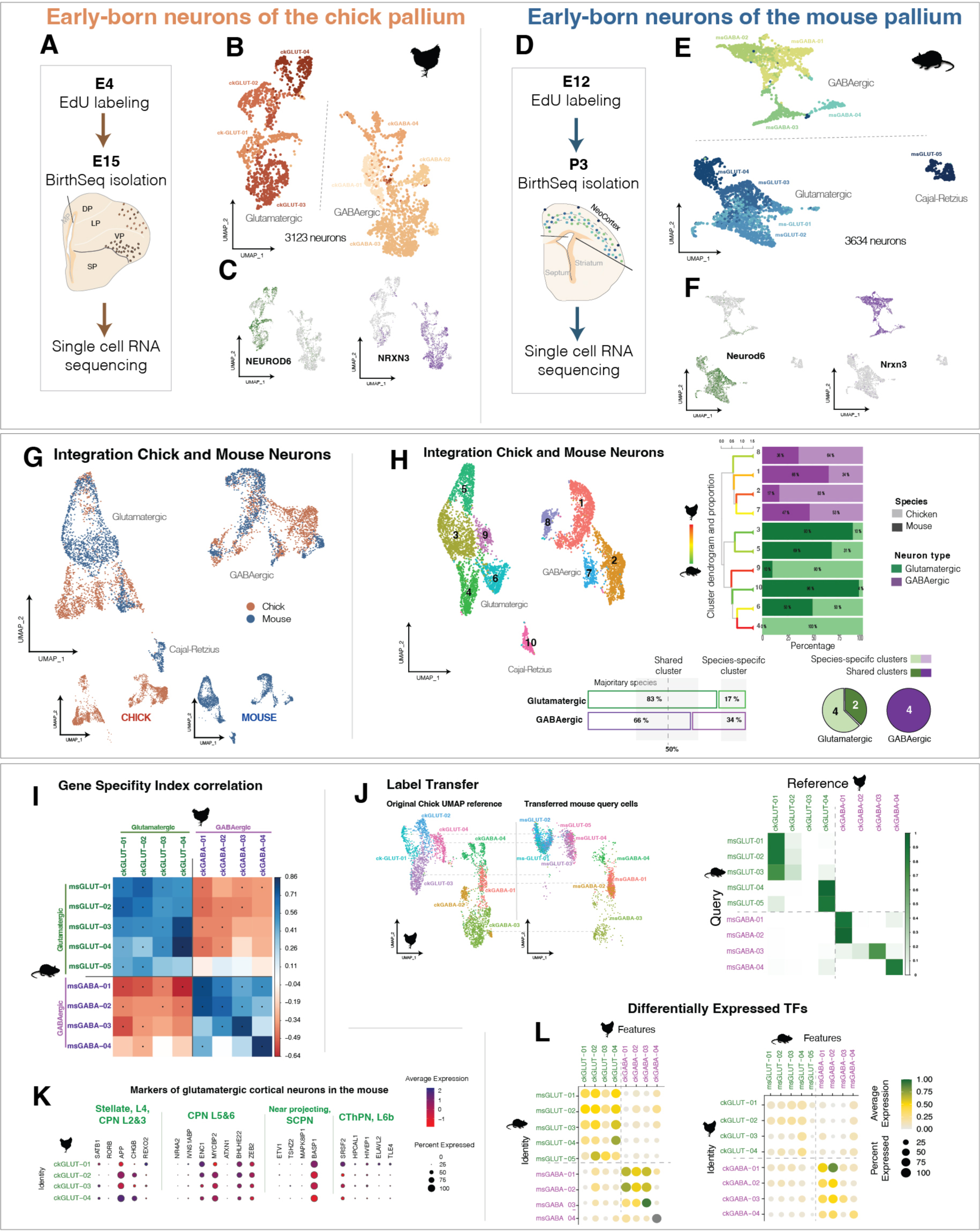
– Transcriptomic comparison of chick and mouse early-born pallial neurons. (**A,F**) Scheme depicting the extraction and analysis of early born neurons in the maturing pallium of chick (**A**) and mouse (**D**). (**B,E**) UMAP graph of the main types of chick (**B**) or mouse (**E**) early-born pallial neurons. (**C,F**) Gene expression UMAP graphs of two key markers for glutamatergic (*NEUROD6*) or GABAergic (*NRXN3*) neurons in chick (**C**) or mouse (**F**). (**G**) UMAP graph of the seurat integration merging the two species datasets. (**H**) Integrated UMAP clustering. At the right, proportion of each species cells to the integrated clusters. Bottom left, average proportion of neurons from each species in either glutamatergic or GABAergic clusters. Bottom right, proportion of clusters considered as species-specific or shared by both species. (**I**) Correlogram of gene specificity indexes between pairs of clusters of the two species. (**J**) Label transfer analysis of mouse cells onto the chick annotated reference UMAP. Right, proportion of cells from a mouse cluster that were assigned to a given chick cluster. (**K**) Gene expression DotPlots of key markers for glutamatergic cortical neurons (*27*) for each chicken glutamatergic cell type (**L**) Percentage of module expression for each cell type (Top4 Marker Genes) on the other species cell types (left, mouse modules; right, chicken modules).

*BirthSeq* provided us two datasets of maturing cells (11 days after EdU administration) enriched in neurons generated early in pallial development (**Figs. 4** and **S11-14+**, as also detailed in (*28*)). Chick pallium dataset comprised 3123 neurons distributed in 8 clusters (**Figs. 4A-C and S13-14**); Mouse pallium dataset comprised 3634 neurons distributed in 9 clusters (**Fig. 4D-F and S11-12**). Each specieś dataset comprised both glutamatergic neurons generated within the pallium and GABAergic interneurons that migrated tangentially from the subpallium (*29–31*). At a first glance, when comparing the differentially expressed genes (DEs) of these neurons, we observed that many of them were not common amongst the glutamatergic clusters of the two species, whereas there were more similarities in the GABAergic clusters (**Figs. 4C,F** and **S11-14 heatmaps**). For instance, mouse glutamatergic clusters (*Neurod6, Slc17a6*) were identified by the expression of *Sox11*, *Id2*, *Pcp4* and *Mef2c* genes (**Fig. S11**), whereas in the case of chick glutamatergic cluster the DE genes were *NRN1*, *NRP2*, *PPP1R17*, *TENM3* or *SATB2* (**Fig. S13**). In the case of GABAergic clusters (*Nrxn3, Gad*), some clusters shared the expression of interneuron subtypes such as *SST*, *ZFHX3*, *ADARB2* or *MEIS2* (**Figs. S12** and **S14**). In order to compare cells from both species, we integrated the two datasets to visually understand cell similarities (**Fig. 4G**). GABAergic interneurons largely overlapped in the integrated UMAP and formed mixed clusters at different resolution values, pointing out to transcriptomic similarities. Glutamatergic neurons, on the other hand, tended to arrange in a more disperse, species-specific organization, with a clear example in Cajal-Retzius cells (msGLUT-05, *Reln*, *Lhx1,* the first cells to be generated in the cortical neuroepithelium (*32*, *33*)), which were specific of the mouse dataset. The integrated dataset clustering showed that, amongst the 6 glutamatergic clusters, 4 were considered species-specific (2 chick-specific, 2 mouse-specific, with over 85% cells coming from the main species), whereas none of the 4 GABAergic clusters comprised such a majority of cells from one single species (**Fig. 4H**). The broader transcriptomic similarity of GABAergic neurons was then confirmed by the correlation of gene specifity indexes (Gene Specificity Index, GSI;(*11*)) (**Fig. 4I**). When comparing the transcription factors (TFs, genes more involved in cellular specification during development), expressed in the two datasets, the largest correlations (up to 0.86) were found among GABAergic cells. Remarkably, transcriptomic correlation was larger between pairs of clusters, suggesting transcriptomic equivalence in a one-to-one cell type fashion. However, only one chick glutamatergic cluster (ckGLUT-04) was found similar to two mouse clusters (msGLUT- 03 and -04). No other cell type-to-cell type equivalence was identified. The general GSI similarities between glutamatergic neurons seemed more related to the transcriptional profile leading the glutamatergic phenotype than to the actual transcriptome involved in specific cell type differentiation. We tested this possibility by comparing the expression of each cluster DE genes, which tend to play roles in the specification of cell types (**Fig. 4L**). We generated modules comprising the most DE genes, from either the whole sequenced transcriptome or only from the transcription factor gene family, and analyzed the average expression of these modules in the cell clusters of the opposite species. In the modules comprising the 5 most differentially expressed TFs for each cluster (**Fig. 4L**) GABAergic cells seemed to be more similar at cell-type level, regardless of the original species used for defining the module. In the case of glutamatergic cells, these shared a common background similarity, especially when mouse clusters were to define the module of expression. ckGLUT-04 cluster (*SATB2*, *MEF2C*, *CCK*, *CRHR2,* **Fig. S13**) was the most similar to a mouse cluster (msGLUT-04, *Mef2c*, *Ptprk*, *Arpp21,* **Fig. S11**) between the glutamatergic clusters. This mouse cluster corresponded to intratelencephalic cortical neurons (*28*), and its cells were likely the latest generated in our experiment. Then, we performed label transfer analysis (*6*) to interrogate each cell from a dataset what cell cluster from the other species dataset were transcriptionally more similar to. This test ultimately confirmed the larger cell type similarity between early born GABAergic neurons respect to early born glutamatergic neurons (**Fig. 4J**). Altogether, these analyses evidenced that early-born glutamatergic neurons showed divergent transcriptomes, as expected from their different developmental origin and circuit function. GABAergic cells, on the other hand, showed higher conservation at transcriptional level.

As a final comparison we asked whether chick early-born glutamatergic neurons expressed the typical molecular signatures of either upper-layer or deeper-layer neurons of the mouse cortex (*27*). We identified that some typical TFs and genes were expressed by one or more chick clusters, but the combination of those TFs typical of a cortical cell type was not found in any chick cluster (**Fig. 4K**). As a conclusion, early-born chick glutamatergic neurons are not only different to mouse early-born glutamatergic neurons at several levels of comparison, but these chick cells are also not similar to either upper- nor deeper-layer cortical neurons. Their transcriptional signature evolved separately from that of both mammalian types.

To understand how early-born glutamatergic neurons diverged in their maturing transcriptome we analyzed the transcriptomic developmental trajectories of these neurons. We birthdated early cells (E4 in chick, E12 in mouse) and applied BirthSeq two days only after EdU administration. We then sequenced birthdated cells comprising early differentiating neuroblasts and progenitors at intermediate pallial neurogenesis (E6 in chick and E14 in mouse) (**Figs. 5 and S15-18**). We performed an equivalent comparison of both species datasets as the one depicted above, including cell cluster identification, integration of datasets, GSI correlation, DE module analysis and label transfer. These experiments brought up three main results. First, that amongst the different cellular types sequenced, the largest similarities were found amongst the neural stem cells of both species. Specifically, pallial radial glial cells (RGCs) were strongly similar at transcriptomic level (**Fig. 5**). Interestingly, we did identify a population of mouse intermediate precursor cells (IPCs) enriched in *Eomes* expression, but its identification in chick was elusive (*19*). No transcriptomically equivalent population of IPCs was found in the chick dataset as shown by the lack of merged IPC cells in the integration of the two datasets (**Fig. 5G**). But when the integration was performed without the usual cell cycle correction (**Fig. S17**) the ckGLUT-01e cluster showed a fraction of cells with some mouse IPC features. Also, there were abventricular division in a germinative SVZ of the developing chick pallium (**Fig. S17G**). Therefore, intermediate precursors of novel transcriptomic nature did exist in the chick pallium, but is relevance over pallial neurogenesis seemed restricted by their low numbers. Secondly, GABAergic cells showed the largest similarities between specific cell types also during their differentiation progress (**Fig. 5**). This implied that the transcriptomic divergence was determined from very early on in their development. And finally, we observed that some transcriptomic dissimilarities between glutamatergic cells were early determined from their neuronal birthdate. Glutamatergic cells from both species showed transcriptomic similarities related to the differentiation of main glutamatergic type (*TBR1*, *SLC17A6*, *NEUROD* genes), and a minor equivalence of specific cell types. As described above, these divergences were more pronounced between the maturing neuron datasets.

**Figure 5.**
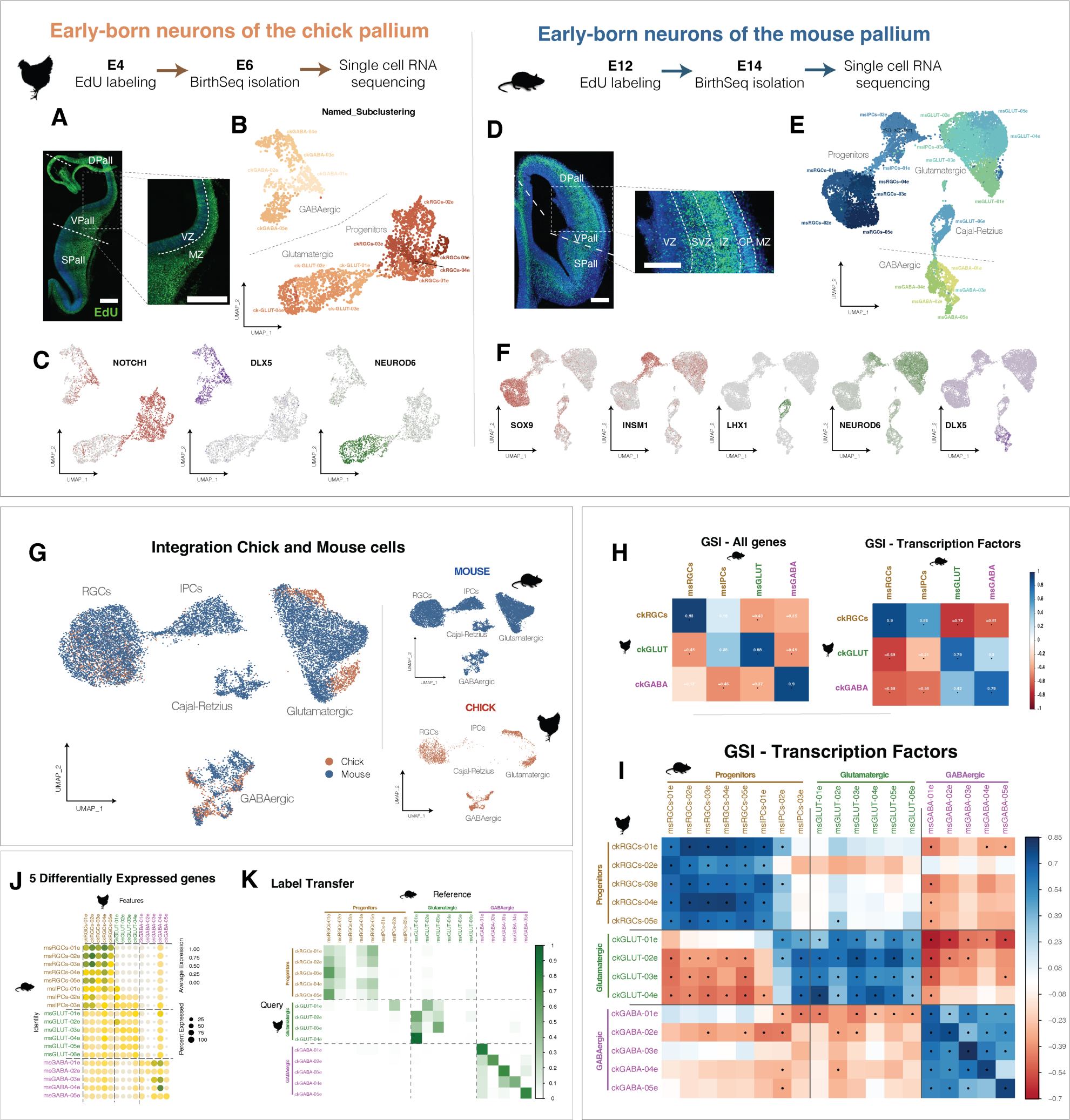
– Transcriptomic comparison of early-born neurons and progenitors between chick and mouse. (**A-F**) BirthSeq isolation of early born populations two days after Edu labelling in chick (**A-C**) and mouse (**D-F**). (**A,D**) EdU staining of chick and mouse pallia at the moment of cell isolation. (**B,E**) UMAP graph of the main types of chick (**B**) or mouse (**E**) early-born pallial populations. (**C,F**) Gene expression UMAP graphs of several key markers for RGCs (*NOTCH1*), IPCs (*INSM1*), glutamatergic neurons (*NEUROD6*), Cajal-Retzius cells (*LHX1*) or GABAergic (*DLX5*) neurons in chick (**C**) or mouse (**F**). (**G**) UMAP graph of the seurat integration merging the two species datasets. (**H**) Correlogram of gene specificity indexes between pairs of main cell types of the two species. (**I**) Correlograms of gene specificity indexes between pairs of clusters of the two species, considering transcription factors only in the comparison. (**J**) Percentage of module expression for each cell type (Top5 Marker Genes) on the other species cell types. (**K**) Label transfer analysis of chick cells onto the mouse annotated referenc; proportion of cells from a chick cluster that were assigned to a given mouse cluster. DAPI counterstain in blue in A, D. Scale bars, 100 µm.

Our scRNA-seq analysis of early-born neurons of the pallium showed substantial disparities between the transcriptomic programs of GABAergic respect to glutamatergic neurons of both species. These programs were mainly divergent on the glutamatergic populations, whereas deeply conserved between GABAergic neurons.

### Distribution of early-born chick pallial neurons

The location of specific cell types within the chick pallium remains largely unknown. To investigate and describe the distribution of these distinct cell types, we conducted *in situ* sequencing (ISS) for genes identifying early-born neuronal clusters (**Figs. 5, S19-S21**). Using 84 genes selected from the scRNAseq dataset we examined their expression at a cellular level in coronal sections of three E15 chick brains. Many of these genes were expressed exclusively in different telencephalic sectors, enabling the identification of various pallial territories and circuit stations (**Figs.6A, S20**).

A total of 911495 cells were sequenced and analyzed. By comparing ISS reads to clusters identified through scRNAseq, we detected 660117 neurons in the tissue that likely corresponded to the scRNAseq clusters (**Fig. 6B**). This correspondence was determined by the expression levels of significant genes in the ISS data (**Fig. 6B,C**). Glutamatergic neurons were found to be distributed in four clusters across different pallial regions, exhibiting distinct gene expression patterns and functions within the pallial circuitry (**Fig. 6D,E, S19**). GABAergic interneurons also displayed differential distribution among four neuronal clusters, with some spanning the entire pallium and others being more restricted to specific regions (**Fig. 6D,F**).

**Figure 6.**
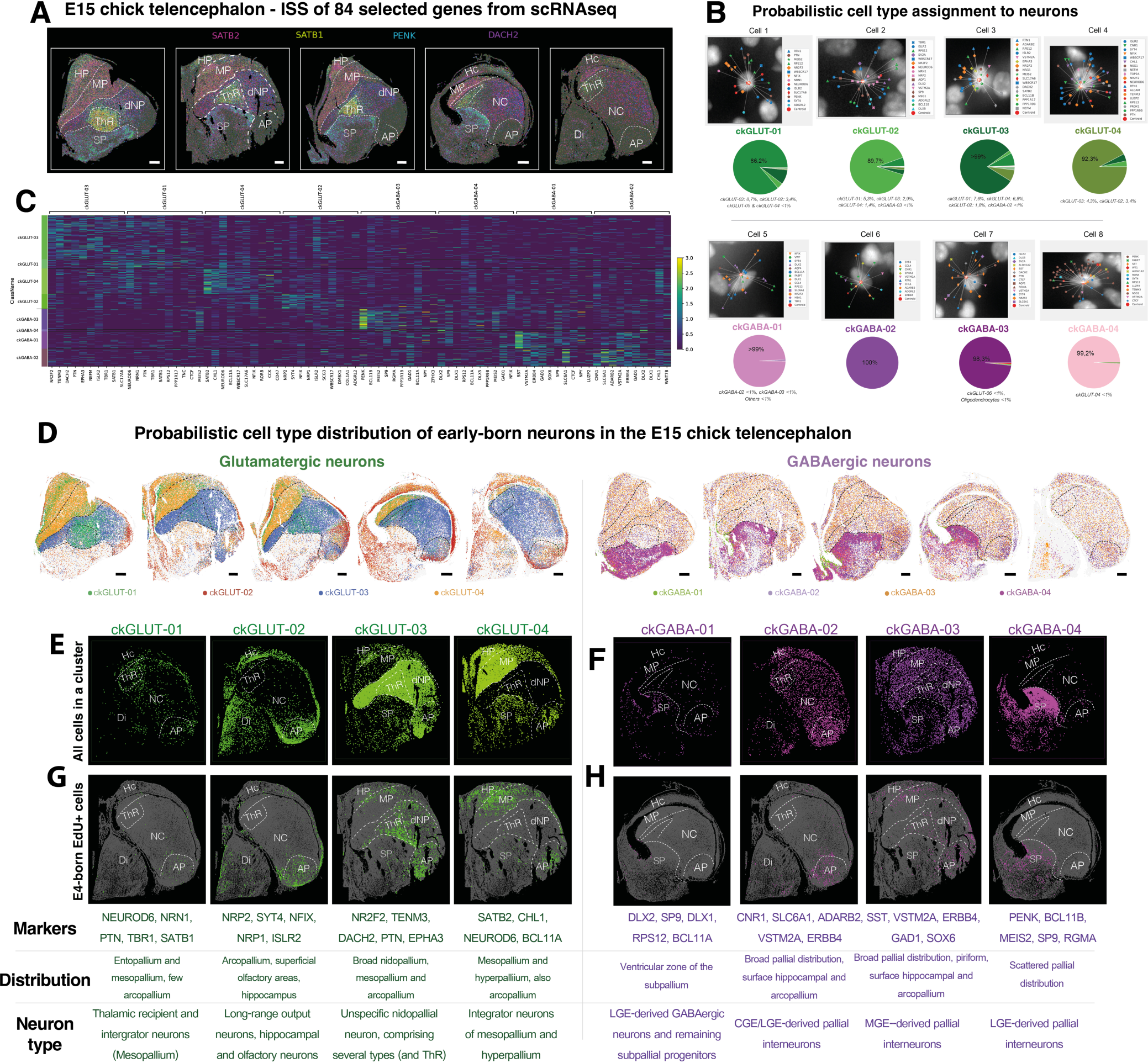
– Identification of early-born neurons of the chick pallium by *in situ sequencing*. (**A**) Expression maps of 4 informative markers, chosen from a panel of 84 ISS genes. (**B**) Examples of in-situ probabilistic cell type assignment for the neuronal clusters: for each cell (identified using DAPI signal), a map of its associated transcripts is built, taking into account the probability of gene co-expression derived from scRNAseq data (images). Based on its expression profile, each cell is then assigned a probability of belonging to each one of the scRNAseq clusters (pie charts). (**C**) Heatmap comparison of the gene expression profiles, as detected by informative genes in ISS, of the different cell clusters. (**D**) Anatomical distribution of the glutamatergic and GABAergic neuronal sub-populations. Left panel shows the distribution of glutamatergic neuron types. (**E-H**) Joint birthdating and transcriptional profiling of selected cell clusters: Glutamatergic (**E**) and GABAergic (**F**) neuronal subclasses distributions are shown on two of the samples, together with a description of their characteristic markers, anatomical distributions and functional annotations (table). Of each neuron subclass, the EdU-positive cells can be extracted and plotted separately (**G-H**), allowing to analyze the distribution, across neuron types, of the birthdated cells. This allows to visually inspect the relationship between birth date and cell identity. Scale bars in A and D, 500 µm. Sections were obtained from 3 different embryos.

The distribution identified through this probabilistic cell typing reflects our scRNAseq clusters correspond to major types of developmental neurons, but not all those neurons were early-generated.

To identify the early-born neurons among these clusters, we employed Neurogenes-iss, a combination of ISS and EdU birthdating (*28*). A fraction of 12.3% of the identified neurons within these clusters were EdU+, indicating their generation shortly after E4 EdU administration. The proportion varied across clusters (from 17.2%: ckGLUT-02 to 5%: ckGABA-04). Neurogenes-iss facilitated the precise localization of early-born neurons, along with their transcriptomic identity (**Fig. 6G,H and S19,S21**). The distribution of early-born neurons ckGLUT-01 to ckGLUT-03 aligned with expectations within the circuit (EPall, APall), as well as other pallial territories, such as the hippocampal and parahippocampal areas, nidopallium caudale, and mesopallium (**Fig. S20**). Remarkably, early-born neurons belonging to cluster ckGLUT-04, which exhibited similarity to msGLUT-04 (**Figs. 4, S18**), predominantly corresponded to mesopallial cells, and were therefore located outside the canonical circuit (*4*, *34*). They were found in the mesopallium and hyperpallium, with some sparsely distributed in parahippocampal areas and APall.

Early-born GABAergic neurons exhibited a preference for subpallial territories but were also widely distributed throughout the pallial territories (**Fig. S21**). Notably, early-born GABAergic PVALB+ neurons in the EPall, primarily generated at E4 in chick (**Fig. 1** and **S03**), were absent in our ISS analysis, possibly due to their absence in our initial scRNAseq. Overall, our novel analysis of early-born neurons’ transcriptomics and distribution in the pallium confirmed their expected presence in various pallial regions. Glutamatergic neurons identified within the circuit regions corresponded to thalamic recipient neurons and long-range output neurons, while those outside the circuit belonged to integrator classes in other pallial territories, including the hippocampal and olfactory areas. These findings demonstrate that early-born neurons in the chick pallium differ from their counterparts in the mouse pallium at developmental, transcriptomic and distribution levels. These divergent neurons are generated from homologous radial glial cells, which follow divergent developmental programs to generate different neurons in mouse and chick. And ultimately, neurons assemble into non-equivalent stations of the pallial circuitry.

### Functional constraints limit circuit architecture

While developed differently, mouse and chick pallial circuits supersede sensory processing alike (*35*). We resort to mathematical modeling to elucidate the possible principles for such functional convergence. Each pallial circuit consists of a population of neurons functionally connected prominently by volume transmission and electrical coupling (*36*). Thus, there is no preferential connection between one cell and another: cell connections within individual pallial circuits and across are normally distributed around zero (no coupling) by their electrical field fluctuations propagating by the extracellular milieu. Thus, the whole pallial circuit can be modeled by a block random matrix **J**, where connections within pallial areas will generally have different standard deviations (g_r) from those across regions (g_x) due to their different physical (spatial) constraints (Supplementary notes). The theory of random matrices then tells us that the activity of pallial circuits is dominated by the leading eigenvalue Λ_1_ of the matrix reflecting the block structure of such deviations (*37*).

We “grow” pallial circuits by progressively increasing the size of our matrix blocks, adding neurons at fixed time stamps based on the neurogenesis probability inferred from our data (**Fig. 7A,B**). The functional readout of this process is the change in the eigenvalue spectrum of the pallial connectivity matrix **J** (**Fig. 7C,D**), which is contained in a circle of radius Λ_1_. Then, the appearance of spontaneous ongoing activity emerging in late embryonic stages and persisting after birth is reflected in our model by Λ_1_ values growing to one and beyond. This can be achieved by constraining the strength of neuronal connections within vs. between regions (*38*). In particular, neither of the connection types alone can account for such emergence, but it is necessary for both connections to be sufficiently strong for the circular spectrum to grow beyond the unitary circle (**Fig. 7E**). Intriguingly, despite the divergent neurogenic programs and structural differences, similar values of coupling strength can be found in both mice and chicks, resulting in similar circuit activity as reflected by similar Λ_1_ curves above one towards approaching birth. This, in turn, suggests that neurogenesis programs in the avian vs. mammalian brain may have converged by evolutionary constraints for adaptation to equivalent functions.

**Figure 7.**
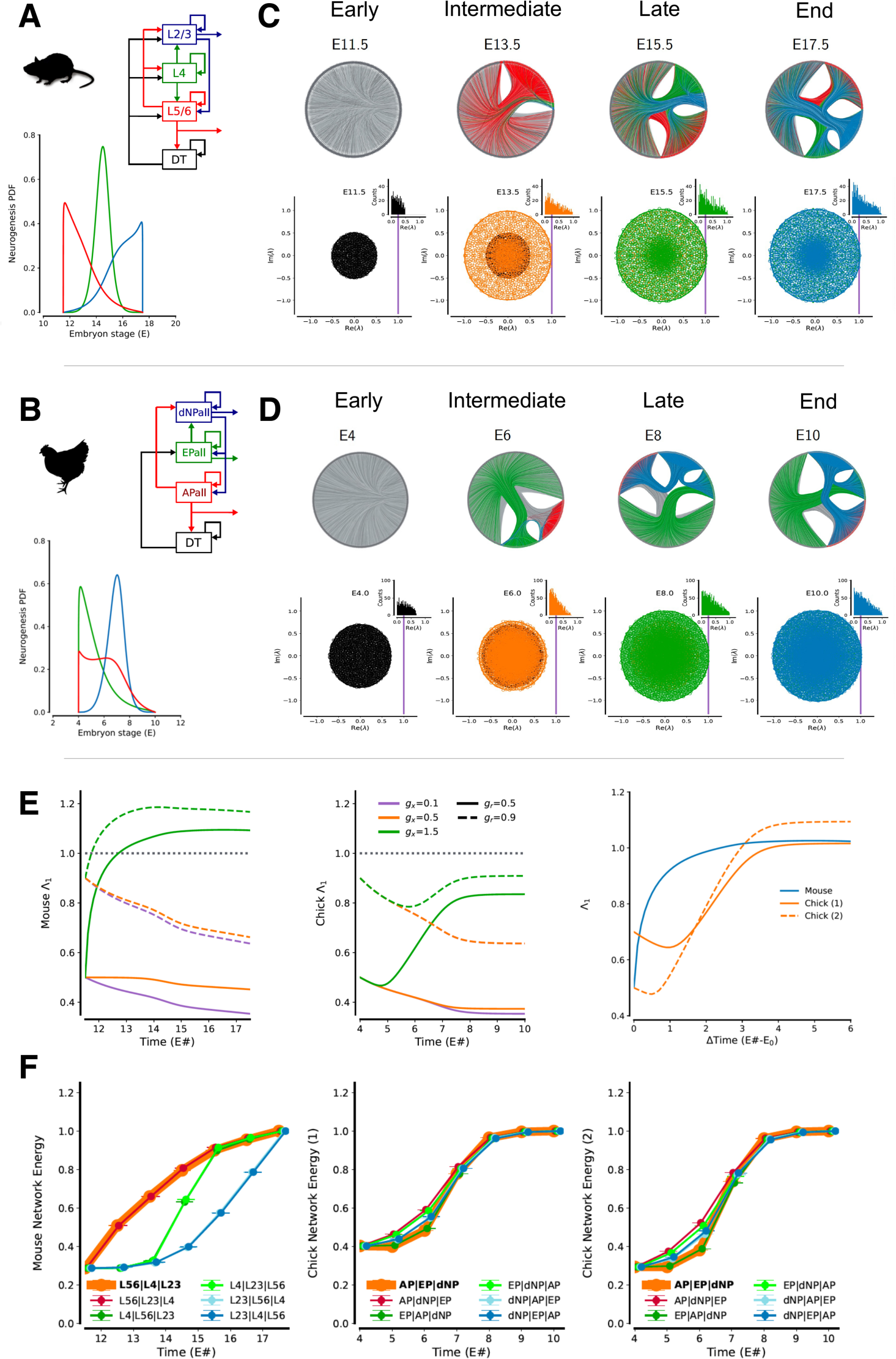
– Mathematical modelling supports convergence by functional constraints. (**A**,**B**) Mouse and chick pallial circuits and associated neurogenesis probabilities (PDF) reconstructed from available data (see **Supplementary Notes S1**). (**C**,**D**) Simulation of neurogenesis by growth of random networks with structured connectivity (top graphs, only 60% connections shown). (**E**) Emergence of spontaneous activity leads to network eigenvalues exceeding one for an appropriate choice of intra-vs. inter-region connection strength. Spectral graph energy is maximal in the mouse and chick for the documented neurogenic program, but not for any other random permutation of the temporal sequence of the different neurogenic niches.

To test the latter hypothesis, we look at the propensity of the circuit to be active in the function of neurogenesis, estimating the spectral energy of the graph associated with the matrix **J**, that is, the sum of all eigenvalues (in absolute value) of **J** (*39*). As we permute the three neurogenic programs across the pallial/cortical regions, we prove that avian and mammalian circuits have evolved to maximize energy independently with respect to neuronal coupling strength (**Fig. 7F**). Given that activity promotes neuro and synaptogenesis, and vice versa (*40*), maximizing the energy function supports the notion that the evolutionary divergence of avian and mammalian circuits is necessary to optimize their genesis. Both avian and mammalian circuits evolved by functional channeling into equivalent circuits. These circuits converged to respond to the same functional requirements

## Discussion

For decades, there has been a heated debate revolving around identifying homologies between the mammalian-specific neocortex and that of other vertebrates. Various claims have emerged regarding these homologies, based either on gene expression patterns observed in embryonic brains or on neuronal connectivity patterns found in adult brains. Here we tackled the debate from alternative views, researching the developmental formation of the pallial circuitry from neurogenic, transcriptional and mathematical perspectives. We uncover that the developmental rules for the construction of these circuits are different in each species and for each pallial region (**Fig. S22**). The neurons of the three stations of the circuit are generated in different developmental times and brain regions. Amniote species also differ in the progenitors required to generate the circuit, as evidenced by the difference in the pallial sources, and the nature and relevance of IPCs. Moreover, by means of single cell RNA sequencing we reveal that equivalent pallial neurons (those generated in homologous brain regions and equivalent neurogenic times) mature into divergent neuronal types, confirming that the formation of the circuits follows different developmental and evolutionary paths in sauropsids and mammals. Finally, mathematical modelling shows evidence that the sensory circuits of birds and mammals are sculpted by the same functional constraints. Altogether, we demonstrate that the circuits in charge of high-order sensory processing have evolved separately in different vertebrate taxa to converge into a functionally similar circuit.

In our comparison, we operate under the assumption that traits which are conserved must also exhibit equivalence in their developmental programs. Homology, in this context, implies the sharing of a feature that originated from a common ancestor (as discussed in (*19*)). Our assumption is grounded in the belief that the last common ancestor of amniotes already possessed a pallium (*41*). If the circuits of present-day species were indeed conserved and homologous, then they should have inherited the same developmental program that was already present in the pallium of the last common ancestor. This assumption has proven to be valid when applied to other distinct brain territories. Recently, through the utilization of the same birthdating techniques across multiple amniote species, we have discovered that the developmental sequence responsible for generating cerebellar circuitry is profoundly conserved (*42*). Purkinje cells, GABAergic interneurons and granule cells of the cerebellar cortex, all are generated following the same temporal sequence in mouse, chick and gecko. In addition, cerebellar neurons are deeply conserved at transcriptomic level (*43*, *44*). These similarities prove that the evolutionary conservation of a brain structure is reflected, and promoted, by its developmental conservation. It is this developmental conservation which causes evolutionary conservation. The fact that the pallium develops its circuitry in several different ways strongly supports a non-homologous character of the amniote pallial circuits.

Another crucial factor indicating the divergent nature of these circuits in amniotes is the significant differences in their progenitor cells. These differences occur at anatomical level, but also at progenitor cell type level. In mammals, the neocortex is generated in the dorsal pallium from multipotent radial glial cells (*45–47*). However, in birds, the proposed homologous circuit develops in the ventral pallium, which is a distinct pallial territory that predominantly gives rise to the amygdala and olfactory structures in mammals (*30*, *48*). Progenitors in the ventral pallium exhibit transcriptomic differences from those in the dorsal pallium, reside in distant regions of the pallium, and are not evolutionary equivalents (*16*).

Furthermore, neural stem cells in charge of generating the neurons of the circuits differ at their neurogenic potential. In birds, no single progenitor can generate all the neurons of the DVR circuit. Through electroporations that reveal whole-cell lineage and neuronal migration in chicks, it has been demonstrated that progenitors in the VPall can generate neurons for either the EPall and dNPall, *or* the APall (*30*, *49*). Therefore, at least two differentially located progenitors are necessary to assemble a functional circuit. This is a crucial difference to mammals, on which a single radial glial cell is capable of generating each and all the different types of neurons of the cortical circuit (*45–47*). But also, it is important to note that at intermediate stages of pallial neurogenesis in chicks (around E6), only a small proportion of pallial division happens in the SVZ, and its constituent progenitors are not transcriptomically similar to mouse IPCs despite *EOMES* expression. At the equivalent neurogenic timepoint in mouse cortical neurogenesis (E14 in mouse) intermediate progenitor cells (IPCs) reach their peak population (*50*, *51*). These facts point to differences in the direct and indirect neurogenic pathways, a feature previously referred as an evolutionary difference between sauropsids and mammals (*52*). Hence, the cellular trajectory of early born pallial neurons in birds differs from that of mammals at all the examined levels, which led to transcriptomic differences in these early-born populations. It is likely, though, that IPCs play a more relevant role in avian pallial neurogenesis later in development, as shown in a parallel article after a complete description of the developing cell types in the chick pallium (*53*).

Altogether, our data show an early transcriptomic divergence in the glutamatergic cell types, which differ between chick and mouse as early as two days after their generation. This divergence of glutamatergic profiles has been further confirmed by two independent articles, which employ different methodologies in developing, post-hatching and adult chicks. Zaremba et al. (*53*) performed a thorough atlas of the transcriptomic profiles of the pallium of chick, and Hecker and Kempynck et al. (*54*) used a novel deep learning analysis to characterize cell-type specific enhancer codes. In both cases, when compared amongst amniote species, glutamatergic cells were the most divergent cell types in the pallium. And amongst those neurons displaying similarities are mesopallial cells, a shared result in all three studies.

On the contrary to glutamatergic cells, our data, as well as those described in recent articles (*53*, *54*), point to a strong conservation of GABAergic pallial neurons. The developmental formation of interneurons follows equivalent sequences of neurogenesis. And transcriptomic analyses also show similar molecular fingerprints of these GABAergic populations. The conservation of GABAergic neurons of the pallium has been now shown equal for many different taxa. This conservation was, as stated before, demonstrated by similarities in developmental lineage and transcriptomic profiling. The vertebrate groups tested by transcriptomic comparison have been turtles (*11*), lizards (*55*), amphibians (*56*, *57*), and the distant group of cyclostomes -lampreys-which are jawless vertebrates (*58*). The results on all these species were the same: homologous populations of GABAergic interneurons are generated in the subpallium, migrate tangentially into the pallium, differentiate into interneuron subtypes of equivalent transcriptomic profile and contribute to the function of pallial circuits. The molecular reasons behind this dual evolution of pallial circuits (diversified glutamatergic cells vs. conserved GABAergic cells) remain unknown.

The functional similarities between these circuits are explained by parallel convergent evolution, which guided the formation of the circuits under efficiency restrictions. Many species evolve in the same environment, under the same physical rules, and are driven by the same optimization constraints. This implies that functionality limits diversity, at all biological levels. For circuit evolution as well, because circuits may act in similar ways in different species: the shared function of independent circuits limits the diversity of them. We speculate that, following this argument, pallial circuits are more efficient and optimal when arranged in this known fashion (*14*). Surely, evolution tinkered with their development, structure and function. But the surviving circuits, those selected after millions of years of evolution, display limited diversity because evolved under the same optimization criteria (*35*). Convergent evolution sculpted the formation of sensory circuits in amniote species.

## Materials and methods

### Animals

All animal experiments were approved by a local ethical review committee and conducted in accordance with personal and project licenses in compliance with the current normative standards of the European Union (Directive 2010/63/EU) and the Spanish Government (Royal Decrees 1201/2005 and 53/2013, Law 32/107).

#### Chick

Fertilized chick eggs (*Gallus gallus*) were purchased from Granja Santa Isabel and incubated at 37.5 °C in humidified atmosphere until required developmental stage. The day when eggs were incubated was considered embryonic day (E)0.

#### Mouse

Adult C57BL/6 mice (*Mus musculus*) were obtained from a mice breeding colony at Achucarro Basque Center for Neuroscience (Spain). They were housed in a 12/12-hour light/dark cycle (8 AM, lights on) and provided with ad libitum food and water. The day when the vaginal plug was detected was referred to as E0.

#### Gecko

Fertilized ground Madagascar gecko eggs (*Paroedura pictus*) were obtained from a breeding colony at Achucarro Basque Center for Neuroscience (Spain). They were housed in a 12/12-hour light/dark cycle (8 AM, lights on, 27°C/ 8 PM light off, 22°C) and provided with ad libitum food (live crickets) and water. Gecko eggs were incubated at 28 °C in a low humidified atmosphere until required developmental stage. The day when eggs were incubated was considered E0.

### Birthdating of neural populations

Manipulation of chick embryos was performed as previously described (*30*). For gecko eggs, the procedure was equivalent but with adjustments to the smaller size of the eggs and the higher fragility of the eggshell (*42*). Briefly, eggs were incubated in a vertical position at either 37.5 or 28 ° C. 5-ethynyl-2-de-oxyuridine (EdU; ThermoFisher) was provided via intracardiac intravenous injection for chick embryos (E4, E6 or E8) or via intraventricular injection for gecko embryos (E7, E9, E11, E12, E15, E18; all these gecko stages were considered in the comparison by only E7, E11 and E15 were quantified), using a fine pulled-glass needle. Drops of sterilized HBSS were added to the egg and chick and gecko embryos were incubated until E15 or E32, respectively.

For birthdating of embryonic mice, pregnant dams were intraperitoneally injected with EdU 50 mg/kg body weight in sterile saline (BD Biosciences).

### Brain tissue preparation

#### For immunohistochemistry and birthdating analysis

Embryonic murine (E14), all gecko brains, and the chick embryonic brains up to E9, were fixed by immersion in 4% paraformaldehyde (PFA, diluted in phosphate buffered saline 0.1M– PBS, pH 7.3). P3 postnatal mouse pups and E15 chick embryos were anesthetized by hypothermia after immersion of either pup or the chick egg in ice. Then they were transcardially perfused with PBS followed by PFA. All brains were transferred to PBS 6h after fixation, and were sectioned in the coronal plane at 50-70 μm thickness in a vibrating microtome (Leica VT1000S).

#### For in situ hybridization

Gecko embryos at E50-55 stage were anesthetized by hypothermia after immersion of the gecko egg in ice. Gecko embryonic brains were fixed by immersion in RNAse-free 4% PFA. Brains were transferred to RNAse-free PBS 24h after fixation. All brains were sectioned in either the coronal or sagittal planes at 50-70 μm thickness in a vibrating microtome (Leica VT1000S).

#### For in situ sequencing

Animals were anesthetized by hypothermia after immersion of either the mouse pup or the chick egg in ice. Postnatal murine (P3) and late chick embryonic brains (E15) were cryopreserved in 30% sucrose, fresh frozen in optimal cutting temperature (OCT) media and stored at -80°C until sectioning. Tissue was sectioned in the coronal plane at 12-16 µm thickness with a cryostat (Leica CM1950) and collected on SuperFrost Plus microscope slides. Slides were stored at -80°C until processing.

### Birthdating analysis

#### Immunohistochemistry and EdU labeling

Single and double immunohistochemical reactions were performed as described previously (*59*) using the following primary antibodies: mouse antibody to parvalbumin (Sigma, P3088; 1:750), mouse antibody to SatB2 (Abcam, ab51502: 1:750; this antibody recognizes both SatB2 and SatB1 expressing neurons), mouse antibody to somatostatin (Santa Cruz Biotechnology, sc74556; 1:750), rabbit antibody to calbindin (Swant, CB D-28k; 1:1000), rabbit antibody to GABA (Genetex, GTX125988, 1:750), rabbit antibody to Tbr1 (1:2500; kind gift by Prof. Hevner, Univ. of California, San Diego).

For secondary antibodies (all 1:1000), we used Alexa 568 goat antibody to rabbit IgG (Molecular Probes, A11011), Alexa 488 goat antibody to rabbit IgG (Molecular Probes, A11034), Alexa 488 goat antibody to mouse IgG (Molecular Probes, A11001) and Alexa 568 antibody to mouse IgG (Molecular Probes, A11004).

At the end of the immunohistochemistry process, EdU molecule was detected in the sections following the *Click* reaction, with the commercial EdU *Click-It kit*, and revealed with Sulfo-Cyanin5 fluorophore (Lumiprobe).

All sections were counterstained with DAPI.

#### Image capture

All fluorescence immunostaining images were collected using a Leica SP8 laser scanning confocal microscope (Leica, Wetzlar, Germany), a Leica Stellaris 5 scanning confocal microscope (Leica, Wetzlar, Germany), a Zeiss LSM 710 confocal microscope (Carl Zeiss Microimaging, Germany) or a 3DHistech Panoramic Midi II digital slidescanner (3DHistech, Hungary).

Images obtained with Leica confocal microscopes were acquired with the LAS X software, using a 20X air objective and a z-step of 3 μm. The signal from fluorophores was collected sequentially-green light and far-red light were collected together. Same image parameters (laser power, gain and wavelengths) were maintained for images from each slide and adjusted for new animals. For big brain sections, tile-scan images were composed. All images shown are projections from z-stacks ranging from 10 to 35 μm of thickness, typically 20 μm.

Images obtained with the Zeiss confocal were acquired with a 20X air objective. Same image parameters (laser power, gain and wavelengths) were maintained for images from the same brain and adjusted for new slides. For big brain sections, tile-scans were composed.

Images obtained with the slidescanner were acquired in single layer mode, using the 20X objective and collecting the signal from fluorophores sequentially.

#### Quantitative analysis of cell populations

Quantitative analysis of cell populations in vivo was performed in images taken from a minimum of 4 successfully birthdated animals (specific numbers for each experimental paradigms are included in figure legends). For cell densities, quantifications were estimated maintaining the same z-stack size between conditions. A minimum of three random sections were quantified per animal, attempting to maintain a complete rostro-caudal representation of all the pallial areas of interest.

For the DVR birthdating experiments, one field per section and region was analyzed. Individual images were imported to ImageJ, and specific regions of interest (ROIs) delimiting APall, EPall and dNPall were created. Next, in each region, cells were quantified on a single confocal plane of a 512 x 512 μm^2^ field. Values were then normalized to the quantification area.

In the hyperpallial birthdating analysis, two fields per section and region were analyzed in order to check if there were inner/outer lateromedial gradients of cell generation. Individual images were imported to LasX, and specific ROIs delimiting HA, IHA and HI/HD were created. Next, cells were quantified in (1) a delimited area of the most inner part of the hyperpallial regions, and (2) a delimited area of the most outer part. Quantified cells were then normalized to the quantification area.

#### Statistical analysis

SigmaPlot v13.0 (San Jose, CA, USA) was used for statistical analysis. Data normality and homoscedasticity were analyzed for each group, by Shapiro-Wilk and Brown-Forsythe tests. When ANOVA assumptions (normality and homoscedasticity) were not accomplished, a logarithmic transformation of the data was performed.

For analysis of pairs of groups, the Student’s t-test was performed on parametric data and the U-Mann Whitney test on data that was non-parametric after logarithmic transformations. For analysis of three groups, one-way ANOVA test was performed to determine the overall effect of each factor, resorting to Kruskal-Wallis test when data were no parametric after logarithmic transformations.

In all cases, all pairwise multiple comparisons (Holm-Sidak method or Tukey test with Dunńs method) were used as a post hoc test to determine the significance between groups in each factor. Only p<0.05 was considered as statistically significant (*p<0.05; **p<0.01; ***p<0.001). Results were presented as mean ± standard error mean (SEM). The number of independent experiments is shown in the respective sections.

All the statistical analysis are summarized in **Supplementary Table S1**.

### In situ hybridization

M13forward (5′GCCAGGGTTTTCCCAGTCAC3′) and M13reverse (5′GGAAACAGCTATGACCATG3′) primers were used to obtain the fragments from our cloned genes that were used to get riboprobes. Sense and antisense digoxygenin-11-UTP-labeled (Roche) riboprobes were synthesized according to the detailed procedures previously described (*60*). All the steps and procedures related with the extraction of brain samples, tissue processing and in situ hybridization in cryostat and floating sections were done as previously described (*61*).

*etv1*, *sulf2*, *rorb*, *dbp* and *satb1* cDNA fragments, obtained by RT-PCR, were cloned into TA vectors to later synthesize the RNA probes as previously described (*62*). The cDNA was used as a template for PCR reactions using Taq polymerase (Promega, Cat. M8305) and specific primers (**Supplementary Table S2**). The PCR products were then cloned into the pGEM-T Easy Vector (Promega, Cat. A1360, Spain) and sequenced (ACTI, University of Murcia).

### Single cell RNA sequencing (scRNAseq)

#### Cell preparation - Birth-Seq

Isolation of birthdated cells for scRNAseq was performed as previously described (*28*). Briefly, chick and mouse embryos were birthdated at early neurogenic timepoints (E4 or E12, respectively) and sacrificed either two or eleven days after EdU injection. P3 postnatal mouse pups and E15 chick embryos were anesthetized by hypothermia after immersion of either pup or the chick egg in ice.

Brains were extracted in ice-cold HBSS. Then, the pallial region was microdissected under a stereomicroscope, collected in ice-cold HBSS, pooled and cut into small pieces. Tissue was dissociated in a species and stage specific manner, according to **Supplementary Table S3**. Following enzymatic digestion, chick (both E6 and E15) and mouse P3 tissue was manually homogenized by carefully pipetting and tissue clogs were removed by filtering the cell suspension through a 40 µm nylon strainer to a 15 ml Falcon tube containing 4 ml of 25% fetal bovine serum (FBS) in HBSS. After mouse E14 tissue digestion, FBS was added to the mix, cells were manually dissociated by pipetting and tissue clogs were removed by filtering the cell suspension though a 40 µm nylon strainer.

Dissociated cells were then centrifuged at 200g for 10 min at room temperature (RT) and EdU molecule was detected in the pelleted cells. Following reaction, cells were washed thrice by centrifugation at 200g for 10 min and suspended in 1 ml of HBSS then passed on a 50 µm cell strainer. PI-/EdU+ cells, gated to include only the top 10% brightest cells, were finally FAC-sorted: for the E6 chick E6 a Synergy 3200 2L sorter (Sony) was used; for the E15 chick sample a FACSAria (BD Biosciences) was used; and for both mice samples a FACS Jazz (BD Biosciences) was used.

FAC-sorted cells were centrifugated at 200g for 10 min and resuspended in HBSS, at a concentration of 1200 cells per µl. Cell concentration and viability was verified using a TC20 Automated Cell Counter (BioRad). Next, cells were processed for single cell GEM formation following the standard Chromium Single Cell 3’ v3.1. Briefly cells were placed at the Chromium chip with the beads and reagents for the RNA capture and cDNA amplification. After the emulsion were obtained, the cDNA was amplificated in a thermocycler. Post amplificated cDNA, now marked with individual cell barcode, were fragmented, ligated the sequencing adapters and amplified. After that, the PCR product were purified and quantified. 24,000 cells per reaction were loaded aiming a targeted cell recovery of 12,000 cells (with 50% efficiency). Single cell library was prepared using “Chromium Next GEM Chip G Single Cell Kit”, 16 rxns PN-1000127, “Chromium Next GEM Single Cell 3ʹ Kit v3.1”, 4 rxns PN-1000269 and “Dual Index Kit TT Set A”, 96 rxns PN-1000215, following “Chromium Next GEM Single Cell 3’ Reagent Kits v3.1(Dual Index) User Guide” (Document number CG000315). Resulting libraries were sequenced on a Novaseq 6000 for an approximated 50.000 reads per cell.

#### Single cell pre-processing

For both species, 10X *CellRanger* v6.0.2 was employed for alignment and demultiplexing of *FASTQ* files to obtain feature-barcode matrices. The genomes used as reference for each species alignment were mm10-2020-A for mouse and Gallus_gallus6_98 for chicken. The quality control statistics related to this alignment steps and others are summarized in **Supplementary Table S4**. Afterwards, data matrices were imported to *R* (v4.1.0), where *Seurat* (v4.1.0) (*63*) was employed to further analysis, as describe in their vignettes (https://sati-jalab.org/seurat/).

The cell-cycle phase was determined by *CellCycleScore*() function, using "RRM2", "PCNA", "SLBP", "WDR76", "MCM5" as S-phase genes and "CENPF", "TPX2", "HMGB2", "UBE2C", "BUB1B", "TOP2A", "CENPE", "TACC3", "BUB1", "AURKA", "CDC20" as G2M genes, and the mitochondrial percentage of each cell was calculated with *PercentageFeatureSet*(*pattern= “^MT-”).* For chicken poor-quality cells and doublets were first filtered considering the number of detected genes (< 8000 and > 1500) and mitochondrial percentage (> 3%). Both for chicken and mouse a manual identification of poor-quality clusters were performed and those cells were excluded for next steps. Samples of equal temporal stages were merged into one Seurat object were subject to normalize and scale, and regress-out cell-cycle variation.

#### Cluster identification

To group cells by transcriptome similarity, expression data was linearly reduced into principal components (*RunPCA*, default parameters). A shared nearest neighbor (SNN) graph was calculated from the first principal components that explains almost 90% of the variability or the %variability explained by the next PC is less than 5% with *FindNeighbors*, default parameters. Based on this graph, the Louvain algorithm allows to detect communities or clusters with multi-level tuning of the resolution parameter (*FindClusters*, resolution (0.05, 0.1, 0.2, 0.3, 0.4, 0.5, 0.6, 0.8, 1.0, 1.2, 1.8, 2.4)). This resolution varied from lower values, to identify general cell types (e.g. “glutamatergic neurons”); to higher values, to identify cell subtypes (e.g. “MGE-derived GABAergic interneurons”). This resolution parameter was optimized through iteration to obtain equivalent number of clusters or cell types despite cell number differences between species. Finally, cells were represented into a two-dimensional space by the non-linear dimensional reduction technique *Uniform Manifold Approximation and Projection* (*RunUMAP*) and also *t-Distributed Stochastic Neighbor Embedding (tSNE)*.

To identify the neurobiological cell identity of each cluster, differential expression analysis was carried out among clusters (*FindAllMarkers*, *min.pct = 0.25, logfc.threshold = 0.25*). Scientific literature, *in situ* hybrization databases (Allen Brain Institute, Geisha Arizona, Mouse Genome Informatics), single cell equivalent experiments from the literature (*27*, *64– 66*) and our own *in situ* sequencing experiments of the resultant cluster biomarkers were used to assign cell type identities. In both species’ datasets, we first filtered out cells not interesting for the comparison, such as some astrocytes (expressing *Aqp4*, *Egf* genes) and oligodendrocytes (*Olig2*, *Pdgfra*) as well as cells that did not survive well the FACS sorting required for BirthSeq (and which showed features of low quality).As said before, there was a first identification at low resolution, where neurons and neural progenitors were subsetted as cells of interest; and a high resolution, where only cells of interest were classified into more specific subtypes.

### Comparative transcriptomic analysis

Before performing all interspecies comparisons, all datasets underwent 1-to-1-orthology filtering as the state-of-the-art gene equivalency. The identification of paired 1-to-1-orthologues was carried out by *Ensembl* interface in R, *biomaRt* (v2.48.3). This tool downloaded gene identifiers in different notations (*external_gene_name, ensembl_gene_id_symbol*) and their respective paired orthologues (*with_mmusculus_homologue / with_ggallus_homologue*, filtered by 1-to-1-orthology). Once paired orthologues were stablished, raw counts and metadata were extracted from each *Seurat* object, 1-to-1-orthologues were subset and transform into a common nomenclature (Mouse *external_gene_name*). In case of additionally comparing gene ontologies, as transcription factors (TFs), the gene list matching with TFs GOs (*’GO:0006366’, ’GO:0000981’, ’GO:0003700’, ’GO:0006383’, ’GO:0000995’, ’GO:0001228’, ’GO:0001227’*) was obtained from *biomaRt* and datasets are additionally subsetted.

For the interspecies comparisons, three complementary statistical analysis were carried out: Multi-Species Integration, Cluster-specific DE module expression analysis, *Gene Specificity Index* (*GSI*) Correlation (*11*) and *Label Transfer* (*67*). A model of the code used is uploaded in https://github.com/phylobrain/Rueda-Alana2024

#### Multi-Species Integration

For this approach, each species dataset was regarded as a replicate, so all 1-to-1 orthologues datasets were integrated into a joint Seurat object by *SCTransform* (v0.3.3) (*68*). This tool normalizes data, obtains variable features and scales merged datasets considering reads and feature numbers, mitochondrial percentage and cell cycle scores as regressors. The mathematical method used for this integration was the *Reciprocal Principal Components Analysis* (*RPCA*), which is more conservative and faster than *Canonical Correlation Analysis* (*CCA*). In RPCA, replicates’ expression data were converged into a common reduced dimension space and stablished nearest neighbors (cells) to remove batch effects. All parameters were set by default, except for the following ones in: *FindIntegrationAnchors (reduction= “rpca”, normalization.method = “SCT”, dims = 1:50)*, and equivalent to the rest of the pipeline. After the identification of clusters in this embedding, we assessed the conservation of an specific cell type regarding the proportion of cells of each species. Being highly conserved if balanced, and diverse if species specific.

#### Cluster-specific DE module expression analysis

To evaluate the level of transcriptome conservation, we assessed the module expression of each equivalent clusters. Each module was created by PercentageFeatureSet() out of the four most differentially expressed genes for each species cell type. Although each gene set was determined independently of the integration by FindAllFeatures in each individual species, the expression module was calculated out of the normalized experiment (“SCT) of both. A modified version of DotPlot() function was employed to visualize these modules, this version normalized all expression modules with respect to its cell type module expression (*me_i,c_* = *me_i_^/^me_c_*). In this way, 1 is assigned to the module expression of its cell type and the rest of cell types have proportional expression. Secondly, the new single cell UMAP was evaluated by the species distribution into the new generated clusters was sized up to determine the conservation of each integrated cell type.

#### *GSI* correlation

The *GSI* correlation analysis starts from the raw counts datasets, from which differential gene expression analysis was carried out to obtain a maximum of 400 markers per cell cluster. From these differentially expressed genes, only those in common for both species were selected. Finally, the raw counts datasets were filtered by this gene set and gene specificity indexes are calculated. These indexes (*S_g,c_*) were estimated as:

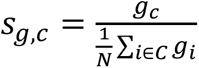

where *g*_c_ is the average gene expression for a cluster or cell type, while 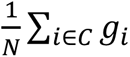, in the denominator, is the average expression for a gene across all cell types. From this matrix of indexes gene and cluster specific, a correlation matrix was obtained by the *Spearman’s* coefficient (ρ, Rho). The visualization package for these matrices is *corrplot* (v0.92).

#### Label transfer

The label transfer tool (*67*) is integrated in *Seurat* utilities and analyses the similarity degree of cells based on their whole transcriptome. This method considers one of the datasets as the reference atlas, and the other the query. Thus, cells from the query were projected into reference *PCA* embedding and are predicted cell type identities based on the chosen reference. Before this prediction, as detailed for individual stage-species dataset, the first step is to pre-process the 1-to-1-orthologues datasets (*SCTransform, PCA, SNN, UMAP*…). Then, execute *FindTransferAnchors* and *TransferData* for projecting one species into the other. This method can be sensible to number of cells, so although the transference was performed in both directions, the most reliable transference was with the larger dataset as reference. *Label Transfer* functions are similar to *FindIntegrationAnchors* and *IntegrateData* in the integration approach described before. However, these functions are more conservative to avoid overfitting and additionally generate a predicted expression matrix for each query.

With this weighted matrix, the algorithm predicted a new reference cell type out of each query cell. To visualize these predictions, we summarized into a matrix the percentage of cells of a query cell type predicted into certain reference cell type. The visualization package for these matrices is *corrplot*, modified to express the transference confidence related to each prediction as size.

### In situ sequencing (ISS)

A gene panel for chicken brain was generated using E15 chick scRNAseq clustered data as a starting point. We first selected, for each cluster, a number of informative markers based on cluster-specific differential expression. From this list we then picked a subset list of genes within an optimal expression range, estimated based on the known capture efficiency of ISS. We finally assessed in silico our ability to reproduce the original scRNAseq clusters using only the subset gene list, and further shortened the list by an iterative process, keeping a minimum number of markers that were sufficient to recapitulate the scRNAseq clusters. We then complemented this gene list with a manually curated list of known markers to identify anatomical hallmarks and broad cell types.

The probes were ordered as DNA Ultramers with a 3’ terminal RNA base from Integrated DNA Technologies (IDT), with a synthesis scale of 4nmol, resuspended in IDTE buffer at a 200 uM concentration, then pooled in equimolar amount, and phosphorylated according to the following protocol: 20 nmol of the pooled probes were phosphorylated in 1X PNK buffer (B0201S, New England Biolabs), 1 mM ATP (P0756S, New England Biolabs) and 20 units of T4 Polynucleotide kinase (M0201L, New England Biolabs), in a total volume of 50 uL, for 2 hours at 37 degrees C. PNK was then heat-inactivated at 65°C for 5 minutes and the pooled phosphorylated probes were stored at -20°C until used. The sequence of all PLPs is available in **External Database S1**

#### Direct RNA ISS

A detailed step-by-step protocol for RNA-ISS is provided here following (*69*): (https://www.protocols.io/view/home-made-direct-rna-detection-kqdg39w7zg25/v1?version_warning=no).

Tissue samples (E15 chicken telencephala of E4-birthdated animals) stored at -80°C were allowed to reach RT, washed with PBS and progressively dehydrated using one wash each of 2’in 70% ethanol and 100% ethanol. SecureSeal™ Hybridization Chambers were applied around tissue sections. Sections were then fixed with 3,7% formaldehyde for 5 min, and washed with PBS. After fixation, sections were permeabilized with 0.1M HCl for 5 min and washed with PBS-tween 0,1%. Next, phosphorylated PLPs were hybridized at a final concentration of 10 nM/PLP in PLP hybridization buffer (2X SSC, 5% ethylene carbonate, 15 mM MgCl_2_, 10 U/µl RiboProtect) overnight at 37°C. Excess probes were washed with 2x washes in 10% formamide, and a ligation was performed at 37 °C for 2 hours with 0.5 U/uL of T4 RNA ligase 2 in a home-made buffer whose composition is indicated in the link above. Rolling circle amplification (RCA) was then performed with phi29 polymerase overnight at 30°C. L-probes were hybridized at a concentration of 125 nM for 30 minutes at room temperature in L-probe hybridization buffer (2X SSC, 20% ethylene carbonate), followed by the hybridization of readout detection probes (100 nM) and DAPI for 30 minutes at room temperature in the same hybridization buffer. Sections were washed with PBS and mounted with SlowFade Gold Antifade Mountant.

#### Iterative Imaging

Imaging was performed using a Leica DMI8 epifluorescence microscope connected to an external LED source (Lumencor® SPECTRA X light engine). Light engine was set up with filter paddles (395/25, 470/24, 555/28, 635/22, 730/40). Images were obtained with a sCMOS camera (2048 × 2048, 16-bit, Leica DFC90000GTC-VSC10726), automatic multi-slide stage, and Leica Apochromat objective 40× (HC PL APO 40x/1.10 WATER, 11506342). The microscope was equipped with filter cubes for 5 dye separation (AF750, Cy5, Cy3, AF488, and DAPI) and an external filter wheel (DFT51011).

Each region of interest (ROI, usually a whole telencephalon in coronal section, comprising one hemisphere) was selected manually and saved in the Leica LASX software, for repeated imaging. Each ROI was automatically subdivided into tiles, and for each tile a z-stack with an interval of 0.5 micron was acquired in all the channels. The tiles are defined so to have a 10% overlap at the edges. The images were saved as thousands of individual tiff files with associated metadata.

After each imaging cycle, a stripping step by 3 washes in 100% formamide was performed, and a new set of L-probes and detection oligos was hybridized following a combinatorial labeling strategy (*70*).

EdU detection was performed after the ISS cycles. First, all the probes were stripped using 3x 100% formamide washes. We then detected the EdU labeling using a rat-raised antibody against BrdU (ab6326, AbCam, cross-reactive to EdU), followed by labeling with a secondary donkey anti-rat AF750-labeled antibody (ab175750, AbCam).

#### Image analysis

The raw images and the metadata from the microscopes were fed into an in-house pre-processing software module (https://github.com/Moldia/Lee_2023/tree/main/ISS_prepro-cessing). The module performs the following steps: first the images are maximum projected. The projected images are simultaneously stitched and aligned across imaging cycles, using ASHLAR (*71*). The aligned stitched images are sliced into smaller tiles, to allow a computationally efficient decoding.

On these sliced images (excluding the DAPI channel), we applied Content-Aware Image Restoration (*72*), following https://github.com/Moldia/Lee_2023/tree/main/ISS_CARE, and using a custom denoising model specifically trained on raw-deconvolved paired of ISS image datasets, spanning different tissues, organisms, and fluorophores. The model is also deposited in the same repository.

The resliced denoised images were then fed into an in-house decoding software module (https://github.com/Moldia/Lee_2023/tree/main/ISS_decoding), which converts them into the SpaceTX format (*73*), and pipes them into the Starfish Python library for decoding of image-based spatial transcriptomics datasets (https://github.com/spacetx/starfish). Here in the decoding module, the images are normalized across channels and imaging cycles, ISS spots are found and decoded, and quality metrics are computed.

All the downstream analysis was performed using the code in this repository (https://github.com/Moldia/Lee_2023/tree/main/ISS_postprocessing). Decoded expression spots were assigned to individual cells by first generating a segmentation mask based on the DAPI signal using stardist (*74*). Each segmented nucleus was then expanded up to a maximum of 20 pixels in each direction, unless another expanding nucleus was found. This slightly expanded segmentation mask and the decoded spots table, as well as the reference BirthSeq scRNAseq dataset, were passed as inputs to a Python implementation of Probabilistic Cell Typing (*75*).

The non-expanded segmentation mask was also applied to the EdU images, and was used to extract EdU intensity values for each nucleus. The median intensity value per nucleus was computed and, for each section, cells falling over a given median intensity threshold were labeled as EdU-positive.

#### ISS data analysis

In order to integrate ISS data with the pre-existing single-cell RNAseq datasets, and to map in-situ the scRNAseq clusters, we used Probabilistic Cell Typing (PCIseq, (*75*)). A cluster-by-gene average expression matrix inferred from the clustered scRNAseq data was fed, together with a segmentation mask and the complete ISS dataset, to the PCIseq algorithm, following the implementation at (https://github.com/acycliq/pciSeq). After the assignment, low probability cells were eliminated (p<0.6) and the remaining cells were plotted, cluster by cluster, over a DAPI image to analyze their anatomical distribution.

### Mathematical modelization

All details of the mathematical modelization are specified in the **Supplementary Notes S1**, including specific references.

## Supporting information

Supplementary notes on the maths model

## Acknowledgments

We thank Prof. Lukas Kratochvil for their significant contribution, which allowed the establishment of our gecko colony; Prof. Henrik Kaessmann and Bastienne Zaremba (Univ. Heidelberg – Germany) for creative and helpful comments on the manuscript; the Servicio General de Microscopía Analítica y de Alta Resolución en Biomedicina, SGIker at the UPV/EHU; the ISS unit at SciLifeLab, who provided some ISS imaging data. We gratefully thank those many researchers who have discussed the data with our team all over these years.

## Author contributions

ERA and FGM conceived the project, interpreted the data, designed and performed experiments and data analysis, and wrote the manuscript with input from all authors. MTGF and AF performed embryo work and experiments. AMA, AB and AQG performed single cell RNA sequencing. EV and RSG performed bioinformatic analysis. LE, MR and MdMV performed FACS sorting analysis. MG, SMS and ERA performed and analyzed the *in situ sequencing* experiments. JLF, DG and AT performed and analyzed in situ hybridization. MP and FGM performed theoretical modeling. MN, AD, JME and FGM provided reagents.

## Data and materials availability

Data generated for this study have been submitted to the NCBI Gene Expression Omnibus (GEO; https://www.ncbi.nlm.nih.gov/geo/) under accession number GSEXXXXXX. The code used for analysis (scRNAseq proccessing, interspecies comparisons, spatial transcriptomics) can be found in https://github.com/phylobrain/Rueda-Alana2024.

## Supplementary Materials

**Extended materials and Methods**

**Supplementary Figures S01-S22 Tables S1-S4**

**Supplementary notes S1**

**External Databases S1**

## List of abbreviations

Acc: accumbens nucleus
aDVR: anterior DVR
APall: arcopallium
Arc: arcuate nucleus
CGE: caudal ganglionic eminence
CP: cortical plate
CPN: cortical projecting neurons
CthPN: cortico-thalamic projecting neurons
DC: dorsal cortex
Di: diencephalon
DLH: dorsolateral hypothalamic nucleus
DMC: dorsomedial cortex
dNPall: dorsal nidopallium
DPall: dorsal pallium
DT: dorsal thalamus
DVR: dorsal ventricular ridge
EPall: entopallium/visual core
GABA: GABAergic neuron
GLUT: glutamatergic neuron
GZ: germinal zone
HA: hyperpallium apicale
Hb: habenular complex
Hc: hippocampus
HD: densocellular hyperpallium
HI: intermediate hyperpallium
HPall: hyperpallium
IHA: intercalated hyperpallium
IIIv: third ventricle
IPC: intermediate precursor cell
IZ: intermediate zone
L1: cortical layer 1 of the gecko
L2: cortical layer 2 of the gecko
L2/3: layer 2 and 3 of the neocortex
L3: cortical layer 3 of the gecko
L4: layer IV of the neocortex
L5/6: layer V of the neocortex
L6b: subplate/layer 6b of the neocortex
LC: lateral cortex
LC: lateral cortex
lfb: lateral forebrain bundle
LGE: lateral ganglionic eminence
LGV: lateral geniculate nucleus, ventral part
LPall: lateral pallium
Lv: lateral ventricle
M: nucleus medialis
MC: medial cortex
mfb: medial forebrain bundle
MGE: medial ganglionic eminence
MP: mesopallium
MPall: medial pallium
MZ: marginal zone
NC: nidopallium caudale
NCx: neocortex
NPall: nidopallium
NSM: medial septal nucleus
NSL: lateral septal nucleus
ot: optic tract
pDVR: posterior DVR
PH: paraventricular hypothalamic nucleus
PT: pallial thickening
RGC: radial glial cell
Rot: rotundus nucleus
SCPN: subcortical projecting neurons
Se: septum
SP: subplate of the neocortex
SPall: subpallium
Sph: nucleus sphericus
Str: striatum
TF: transcription factor
Thal: thalamus
ThR: Thalamic recipient area
UMAP.: Uniform Manifold Approximation and Projection
VMH: ventromedial hypothalamic nucleus
VPall: ventral palliu

