## Supplementary figures and tables for "Evolutionary convergence of sensory circuits in the pallium of amniotes"

**The PDF includes:**

Materials and Methods

Supplementary Tables S1-S4

Figures S01-S22

**Other Supplementary Material for this manuscript includes the following:**

External database S1

Supplementary notes S1

### Supplementary Tables

**Table S1:** Summary of the statistical tests performed on the DVR and hyperpallium birthdating analysis.

| Figure | Summary of statistical analyses |
|---|---|
| **Figure 1** | In [H], One-way ANOVA followed by all pairwise multiple comparisons by Holm-Sidak post-hoc test was performed for E4 and E8; for E6, Kruskall-Wallis One-way ANOVA followed by all pairwise multiple comparisons (Dunn´s method) was performed. In [I], One-way ANOVA followed by all pairwise multiple comparisons by Holm-Sidak post-hoc test was performed for APall and EPall; for dNPall, Kruskall-Wallis One-way ANOVA followed by all pairwise multiple comparisons (Dunn´s method) was performed. |
| **Figure 2** | In [I], One-way ANOVA followed by all pairwise multiple comparisons by Holm-Sidak post-hoc test was performed. In [J], datasets did not comply ANOVA assumptions (normality for IHA and HI/HD; and homoscedasticity for HA), so they were logarithmically transformed before the analysis. For all transformed data sets, One-way ANOVA followed by all pairwise multiple comparisons by Holm-Sidak post-hoc test was performed. |
| **Figure S01** | In [N], One-way ANOVA followed by all pairwise multiple comparisons by Holm-Sidak post-hoc test was performed. In [P], One-way ANOVA followed by all pairwise multiple comparisons by Holm-Sidak post-hoc test was performed for E6; for E4 and E8, Kruskall-Wallis One-way ANOVA followed by all pairwise multiple comparisons (Dunn´s method) was performed. E6 dataset did not comply ANOVA normality assumption, so it was logarithmically transformed before the analysis. In [Q], One-way ANOVA followed by all pairwise multiple comparisons by Holm-Sidak post-hoc test was performed for EPall and dNPall; for APall Kruskall-Wallis One-way ANOVA followed by all pairwise multiple comparisons (Dunn´s method) was performed. dNPall dataset did not comply ANOVA normality assumption, so it was logarithmically transformed before the analysis. |
| **Figure S02** | In [N], One-way ANOVA followed by all pairwise multiple comparisons by Holm-Sidak post-hoc test was performed. In [P], One-way ANOVA followed by all pairwise multiple comparisons by Holm-Sidak post-hoc test was performed for E4 and E8; for E6, Kruskall-Wallis One-way ANOVA followed by all pairwise multiple comparisons (Tukey test) was performed. E8 dataset did not comply ANOVA normality assumption, so it was logarithmically transformed before the analysis. In [Q], One-way ANOVA followed by all pairwise multiple comparisons by Holm-Sidak post-hoc test was performed for APall, EPall and dNPall. EPall and dNPall datasets did not comply ANOVA homoscedasticity assumption, so they were logarithmically transformed before the analysis. |
| **Figure S03** | In [N], One-way ANOVA followed by all pairwise multiple comparisons by Holm-Sidak post-hoc test was performed. In [P], One-way ANOVA followed by all pairwise multiple comparisons by Holm-Sidak post-hoc test was performed for E6 and E8; for E4, Kruskall-Wallis One-way |

| | |
|---|---|
| | ANOVA was performed. E8 dataset did not comply ANOVA normality assumption, so it was logarithmically transformed before the analysis. In [Q], One-way ANOVA followed by all pairwise multiple comparisons by Holm-Sidak post-hoc test was performed for APall, EPall and dNPall. APall and dNPall datasets did not comply ANOVA normality assumption, so they were logarithmically transformed before the analysis. |
| **Figure S04** | In [N], One-way ANOVA followed by all pairwise multiple comparisons by Holm-Sidak post-hoc test was performed. In [P], One-way ANOVA followed by all pairwise multiple comparisons by Holm-Sidak post-hoc test was performed for E4, E6 and E8. E6 dataset did not comply ANOVA homoscedasticity assumption, so it was logarithmically transformed before the analysis. In [Q], One-way ANOVA followed by all pairwise multiple comparisons by Holm-Sidak post-hoc test was performed for APall, EPall and dNPall. dNPall dataset did not comply ANOVA homoscedasticity assumption, so it was logarithmically transformed before the analysis. |
| **Figure S07** | In [M], Kruskall-Wallis One-way ANOVA followed by all pairwise multiple comparisons (Tukey Test) was performed. In [N], Student´s t test was performed for E4 HA, E4 IHA, E4 HI/HD, E6 HA, E6 IHA, E6 HI/HD, E8 IHA and E8 HI/HD; for E8 HA Mann-Whitney Rank Sum Test was performed. E6 HA and E6 IHA datasets did not comply t-test normality assumption, so they were logarithmically transformed before the analysis. In [P], One-way ANOVA followed by all pairwise multiple comparisons by Holm-Sidak post-hoc test was performed for E6 and E8; for E4, Kruskall-Wallis One-way ANOVA followed by all pairwise multiple comparisons (Tukey Test) was performed. E6 and E8 datasets did not comply ANOVA normality assumption, so they were logarithmically transformed before the analysis. In [Q], One-way ANOVA followed by all pairwise multiple comparisons by Holm-Sidak post-hoc test was performed for IHA; for HA and HI/HD Kruskall-Wallis One-way ANOVA followed by all pairwise multiple comparisons (Dunn´s method) was performed. IHA dataset did not comply ANOVA normality assumption, so it was logarithmically transformed before the analysis. |
| **Figure S08** | In [M], One-way ANOVA followed by all pairwise multiple comparisons by Holm-Sidak post-hoc test was performed. In [N], Student´s t test was performed for E4 HA, E4 IHA, E6 HA and E6 IHA; for E4 HI/HD, E6 HI/HD, E8 HA, E8 IHA and E8 HI/HD Mann-Whitney Rank Sum Test was performed. E4 IHA dataset did not comply t-test normality assumption, so it was logarithmically transformed before the analysis. In [P], One-way ANOVA followed by all pairwise multiple comparisons by Holm-Sidak post-hoc test was performed for E4; for E6 and E8, Kruskall-Wallis One-way ANOVA was performed. E4 dataset did not comply ANOVA normality assumption, so it was logarithmically transformed before the analysis. In [Q], One-way ANOVA followed by all pairwise multiple comparisons by Holm-Sidak post-hoc test was performed for IHA and HI/HD; for HA Kruskall-Wallis One-way ANOVA followed by all pairwise multiple comparisons (Dunn´s method) was performed. IHA and HI/HD dataset did not comply ANOVA normality and homoscedasticity assumptions, so they were logarithmically transformed before the analysis. |
| **Figure S09** | In [M], Kruskall-Wallis followed by all pairwise multiple comparisons (Tukey Test) was performed. In [N], Student´s t test was performed for E4 HI/HD, E6 HA, E6 IHA, E6 HI/HD, E8 |

| | |
|---|---|
| | IHA, E8 HI/HD and E8 HA; for E4 HA and E4 IHA Mann-Whitney Rank Sum Test was performed. E8 HA dataset did not comply t-test normality assumption, so it was logarithmically transformed before the analysis. In [P], One-way ANOVA followed by all pairwise multiple comparisons by Holm-Sidak post-hoc test was performed for E6 and E8; for E4, Kruskall-Wallis One-way ANOVA was performed. E8 dataset did not comply ANOVA normality assumption, so it was logarithmically transformed before the analysis. In [Q], One-way ANOVA followed by all pairwise multiple comparisons by Holm-Sidak post-hoc test was performed for HA, IHA and HI/HD. HA, IHA and HI/HD datasets did not comply ANOVA normality and homoscedasticity assumptions, so they were logarithmically transformed before the analysis. |
| **Figure S10** | In [M], One-way ANOVA followed by all pairwise multiple comparisons by Holm-Sidak post-hoc test was performed. Dataset did not comply ANOVA normality assumption, so it was logarithmically transformed before the analysis. In [N], Student´s t test was performed for E4 HA, E4 IHA, E4 HI/HD, E6 HA, E6 IHA, E6 HI/HD and E8 HI/HD; for E8 HA and E8 IHA Mann-Whitney Rank Sum Test was performed. E6 HI/HD dataset did not comply t-test normality assumption, so it was logarithmically transformed before the analysis. In [P], One-way ANOVA followed by all pairwise multiple comparisons by Holm-Sidak post-hoc test was performed for E4, E6 and E8. E8 dataset did not comply ANOVA normality assumption, so it was logarithmically transformed before the analysis. In [Q], One-way ANOVA followed by all pairwise multiple comparisons by Holm-Sidak post-hoc test was performed for HA, IHA and HI/HD. IHA dataset did not comply ANOVA normality assumption, so it was logarithmically transformed before the analysis. |

**Table S2** –Primers for the generation of riboprobes to detect the expression of mRNA in gecko brains. Based on *Paroedura picta* genome version 1 (*76*) (Ppicta_assembly_v1, https://transcriptome.riken.jp/reptiliomix/resources.html).

| Gene | Forward | Reverse |
|---|---|---|
| **etv1** | 5´-TCTGGCTCATGACTCAGAAG-3´ | 5´-CGTCCTTCCCTTGGCATTGTG-3´ |
| **sulf2 (a)** | 5´-GCGAGTCAAAGATCTCTGCC-3´ | 5´- CGAGGAGAATGAGTACAAGC-3´ |
| **sulf2 (b)** | 5´-TGCCTCAGTTTGCCCTTGTG-3´ | 5´-GGTTGTCATGGGTGAAGCAG-3´ |
| **rorb** | 5´-AGCTCCAGGAATAACCATGG-3´ | 5´-TCTTGGTTCCAACAGCCAAG-3´ |
| **dbp** | 5´-AGGAGGAGCTGAAACCACAG-3´ | 5´-CTCGTTCCAAACATCCGTTG-3´ |
| **satb1 (a)** | 5´-TCACAGCTCTGCTGCCCAAG-3´ | 5´-ACTCGAAGACTTGCCTCCTG-3´ |
| **satb1 (b)** | 5´-TCCTTTCCCTTTCATCCTGG-3´ | 5´-CCTTTCGCAGGATTTCTGAG-3´ |

**Table S3 -** Conditions used for the enzymatic tissue dissociation for scRNAseq experiments.

| Species | Stage | Enzyme | [Enzyme] | Solution | Incubation conditions |
|---|---|---|---|---|---|
| Chick | E6 | Proteinase K | 50 µg/ml | Enzymatic solution (116 mM NaCl, 5.4 mM KCl, 26 mM $NaHCO_3$, 1 mM $NaH_2PO_4$, 1.5 mM $CaCl_2$, 1 mM $MgSO_4$, 0.5 mM EDTA, 25 | 15 min at 37 °C |
| | E15 | | | | 20 min at 37 °C |

| | | | | | |
|---|---|---|---|---|---|
| | | | | mM glucose, 1 mM L-cysteine) containing **150 U/ml DNase I**. | |
| Mouse | E14 | TrypLE Express | 1X (confidential) | TrypLE Express commercial media (2.67 mM KCl, 1.47 mM $KH_2PO_4$, 137.93 mM NaCl, 8.06 mM $Na_2HPO_4 \cdot 7 H_2O$, 1.1 mM EDTA). | 10 min at 37 °C |
| | P3 | Papain | 11.3 U/ml | Enzymatic solution (116 mM NaCl, 5.4 mM KCl, 26 mM $NaHCO_3$, 1 mM $NaH_2PO_4$, 1.5 mM $CaCl_2$, 1 mM $MgSO_4$, 0.5 mM EDTA, 25 mM glucose, 1 mM L-cysteine) containing **150 U/ml DNase I**. | 15 min at 37 °C |

**Table S4**. Quality control statistics related to this alignment steps and others relevant features pf the sequenced libraries.

| **Species** | **MOUSE** | | | | **CHICKEN** | | | |
|---|---|---|---|---|---|---|---|---|
| **Day** | Day1 | | Day2 | | Day1 | | Day2 | |
| **Replicate** | Rep1 | Rep2 | Rep1 | Rep2 | Rep1 | Rep1 | Rep2 | Rep3 |
| **Facility** | CICbioGUNE | | CNIC | | CNIC | | CICbioGUNE | |
| **Reads Mapped to Genome** | 95,40% | 96% | 96,70% | 96,80% | 92,20% | 91,60% | 91,80% | 92,10% |
| **Reads Mapped Conidently to Transcriptome** | 58,80% | 58,90% | 63,40% | 59,90% | 53,30% | 48,80% | 43,60% | 49,10% |
| **Number of Reads** | 484789730 | 397651643 | 380632455 | 421186775 | 351753820 | 351939123 | 417367984 | 463669373 |
| **Estimated Number of Cells** | 19101 | 19046 | 13858 | 14513 | 23903 | 10361 | 8812 | 10214 |
| **Median Genes per Cell** | 2061 | 1913 | 2549 | 2784 | 436 | 600 | 868 | 866 |
| **Mean Reads per Cell** | 25380 | 20878 | 27467 | 29021 | 14716 | 33968 | 47364 | 45395 |

### Supplementary Figures

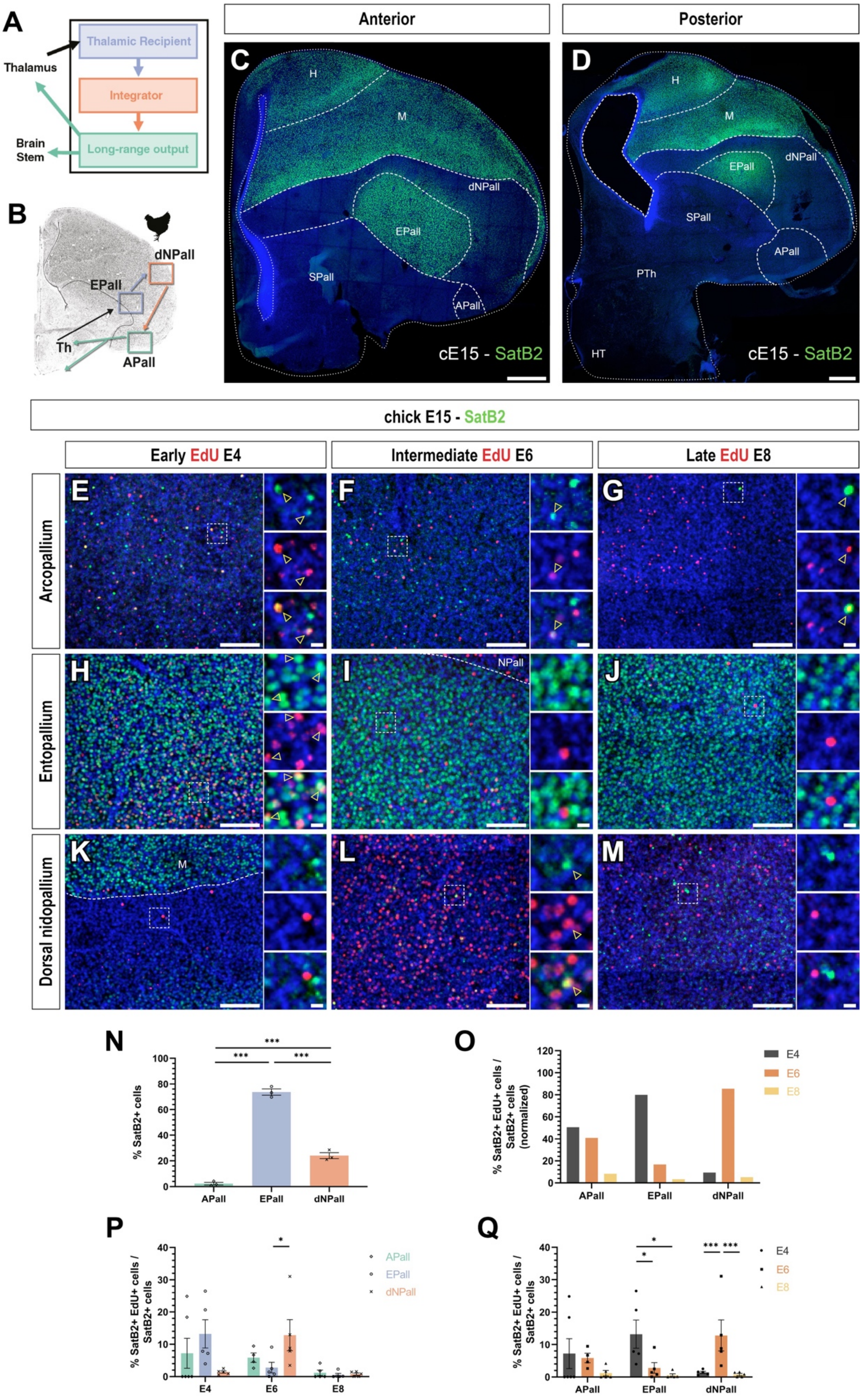

**Figure S01 – Early and intermediate neurogenesis in the chick telencephalon generate SatB2+ glutamatergic neurons of the DVR circuit.** (**A**) Representative scheme of the canonical high-order sensory processing trisynaptic circuit. (**B**) Representative diagram of the trisynaptic circuit in the DVR. (**C-D**) Anterior and posterior coronal sections of E15 chick brains showing the distribution pattern of SatB2+ and SatB1+ glutamatergic neurons. (**E-M**) High-power view and split channels of the APall (**E-G**), EPall (**H-J**) and dNPall (**K-M**) in coronal sections of E15 chick brains birthdated at E4 (**E, H, K**), E6 (**F, I, L**) and E8 (**G, J, M**). Immunohistochemistry against SatB2 shows that these glutamatergic neurons are generated following the general neurogenic pattern of the DVR. (**N**) Quantification of the distribution of SatB2+ glutamatergic neurons throughout DVR circuit regions. (**O**) Graphic representation of the normalized percentage of neurons of each circuit role generated at early, intermediate or late neurogenic timepoints. (**P**) Quantification of the percentage of EdU-labeled SatB2+ among the total SatB2+ neurons in each of the DVR circuit regions after birthdating at different neurogenic stages. (**O**) Graphic representation of the effect of age in the generation of each of the DVR circuit regions. (**Q**) Graphic representation of DVR circuit cell distribution depending on their birthdate. For nomenclature, refer to the list of abbreviations. DAPI counterstain in blue. Dashed white lines demarcate anatomical boundaries. Yellow arrowheads indicate representative neurons co-labeled with EdU and the immunohistochemical marker. Scale bars, 500 µm (**C-D**), 100 µm (**E-M**) and 10 µm (E-M insets). *$p<0.05$; ***$p<0.001$. Statistical tests for (**N-Q**) are described in **Supplementary Materials.** Bars represent mean ± SEM. Dots in (**N**) shown the mean of the data obtained at each analyzed timepoint; whereas in (**O, P**) show individual data.

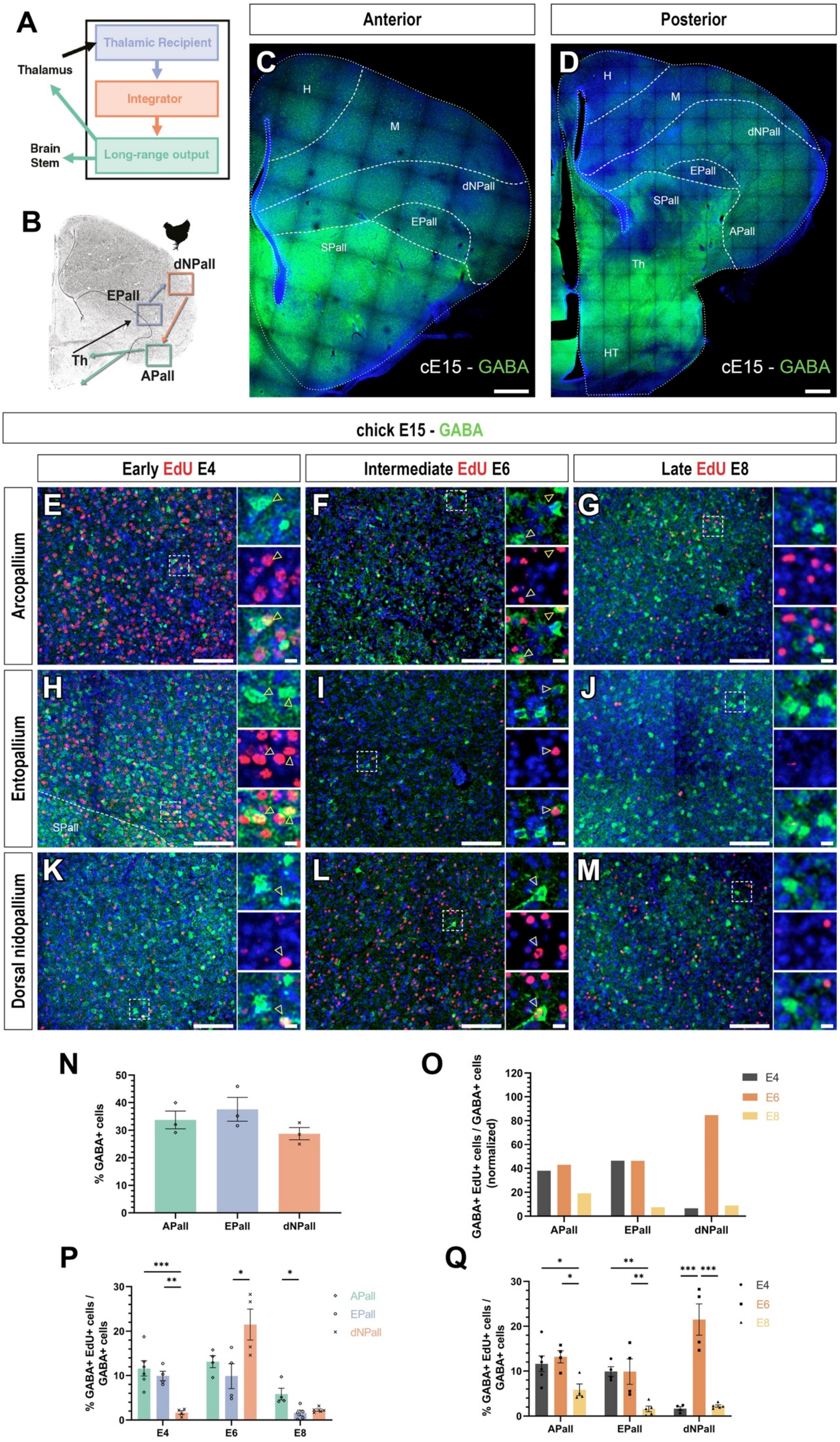
A
Thalamic Recipient
Integrator
Long-range output
Thalamus
Brain Stem
B
dNPall
EPall
Th
APall
Anterior
Posterior
C
H
M
dNPall
EPall
SPall
cE15 - GABA
D
H
M
dNPall
EPall
SPall
APall
Th
HT
cE15 - GABA
chick E15 - GABA
Early EdU E4
Intermediate EdU E6
Late EdU E8
Arcopallium
Entopallium
Dorsal nidopallium
E
F
G
H
I
J
K
L
M
SPall
N
% GABA+ cells
APall
EPall
dNPall
O
GABA+ EdU+ cells / GABA+ cells (normalized)
E4
E6
E8
APall
EPall
dNPall
P
% GABA+ EdU+ cells / GABA+ cells
APall
EPall
dNPall
E4
E6
E8
Q
% GABA+ EdU+ cells / GABA+ cells
E4
E6
E8
APall
EPall
dNPall

**Figure S02 - Chick GABA+ interneurons of the DVR circuit are born during early and intermediate neurogenesis.** (**A**) Representative scheme of the canonical high-order sensory processing trisynaptic circuit. (**B**) Representative diagram of the trisynaptic circuit in the DVR. (**C-D**) Anterior and posterior coronal sections of E15 chick brains showing the distribution pattern of GABA+ interneurons. (**E-M**) High-power view and split channels of the APall (E-G), EPall (**H-J**) and dNPall (**K-M**) in coronal sections of E15 chick brains birthdated at E4 (**E, H, K**), E6 (**F, I, L**) and E8 (**G, J, M**). Immunohistochemistry against GABA shows that these GABAergic interneurons are generated during early- and mid-neurogenic timepoints. (**N**) Quantification of the distribution of GABA+ interneurons throughout DVR circuit regions. (**O-P**) Quantification of the percentage of EdU-labeled GABA+ cells among the total GABA+ interneurons in each of the DVR circuit regions after birthdating at different neurogenic stages. (**O**) Graphic representation of the effect of age in the generation of each of the DVR circuit regions. (**P**) Graphic representation of DVR circuit cell distribution depending on their birthdate. For nomenclature, refer to the list of abbreviations. DAPI counterstain in blue. Dashed white lines demarcate anatomical boundaries. Yellow arrowheads indicate representative neurons co-labeled with EdU and the immunohistochemical marker. Scale bars, 500 µm (**C-D**), 100 µm (E-M) and 10 µm (E-M insets). $^{*}p<0.05$; $^{***}p<0.001$. In (**N**), One-way ANOVA followed by all pairwise multiple comparisons by Holm-Sidak post-hoc test was performed. In (O), One-way ANOVA followed by all pairwise multiple comparisons by Holm-Sidak post-hoc test was performed for APall, EPall and dNPall. EPall and dNPall datasets did not comply ANOVA homoscedasticity assumption, so they were logarithmically transformed before the analysis. In (**P**), One-way ANOVA followed by all pairwise multiple comparisons by Holm-Sidak post-hoc test was performed for E4 and E8; for E6, Kruskall-Wallis One-way ANOVA followed by all pairwise multiple comparisons (Tukey test) was performed. E8 dataset did not comply ANOVA normality assumption, so it was logarithmically transformed before the analysis. Bars represent mean ± SEM. Dots in (**N**) shown the mean of the data obtained at each analyzed timepoint; whereas in (**O, P**) show individual data.

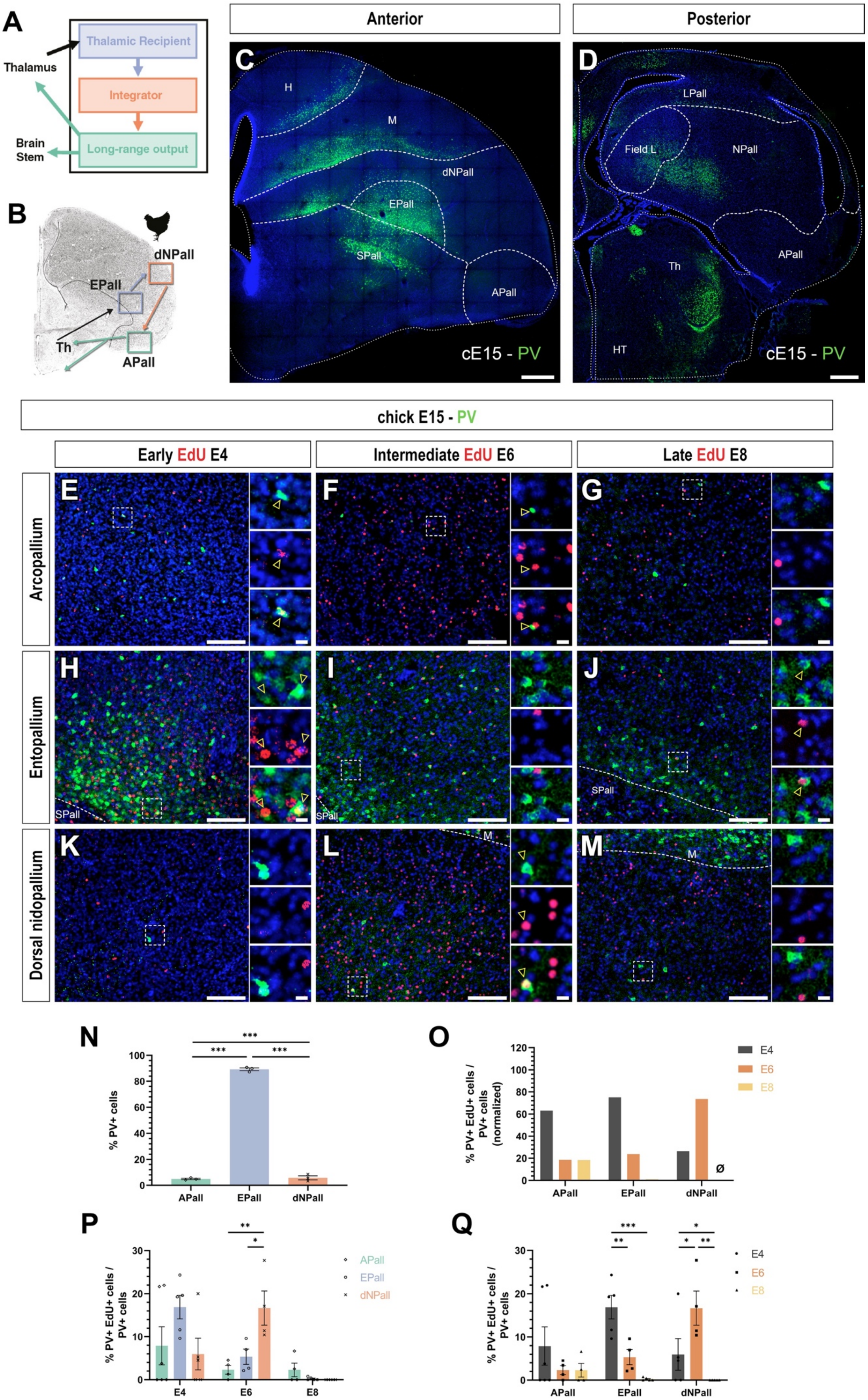
A
Thalamic Recipient
Integrator
Long-range output
Thalamus
Brain Stem
B
dNPall
EPall
Th
APall
Anterior
Posterior
C
H
M
dNPall
EPall
SPall
APall
cE15 - PV
D
LPall
Field L
NPall
Th
APall
HT
cE15 - PV
chick E15 - PV
Early EdU E4
Intermediate EdU E6
Late EdU E8
Arcopallium
Entopallium
Dorsal nidopallium
E
F
G
H
I
J
K
L
M
SPall
N
% PV+ cells
APall
EPall
dNPall
O
% PV+ EdU+ cells / PV+ cells (normalized)
E4
E6
E8
ø
P
% PV+ EdU+ cells / PV+ cells
APall
EPall
dNPall
E4
E6
E8
Q
% PV+ EdU+ cells / PV+ cells
E4
E6
E8
APall
EPall
dNPall

**Figure S03 - PV+ GABAergic interneurons of the DVR circuit are generated at early and intermediate neurogenic stages.** (**A**) Representative scheme of the canonical high-order sensory processing trisynaptic circuit. (**B**) Representative diagram of the trisynaptic circuit in the DVR. (**C-D**) Anterior and posterior coronal sections of E15 chick brains showing that PV+ GABAergic interneurons in the DVR are almost exclusively located in the EPall. (**E-M**) High-power view and split channels of the APall (**E-G**), EPall (**H-J**) and dNPall (**K-M**) in coronal sections of E15 chick brains birthdated at E4 (**E, H, K**), E6 (**F, I, L**) and E8 (**G, J, M**). Immunohistochemistry against PV shows that these GABAergic interneurons are generated following the general neurogenic pattern of the DVR. (**N**) Quantification of the distribution of PV+ glutamatergic neurons throughout DVR circuit regions. (**O-Q**) Quantification of the percentage of EdU-labeled PV+ cells among the total PV+ GABAergic interneurons in each of the DVR circuit regions after birthdating at different neurogenic stages. (**O**) Graphic representation of the normalized percentage of neurons of each circuit role generated at early, intermediate or late neurogenic timepoints. (P) Graphic representation of the effect of age in the generation of each of the DVR circuit regions. (**Q**) Graphic representation of DVR circuit cell distribution depending on their birthdate. For nomenclature, refer to the list of abbreviations. DAPI counterstain in blue. Dashed white lines demarcate anatomical boundaries. Yellow arrowheads indicate representative neurons co-labeled with EdU and the immunohistochemical marker. Scale bars, 500 µm (C-D), 100 µm (E-M) and 10 µm (**E-M** insets). *$p<0.05$; ***$p<0.001$. Statistical tests for (**N-Q**) are described in **Supplementary Materials.** Bars represent mean ± SEM. Dots in (N) shown the mean of the data obtained at each analyzed timepoint; whereas in (**O, P**) show individual data.

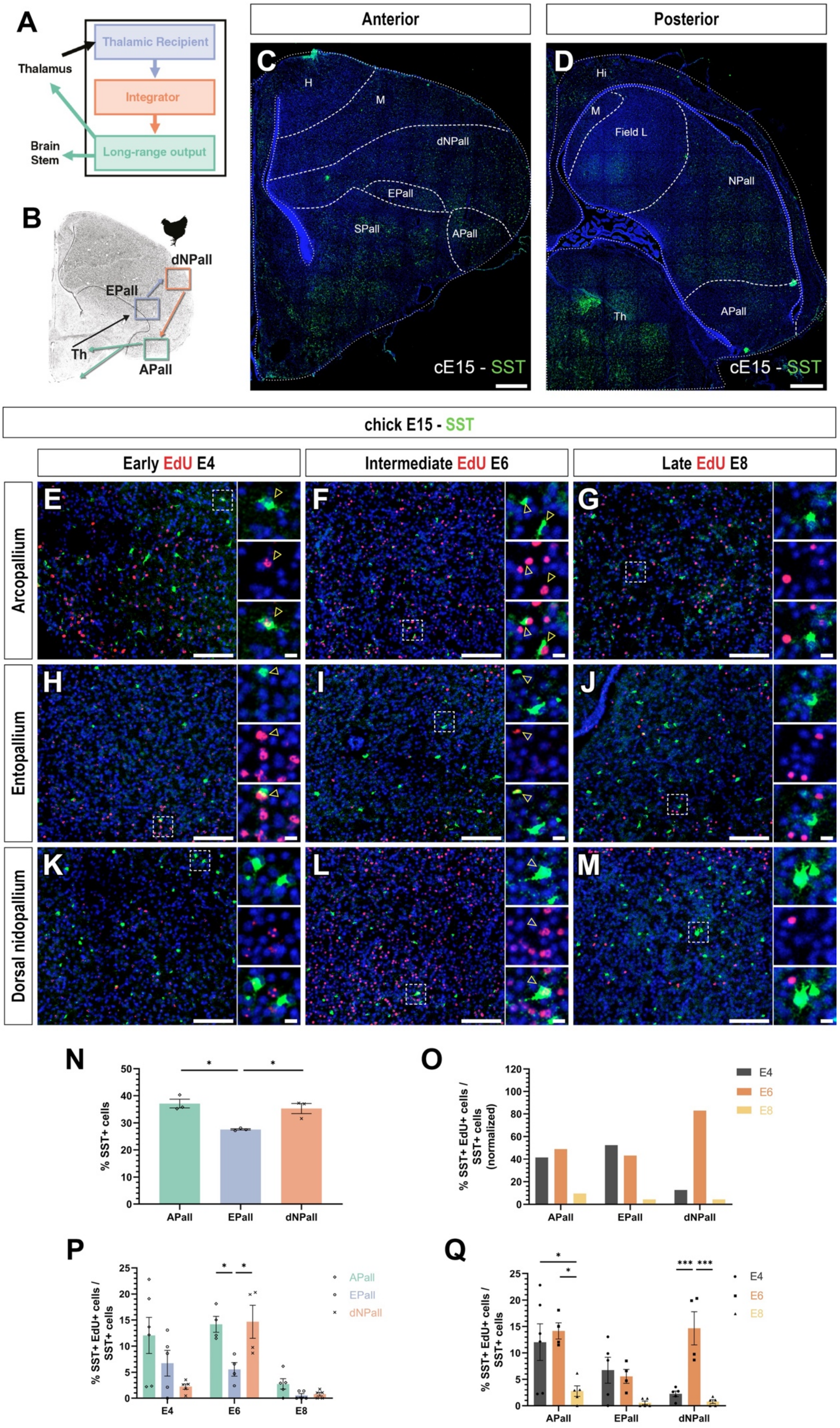
A
Thalamus
Thalamic Recipient
Integrator
Long-range output
Brain Stem
B
dNPall
EPall
Th
APall
Anterior
Posterior
C
H
M
dNPall
EPall
SPall
APall
cE15 - SST
D
Hi
M
Field L
NPall
Th
APall
cE15 - SST
chick E15 - SST
Early EdU E4
Intermediate EdU E6
Late EdU E8
Arcopallium
Entopallium
Dorsal nidopallium
E
F
G
H
I
J
K
L
M
N
% SST+ cells
APall
EPall
dNPall
O
% SST+ EdU+ cells / SST+ cells (normalized)
E4
E6
E8
APall
EPall
dNPall
P
% SST+ EdU+ cells / SST+ cells
APall
EPall
dNPall
E4
E6
E8
Q
% SST+ EdU+ cells / SST+ cells
E4
E6
E8
APall
EPall
dNPall

**Figure S04 - Early and intermediate neurogenesis in the chick telencephalon generate SST+ GABAergic interneurons of the DVR circuit.** (**A**) Representative scheme of the canonical high-order sensory processing trisynaptic circuit. (**B**) Representative diagram of the trisynaptic circuit in the DVR. (**C-D**) Anterior and posterior coronal sections of E15 chick brains showing the distribution pattern of SST+ GABAergic interneurons. (E-M) High-power view and split channels of the APall (**E-G**), Epall (**H-J**) and dNPall (**K-M**) in coronal sections of E15 chick brains birthdated at E4 (**E, H, K**), E6 (**F, I, L**) and E8 (**G, J, M**). Immunohistochemistry against SST shows that these GABAergic interneurons are generated during early and intermediate neurogenic stages. (**N**) Quantification of the distribution of SST+ GABAergic interneurons throughout DVR circuit regions. (**O-Q**) Quantification of the percentage of EdU-labeled SST+ cells among the total SST+ GABAergic interneurons in each of the DVR circuit regions after birthdating at different neurogenic stages. (**O**) Graphic representation of the normalized percentage of neurons of each circuit role generated at early, intermediate or late neurogenic timepoints. (**P**) Graphic representation of the effect of age in the generation of each of the DVR circuit regions. (**Q**) Graphic representation of DVR circuit cell distribution depending on their birthdate. For nomenclature, refer to the list of abbreviations. DAPI counterstain in blue. Dashed white lines demarcate anatomical boundaries. Yellow arrowheads indicate representative neurons co-labeled with EdU and the immunohistochemical marker. Scale bars, 500 µm (**C-D**), 100 µm (**E-M**) and 10 µm (**E-M** insets). *p<0.05; ***p<0.001. Statistical tests for (**N-Q**) are described in **Supplementary Materials.**. Bars represent mean ± SEM. Dots in (**N**) shown the mean of the data obtained at each analyzed timepoint; whereas in (**O, P**) show individual data.

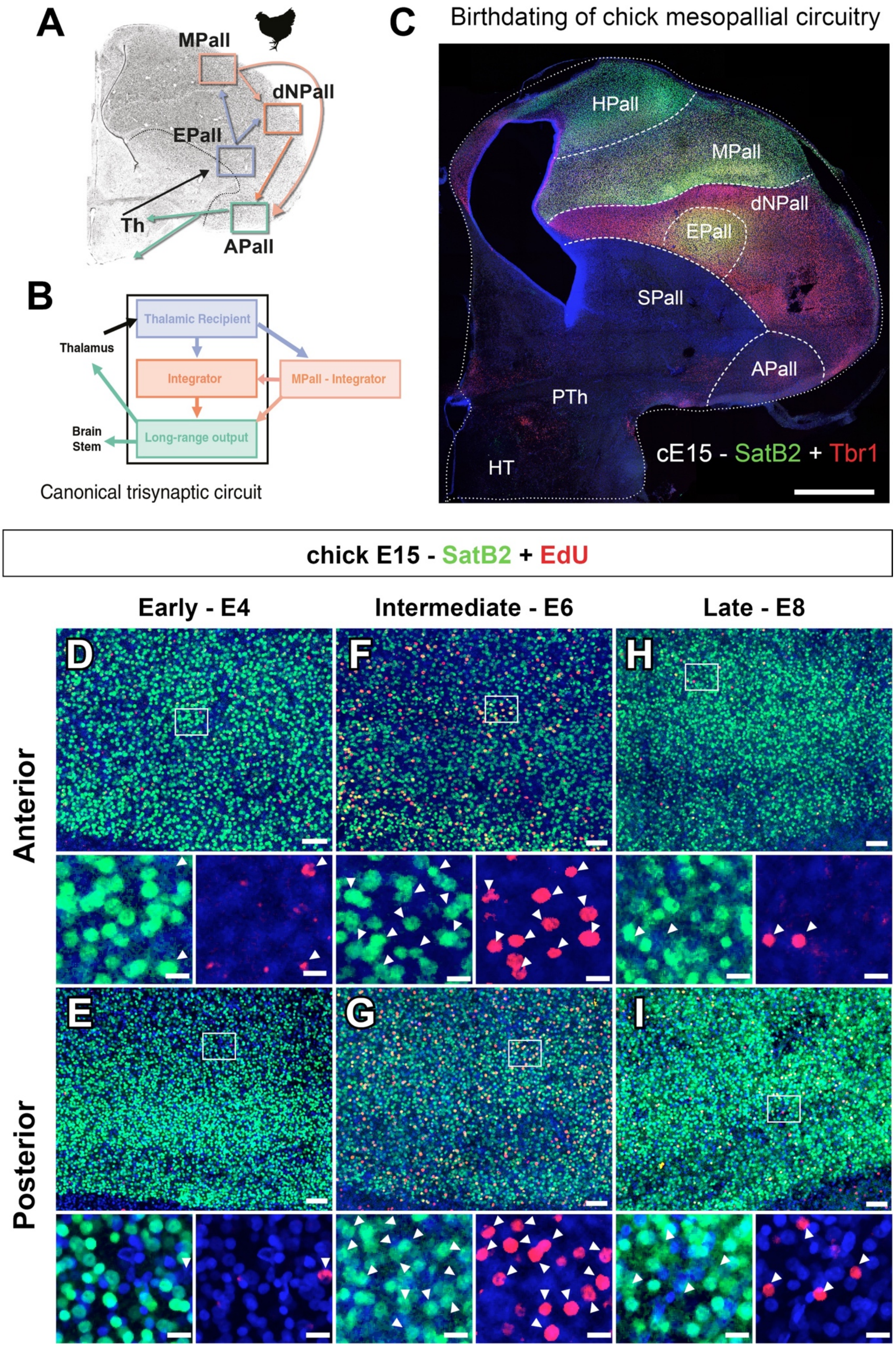
A
MPall
dNPall
EPall
Th
APall
B
Thalamic Recipient
Thalamus
Integrator
MPall - Integrator
Brain Stem
Long-range output
Canonical trisynaptic circuit
C
Birthdating of chick mesopallial circuitry
HPall
MPall
dNPall
EPall
SPall
APall
PTh
HT
cE15 - SatB2 + Tbr1
chick E15 - SatB2 + EdU
Early - E4
Intermediate - E6
Late - E8
D
F
H
Anterior
E
G
I
Posterior

**Figure S05 – Neurogenesis in the avian Field L takes place at late neurogenic stages.** (**A-C**) E15 chick embryonic brain coronal sections, being (**A**) the most anterior and (**C**) the most posterior, showing the anatomical location of the Field L, identified as a PV-expressing region. (**D-I**) High-power views and split channels of the Field L in coronal sections of E15 chick brains birthdated at E4 (**D. E**), E6 (**F, G**) and E8 (**H, I**). Immunohistochemistry against PV and EdU birthdating reveal that Field L is generated at the end of pallial neurogenesis, by E8. For nomenclature, refer to the list of abbreviations. DAPI counterstain in blue. Dashed white lines demarcate anatomical boundaries. Scale bars, 500 µm (**A-C**), 100 µm (**F, H**), 50 µm (**D**), 20 µm (**E, G, I**) and 5 µm (**E, G, I** insets).

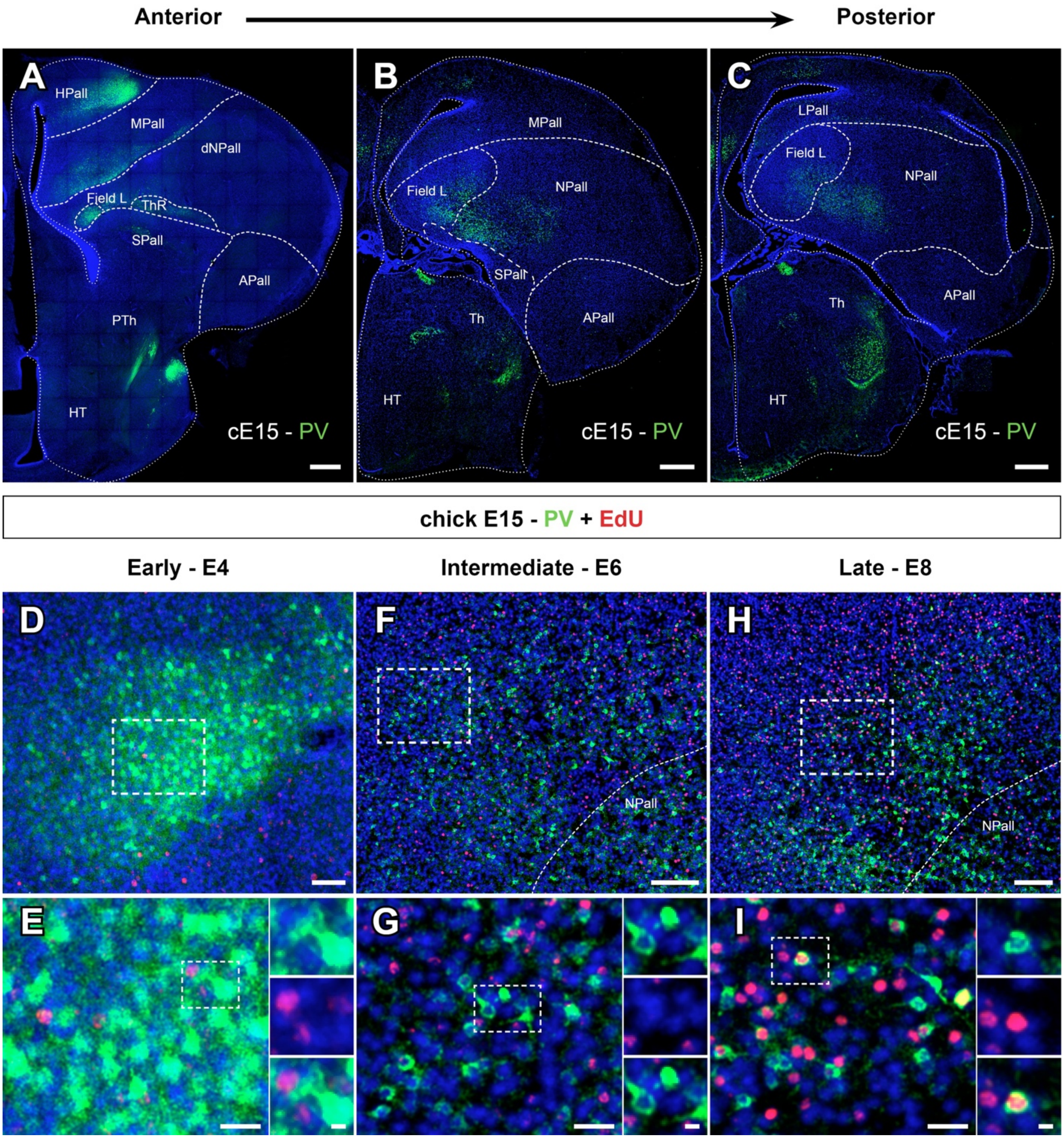


**Figure S06 – Avian mesopallium is generated at intermediate neurogenic timepoints.** (**A**) Representative scheme highlighting the connections of the chick mesopallial sensory processing circuit. (**B**) Scheme of the basic cell type organization in the chick mesopallial circuit. (**C**) Coronal section of E15 chick embryonic brain showing the anatomical location of the mesopallium delimited by the co-expression of SatB2 and Tbr1 markers. (**D-I**) High-power views and split channels of the Field L in anterior (**D, F, H**) and posterior (**E, G, I**) coronal sections of E15 chick brains birthdated at E4 (**D, E**), E6 (**F, G**) and E8 (**H, I**). Immunohistochemistry against SatB2 and EdU birthdating show mesopallial neurons are mostly generated towards the middle of pallial neurogenesis. For nomenclature, refer to the list of abbreviations. DAPI counterstain in blue. Dashed white lines demarcate anatomical boundaries. Scale bars, 1 mm (**C**), 50 µm (**D-I**), and 20 µm (**D-I** insets).

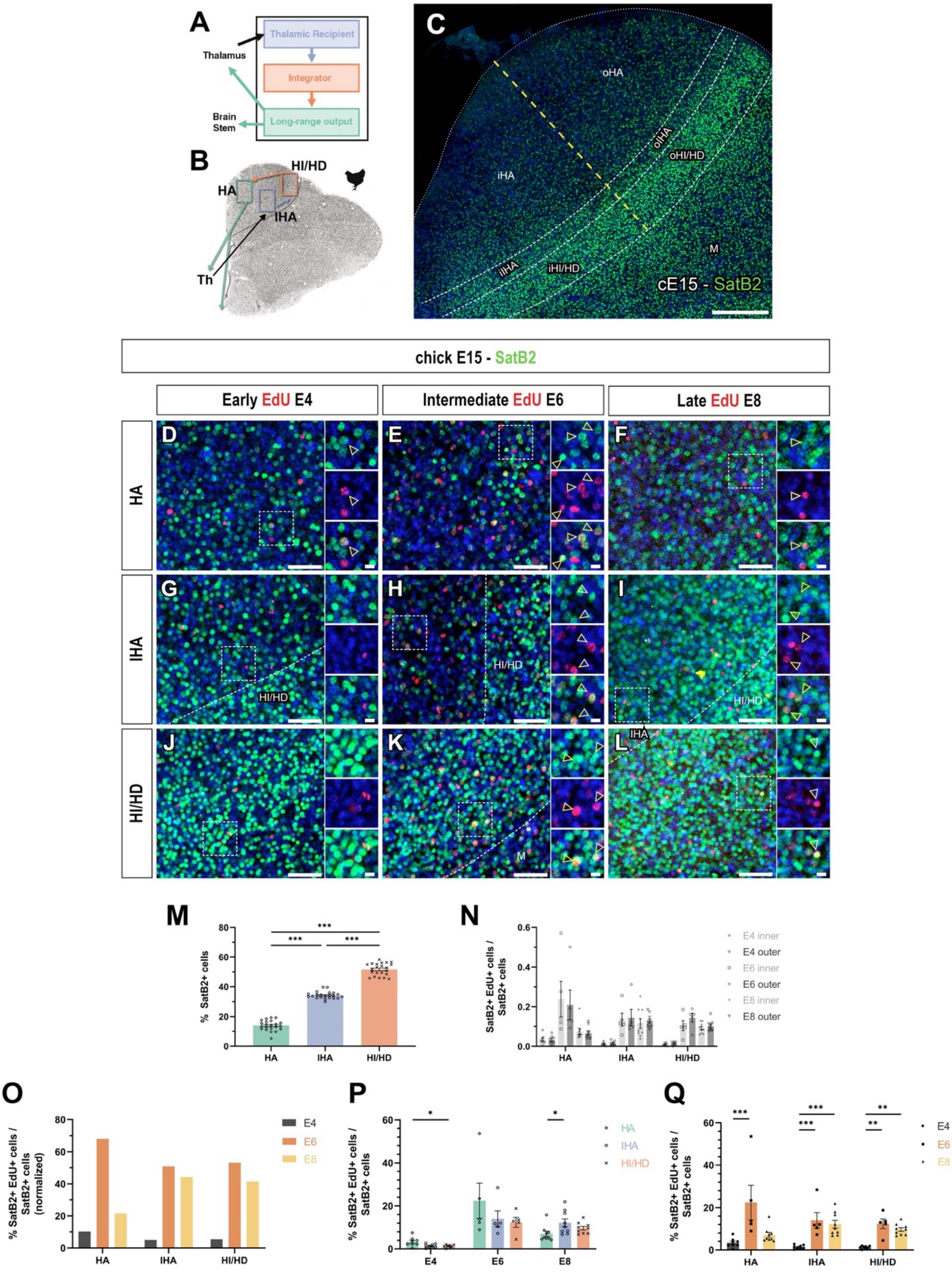


**Figure S07 - SatB2+ and SatB1+ glutamatergic hyperpallial neurons are born during intermediate and late neurogenesis.** (**A**) Representative scheme of the canonical high-order sensory processing trisynaptic circuit. (**B**) Representative diagram of the trisynaptic circuit in the hyperpallium. (**C**) High-power view of the hyperpallium in coronal sections of E15 chick brain showing the distribution pattern of SatB2+ and SatB1glutamatergic neurons. (**D-L**) High-power view and split channels of the HA (**D-F**), IHA (**G-I**) and HI/HD (**J-L**) in coronal sections of E15 chick brains birthdated at E4

(**D, G, J**), E6 (**E, H, K**) and E8 (**F, I, L**). Immunohistochemistry against SatB2 shows that these glutamatergic neurons are generated following the general neurogenic pattern of the hyperpallium. (**M**) Quantification of the distribution of SatB2+ and SatB1 glutamatergic neurons throughout hyperpallial regions. (**N**) Quantification of the differences in the proportion of EdU-labeled SatB2+ and SatB1cells to the total SatB2+ and SatB1 neurons in both inner and outer parts of hyperpallial circuit regions after birthdating at different neurogenic stages. (**O-Q**) Quantification of the percentage of EdU-labeled SatB2+ c and SatB1ells among the total SatB2+ neurons in each of the hyperpallial circuit regions after birthdating at different neurogenic stages. (**O**) Graphic representation of the normalized percentage of neurons of each circuit role generated at early, intermediate or late neurogenic timepoints. (**P**) Graphic representation of hyperpallial circuit cell distribution depending on their birthdate. (**Q**) Graphic representation of hyperpallial circuit cell distribution depending on their birthdate. For nomenclature, refer to the list of abbreviations. DAPI counterstain in blue. Dashed white lines demarcate anatomical boundaries. Dashed yellow line demarcates the inner-outer boundary. Yellow arrowheads indicate representative neurons co-labeled with EdU and the immunohistochemical marker. Scale bars, 250 µm (**C**), 50 µm (**D-L**) and 10 µm (**D-L** insets). *p<0.05; **p<0.01; ***p<0.001. Statistical tests for (**M-Q**) are described in **Supplementary Materials.** Bars represent mean ± SEM. Dots show individual data.

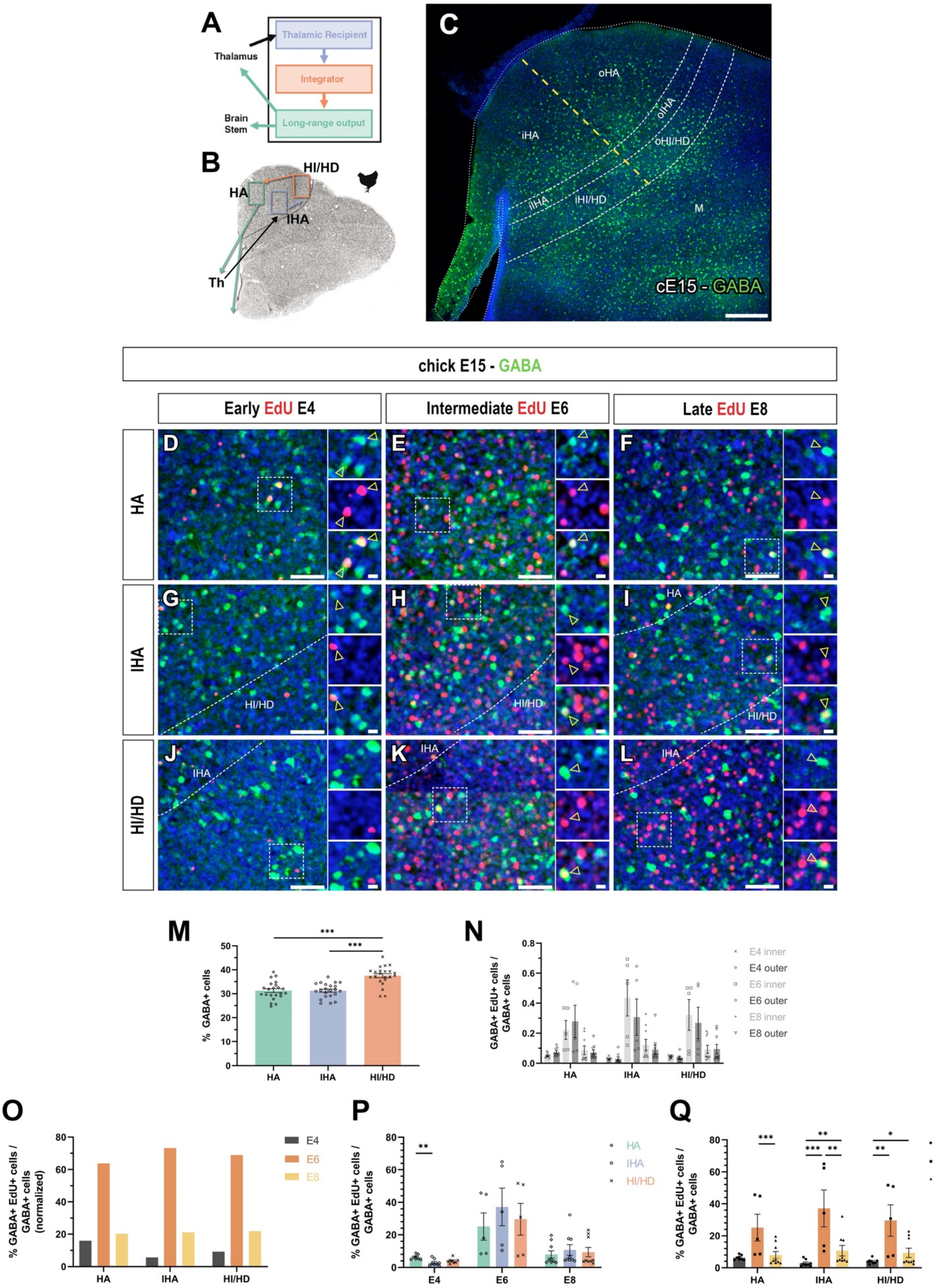


**Figure S08 - Intermediate and late neurogenesis in the chick telencephalon generate GABA+ interneurons of the hyperpallial circuit.** (**A**) Representative scheme of the canonical high-order sensory processing trisynaptic circuit. (**B**) Representative diagram of the trisynaptic circuit in the hyperpallium. (**C**) High-power view of the hyperpallium in coronal sections of E15 chick brain showing the distribution pattern of GABA+ interneurons. (**D-L**) High-power view and split channels of the HA (**D-F**), IHA (**G-I**) and HI/HD (**J-L**) in coronal sections of E15 chick brains birthdated at E4 (**D, G, J**), E6 (**E, H, K**) and E8 (**F, I, L**). Immunohistochemistry against GABA shows that GABAergic interneurons are generated following the general neurogenic pattern of the hyperpallium. (**M**) Quantification of the distribution of GABA+ interneurons throughout hyperpallial regions. (**N**) Quantification of the differences in the proportion of EdU-labeled GABA+ cells to the total GABA+ neurons in both inner and outer parts of hyperpallial circuit regions after birthdating at different neurogenic stages. (**O-Q**) Quantification of the percentage of EdU-labeled GABA+ cells among the total GABA+ neurons in each of the hyperpallial circuit regions after birthdating at different neurogenic stages. (**O**) Graphic representation of the normalized percentage of neurons of each circuit role generated at early, intermediate or late neurogenic timepoints. (**P**) Graphic representation of the effect of age in the generation hyperpallial circuit regions. (**Q**) Graphic representation of hyperpallial circuit cell distribution depending on their birthdate. For nomenclature, refer to the list of abbreviations. DAPI counterstain in blue. Dashed white lines demarcate anatomical boundaries. Dashed yellow line demarcates the inner-outer boundary. Yellow arrowheads indicate representative neurons co-labeled with EdU and the immunohistochemical marker. Scale bars, 250 µm (**C**), 50 µm (**D-L**) and 10 µm (**D-L** insets). *$p<0.05$; **$p<0.01$; ***$p<0.001$. Statistical tests for (**M-Q**) are described in **Supplementary Materials.** Bars represent mean ± SEM. Dots show individual data.

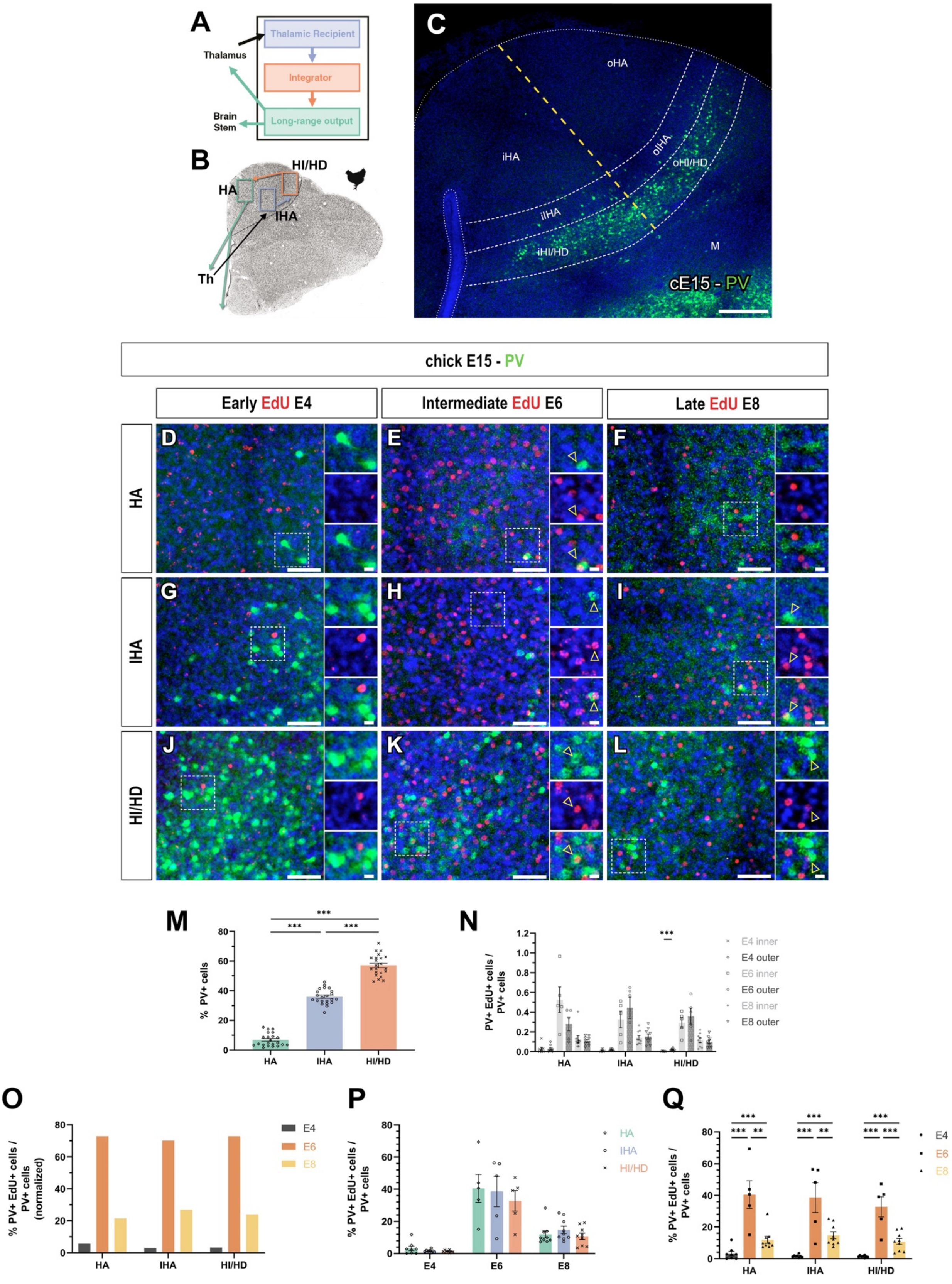


**Figure S09 - Intermediate and late neurogenesis in the chick telencephalon generate PV+ GABAergic interneurons of the hyperpallial circuit.** (**A**) Representative scheme of the canonical high-order sensory processing trisynaptic circuit. (**B**) Representative diagram of the trisynaptic circuit in the hyperpallium. (**C**) High-power view of the hyperpallium in coronal sections of E15 chick brain showing the distribution pattern of PV+ GABAergic interneurons. (**D-L**) High-power view and split channels of the HA (**D-F**), IHA (**G-I**) and HI/HD (**J-L**) in coronal sections of E15 chick brains birthdated at E4 (**D, G,**

**J**), E6 (**E, H, K**) and E8 (**F, I, L**). Immunohistochemistry against PV shows that these glutamatergic neurons are generated following the general neurogenic pattern of the hyperpallium. (**M**) Quantification of the distribution of PV+ GABAergic interneurons throughout hyperpallial regions. (**N**) Quantification of the differences in the proportion of EdU-labeled PV+ cells to the total PV+ GABAergic interneurons in both inner and outer parts of hyperpallial circuit regions after birthdating at different neurogenic stages. (**O-Q**) Quantification of the percentage of EdU-labeled PV+ cells among the total PV+ interneurons in each of the hyperpallial circuit regions after birthdating at different neurogenic stages. (**O**) Graphic representation of the normalized percentage of neurons of each circuit role generated at early, intermediate or late neurogenic timepoints. (**P**) Graphic representation of the effect of age in the generation hyperpallial circuit regions. (**Q**) Graphic representation of hyperpallial circuit cell distribution depending on their birthdate. For nomenclature, refer to the list of abbreviations. DAPI counterstain in blue. Dashed white lines demarcate anatomical boundaries. Dashed yellow line demarcates the inner-outer boundary. Yellow arrowheads indicate representative neurons co-labeled with EdU and the immunohistochemical marker. Scale bars, 250 µm (**C**), 50 µm (**D-L**) and 10 µm (**D-L** insets). $^{**}p<0.01$; $^{***}p<0.001$.Statistical tests for (**M-Q**) are described in **Supplementary Materials.** Bars represent mean ± SEM. Dots show individual data.

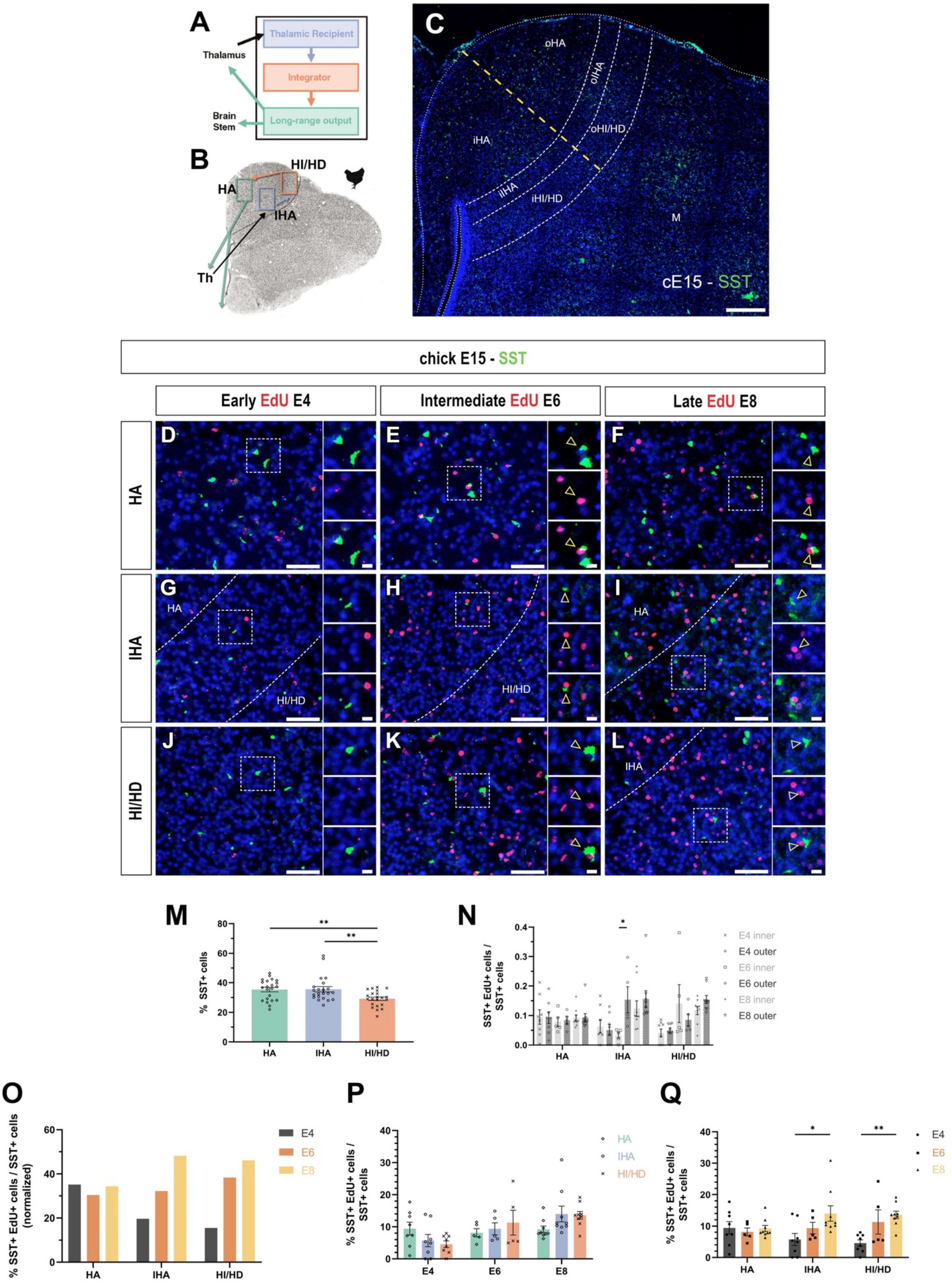


**Figure S10 - SST+ GABAergic interneurons of the hyperpallium are born throughout the whole telencephalic neurogenesis process.** (**A**) Representative scheme of the canonical high-order sensory processing trisynaptic circuit. (**B**) Representative diagram of the trisynaptic circuit in the hyperpallium. (**C**) High-power view of the hyperpallium in coronal sections of E15 chick brain showing the distribution pattern of SST+ GABAergic interneurons. (**D-L**) High-power view and split channels of the HA (**D-F**), IHA (**G-I**) and HI/HD (**J-L**) in coronal sections of E15 chick brains birthdated at E4 (**D, G,**

**J**), E6 (**E, H, K**) and E8 (**F, I, L**). Immunohistochemistry against SST shows that these GABAergic interneurons preferentially generated at intermediate and late neurogenic stages. (**M**) Quantification of the distribution of SST+ GABAergic interneurons throughout hyperpallial regions. (**N**) Quantification of the differences in the proportion of EdU-labeled SST+ cells to the total SST+ interneurons in both inner and outer parts of hyperpallial circuit regions after birthdating at different neurogenic stages. (**O-Q**) Quantification of the percentage of EdU-labeled SST+ cells among the total SST+ GABAergic interneurons in each of the hyperpallial circuit regions after birthdating at different neurogenic stages. (**O**) Graphic representation of the normalized percentage of neurons of each circuit role generated at early, intermediate or late neurogenic timepoints. (**P**) Graphic representation of the effect of age in the generation hyperpallial circuit regions. (**Q**) Graphic representation of hyperpallial circuit cell distribution depending on their birthdate. For nomenclature, refer to the list of abbreviations. DAPI counterstain in blue. Dashed white lines demarcate anatomical boundaries. Dashed yellow line demarcates the inner-outer boundary. Yellow arrowheads indicate representative neurons co-labeled with EdU and the immunohistochemical marker. Scale bars, 250 µm (**C**), 50 µm (**D-L**) and 10 µm (**D-L** insets). *p<0.05; **p<0.01. Statistical tests for (**M-Q**) are described in **Supplementary Materials.** Bars represent mean ± SEM. Dots show individual data.

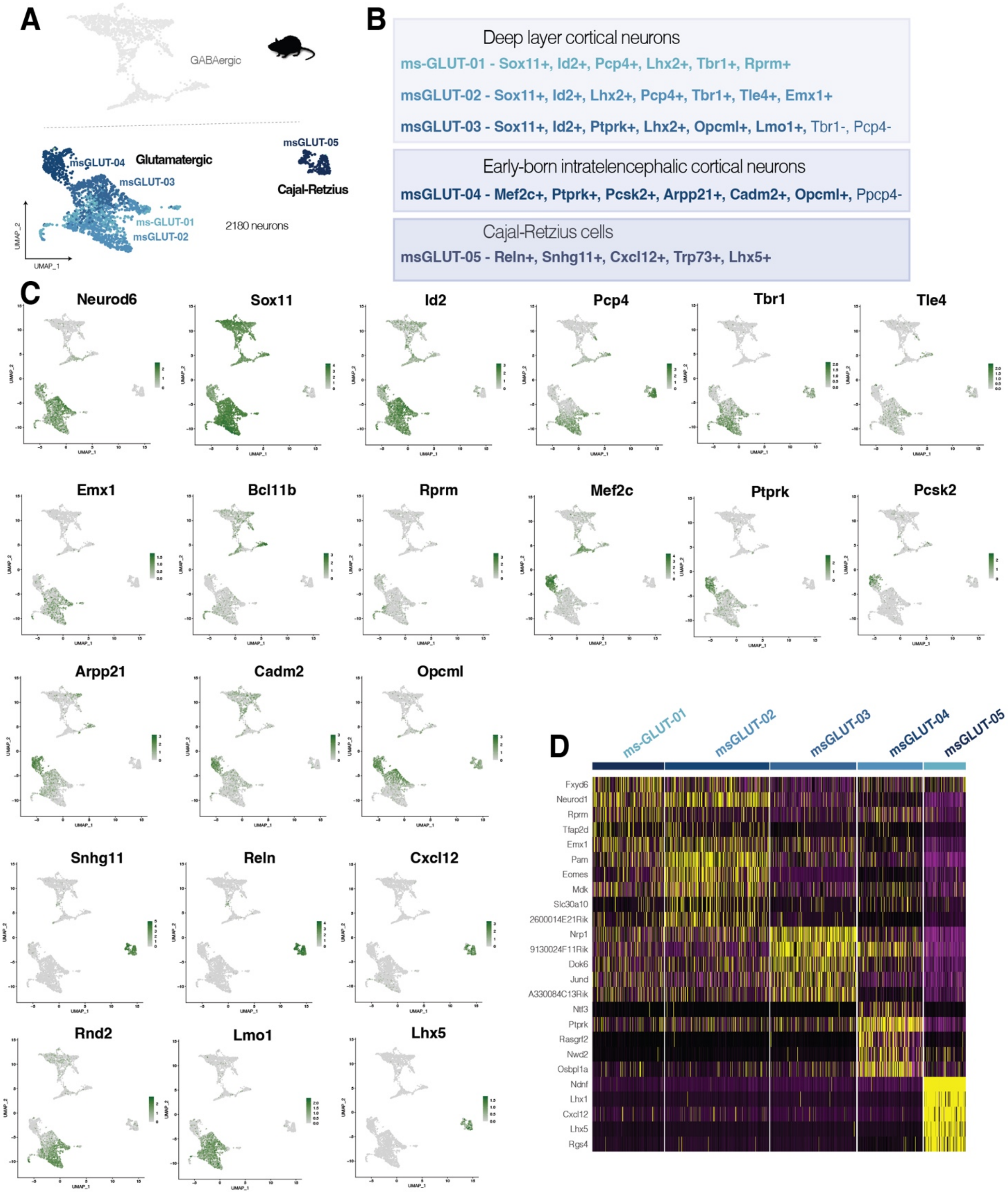


**Figure S11– Transcriptomic profile of early born glutamatergic neurons of the mouse pallium at maturing timepoint.** EdU was injected at E12 and neurons were FAC-sorted and sequenced at P3. (**A**) UMAP representation of the neurons and clusters of E12-generated neurons. Clusters analyzed in this figure comprised glutamatergic cells, including Cajal-Retizus cells. (**B**) Identification of glutamatergic neurons according to expression of gene markers. (**C**) UMAP distribution of cells expressing the differentially expressed genes for each cluster. (**D**) Heatmap to represent the gene expression of E12-generated glutamatergic neurons of the mouse pallium.

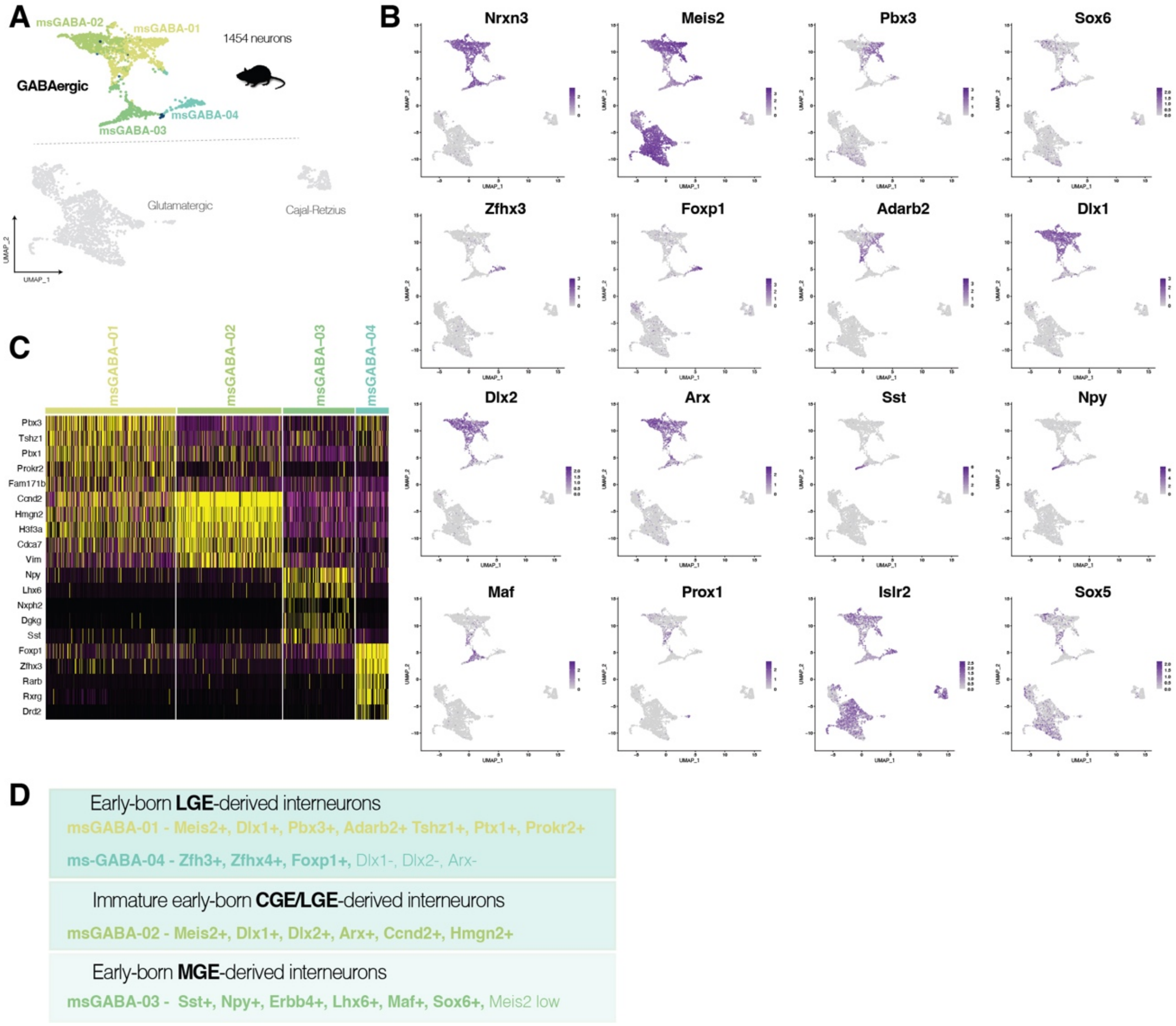


**Figure S12– Transcriptomic profile of early born GABAergic neurons of the mouse pallium at maturing timepoint.** EdU was injected at E12 and neurons were FAC-sorted and sequenced at P3. (**A**) UMAP representation of the neurons and clusters of E12-generated neurons. Clusters analyzed in this figure comprised GABAergic interneurons only. (**B**) UMAP distribution of cells expressing the differentially expressed genes for each cluster. (**C**) Heatmap to represent the gene expression of E12-generated glutamatergic neurons of the mouse pallium. (**D**) Identification of GABAergic neurons according to expression of gene markers.

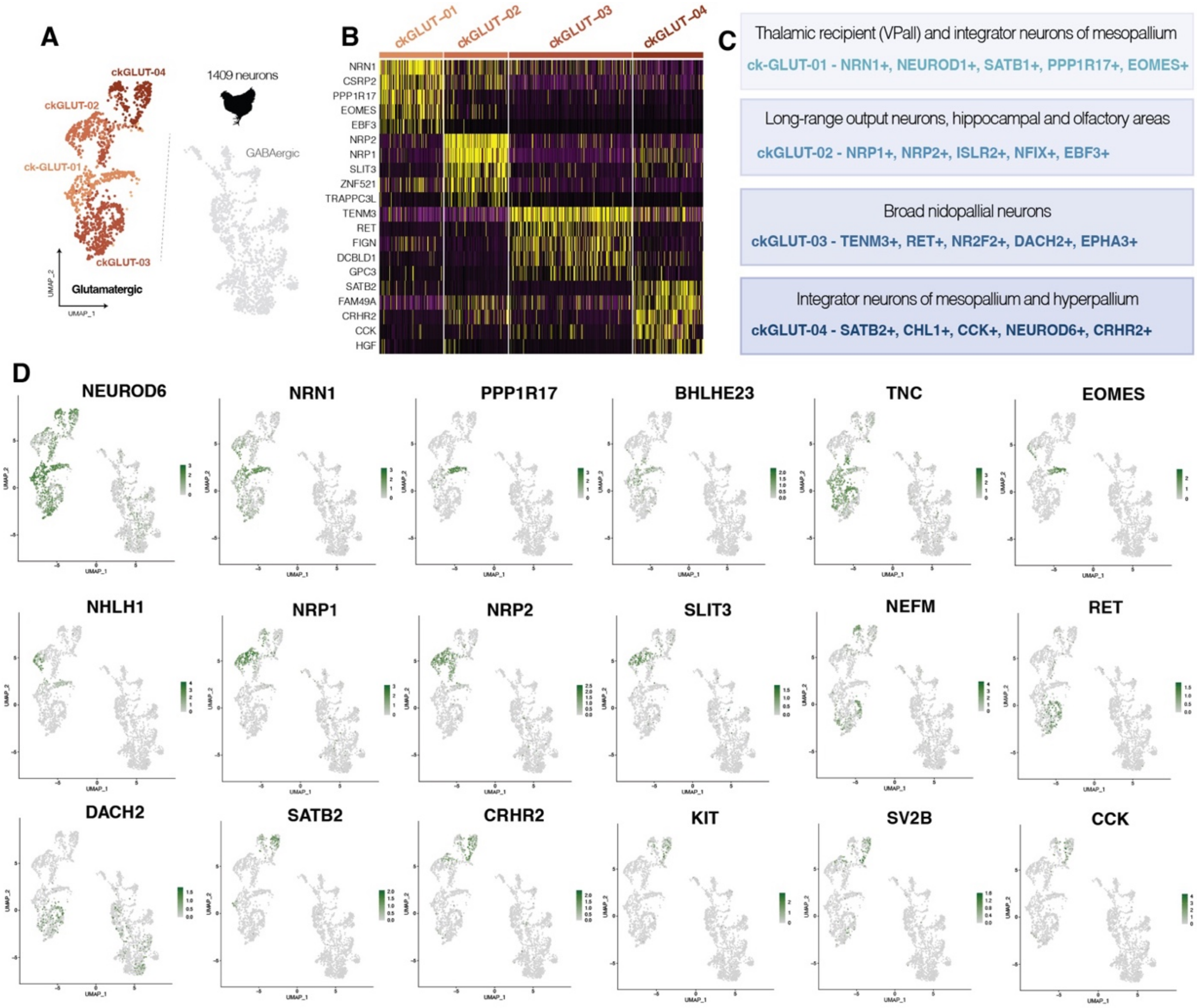


**Figure S13– Transcriptomic profile of early born glutamatergic neurons of the chick pallium at maturing timepoint.** EdU was injected at E4 and neurons were FAC-sorted and sequenced at E15. (**A**) UMAP representation of the neurons and clusters of E4-generated neurons. Clusters analyzed in this figure comprised glutamatergic neurons only. (**B**) Heatmap to represent the gene expression of E4-generated glutamatergic neurons of the chick pallium. (**C**) Identification of glutamatergic neurons according to expression of gene markers and aided by the ISS experiments of figures **5** and **S20-S22**. (**D**) UMAP distribution of cells expressing the differentially expressed genes for each cluster.

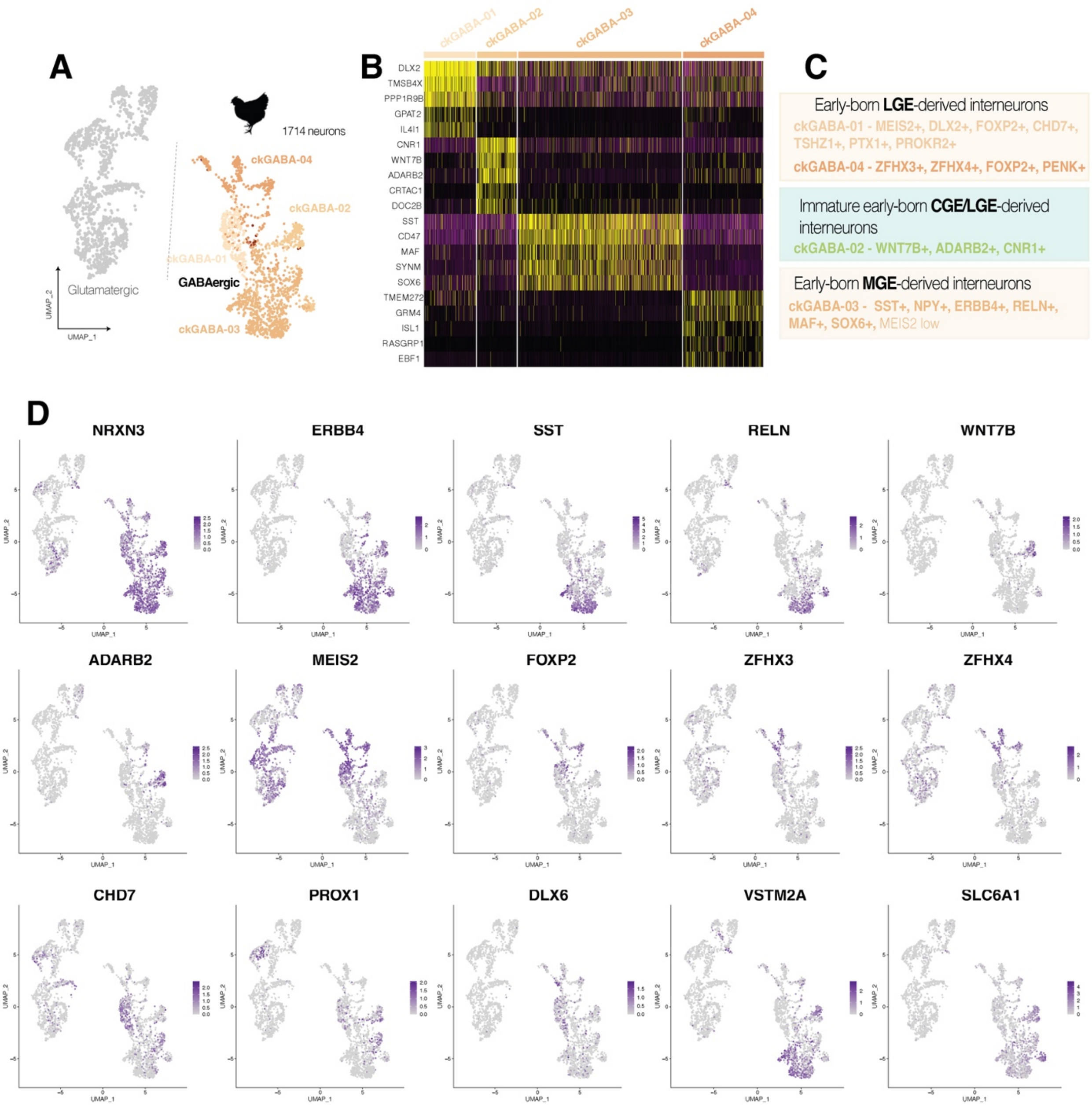


**Figure S14– Transcriptomic profile of early born GABAergic neurons of the chick pallium at maturing timepoint.** EdU was injected at E4 and neurons were FAC-sorted and sequenced at E15. (**A**) UMAP representation of the neurons and clusters of E4-generated neurons. Clusters analyzed in this figure comprised GABAergic neurons only. (**B**) Heatmap to represent the gene expression of E4-generated GABAergic neurons of the chick pallium. (**C**) Identification of GABAergic neurons according to expression of gene markers and aided by the ISS experiments of figures **5** and **S20-S22**. (**D**) UMAP distribution of cells expressing the differentially expressed genes for each cluster.

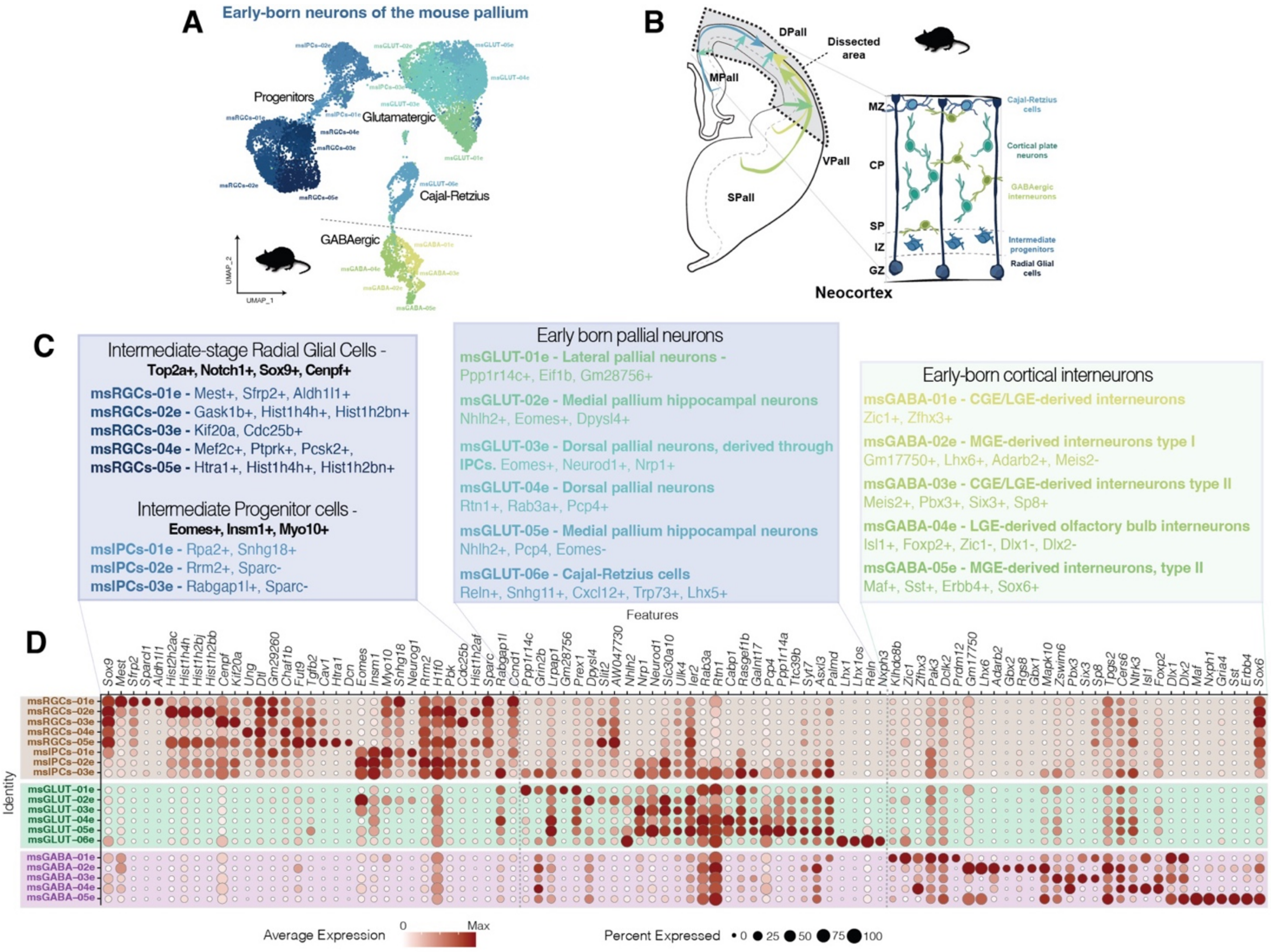


**Figure S15– Transcriptomic profile of early born neurons of the mouse pallium early after neurogenesis.** EdU was injected at E12 and neurons were FAC-sorted and sequenced two days later, at E14. (**A**) UMAP representation of the neurons and clusters of E12-generated cells. Clusters -labeled with an *e*, standing for embryonic- analyzed in this figure comprised radial glial cells, intermediate precursor cells and neurons of both, glutamatergic and GABAergic lineage. (**B**) Proposed location of the cells of each cluster in the developing pallium of the mouse. Two schemes depict a coronal section of the E14 telencephalon and a close-up view of the pallial neuroepithelium (**C**) Identification of glutamatergic neurons according to expression of gene markers. (**D**) Dot plot of the differentially expressed genes for each cluster, showing the percentage of cells of the cluster and the average level of expression. Abbreviations: CP, cortical plate; IZ. intermediate zone; GZ, germinal zone; MZ, marginal zone; SP, subplate.

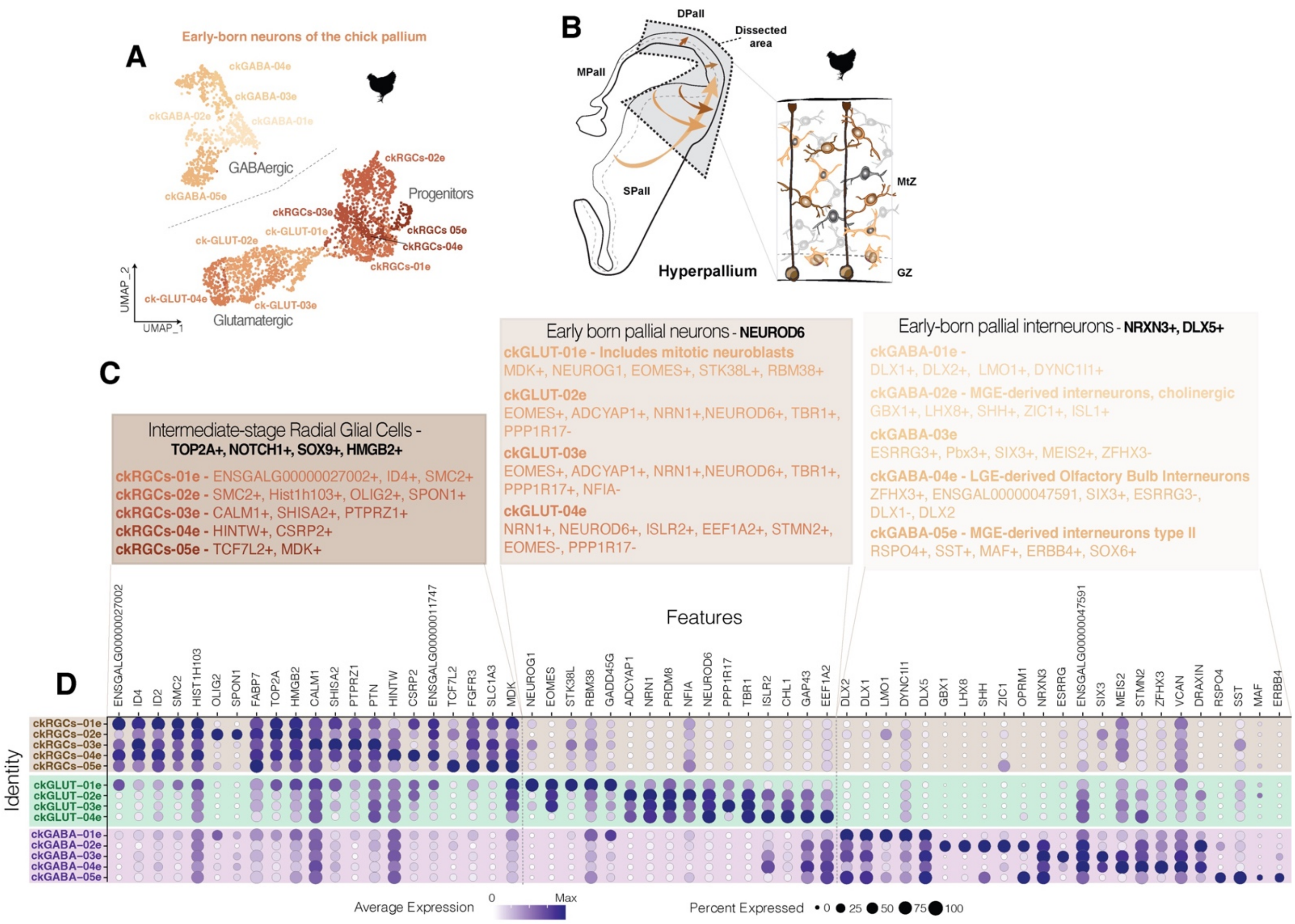


**Figure S16– Transcriptomic profile of early born neurons of the chick pallium early after neurogenesis.** EdU was injected at E4 and neurons were FAC-sorted and sequenced two days later, at E6. (**A**) UMAP representation of the neurons and clusters of E4-generated cells. Clusters -labeled with an *e*, standing for embryonic- analyzed in this figure comprised radial glial cells and neurons of both, glutamatergic and GABAergic lineage. (**B**) Proposed location of the cells of each cluster in the developing pallium of the chick. Two schemes depict a coronal section of the E14 telencephalon and a close-up view of the pallial neuroepithelium (**C**) Identification of glutamatergic neurons according to expression of gene markers. (**D**) Dot plot of the differentially expressed genes for each cluster, showing the percentage of cells of the cluster and the average level of expression.

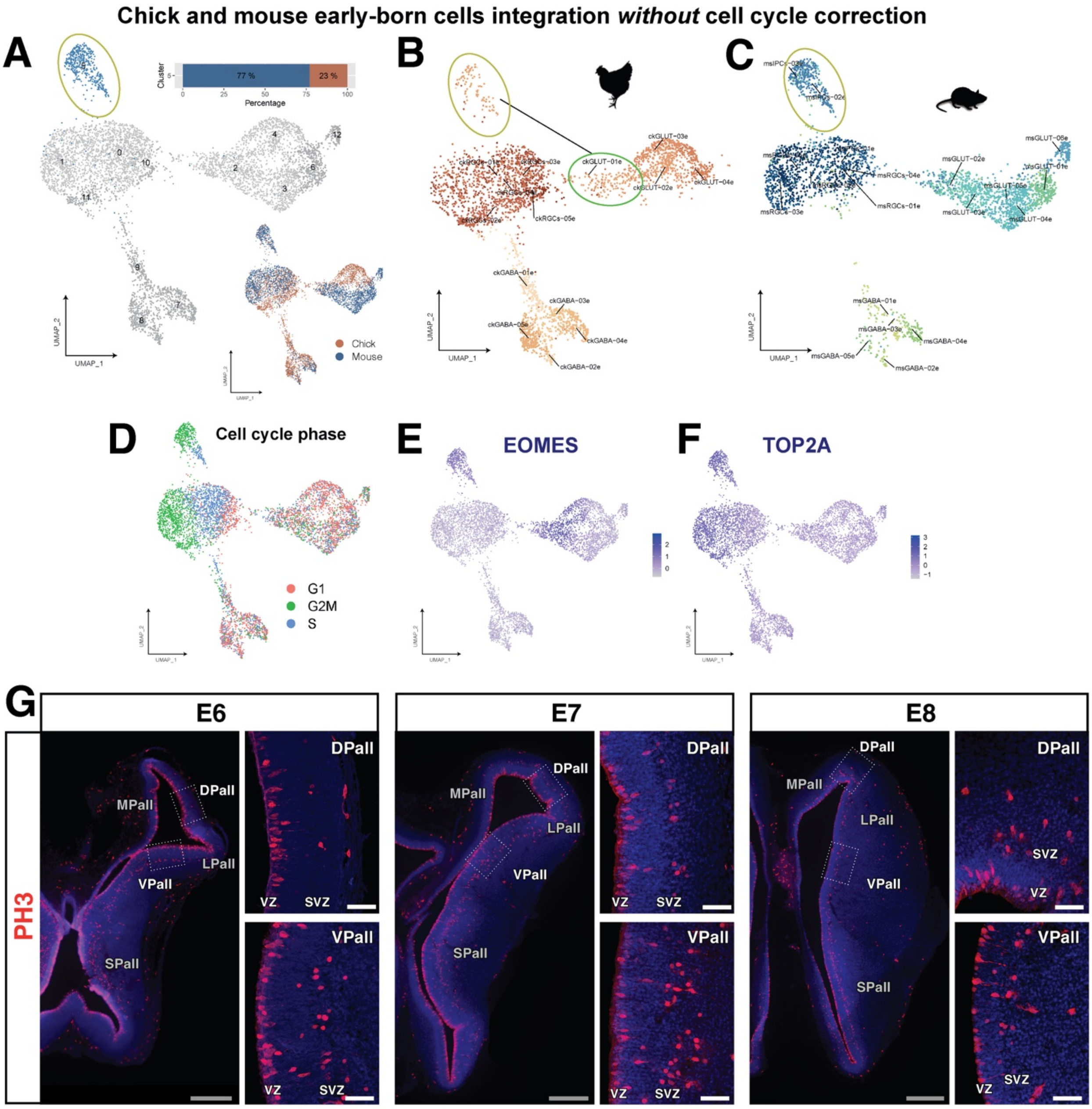


**Figure S17– Transcriptomic similarities and differences of mouse and chick IPCs.** Integration of chicken and mouse day 1 datasets without cell cycle regression (**A-F**). UMAP with newly integrated IPCs cluster with the species distribution in bar percentage and UMAP (**A**). Subset of chicken (**B**) and mouse (**C**) cells in the integrated UMAP with the original annotations. Other features of the integrated UMAP such as the cell cycle (D), IPCs markers like EOMES (**E**) and proliferation markers like TOP2A (**F**) showing that ckGLUT-01e cluster can be split into a mitotic subcluster and a postmitotic subcluster (golden circle and green circle respectively, in **B**). (**G**) Proliferative cells in the SVZ of chicken brain as revealed by PH3 immunohistochemistry. Images taken at three developmental stages (E6, E7 and E8), presenting a persistent subventricular zone in the VPall of chick telencephalon in opposition to low number of abventricular mitotic cells in the DPall germinative zone. DAPI counterstain in blue. Colored scale bars: grey, 500 µm; white, 50 µm.

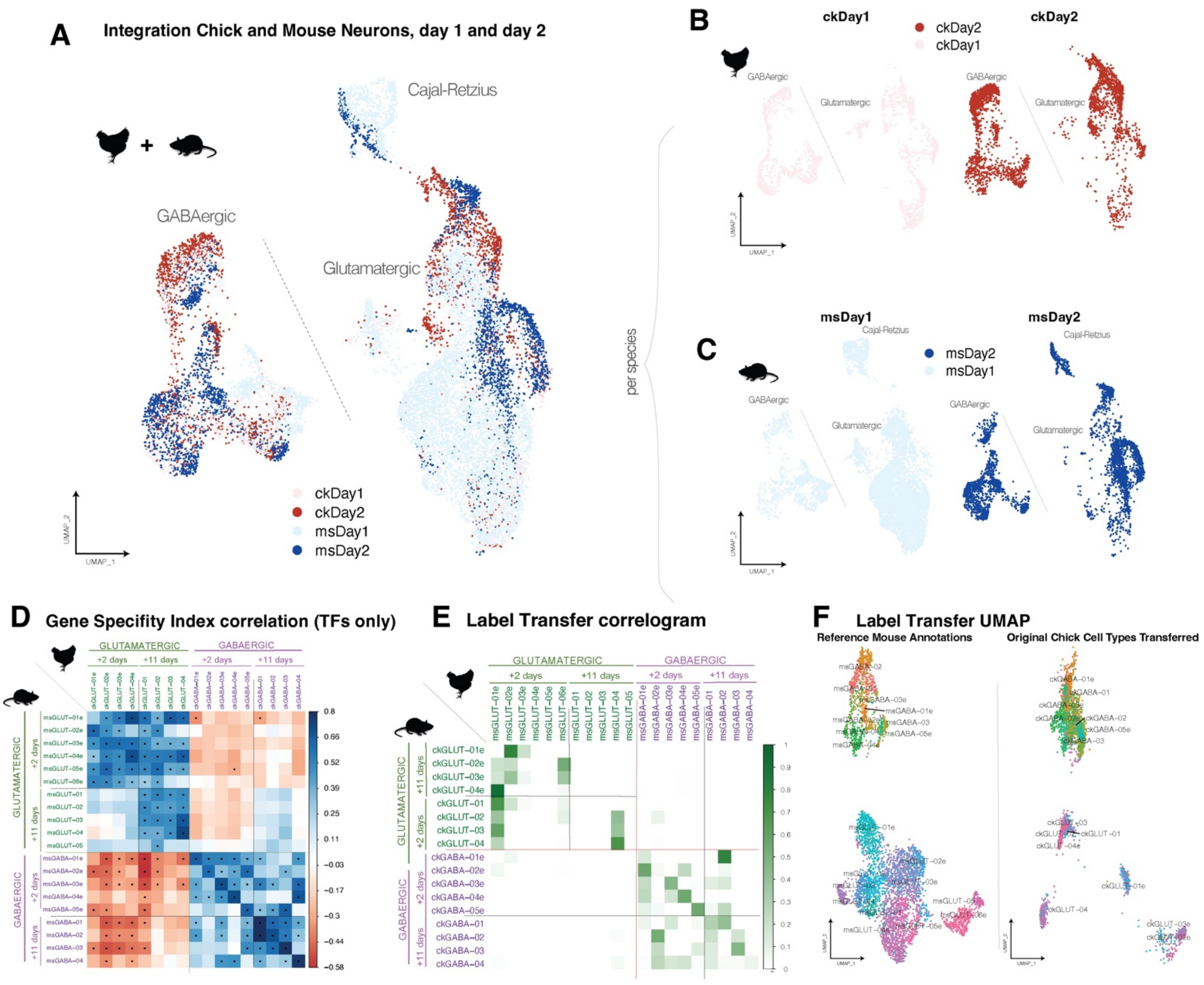


**Figure S18 – Extended transcriptomic comparison of early born pallial neurons.** (**A**) UMAP graph of the CCA integration of all early born pallial neurons in the datasets, two for each species: one of cells early after neurogenesis (*day 1*; cells sequenced 2 days after EdU birthdating, pale color) and another one at early maturation timepoint (*day 2*; 11 days after EdU birthdating, dark color). Chick datasets in pink-red, mouse datasets in blue. (**B,C**) UMAP representation of the datasets in the integration separated by species and stages. (**D**) Correlogram of gene specificity indexes between pairs of main cell types of the two species considering the expression of TFs only; clusters from the day 1 sample are labeled with an *e*, standing for embryonic. (**E,F**) Label transfer analysis of all mouse neurons onto the mouse annotated reference (**E**) Proportion of cells from a mouse cluster that were assigned to a given chick cluster. (**F**) UMAP representation of the label transfer analysis.

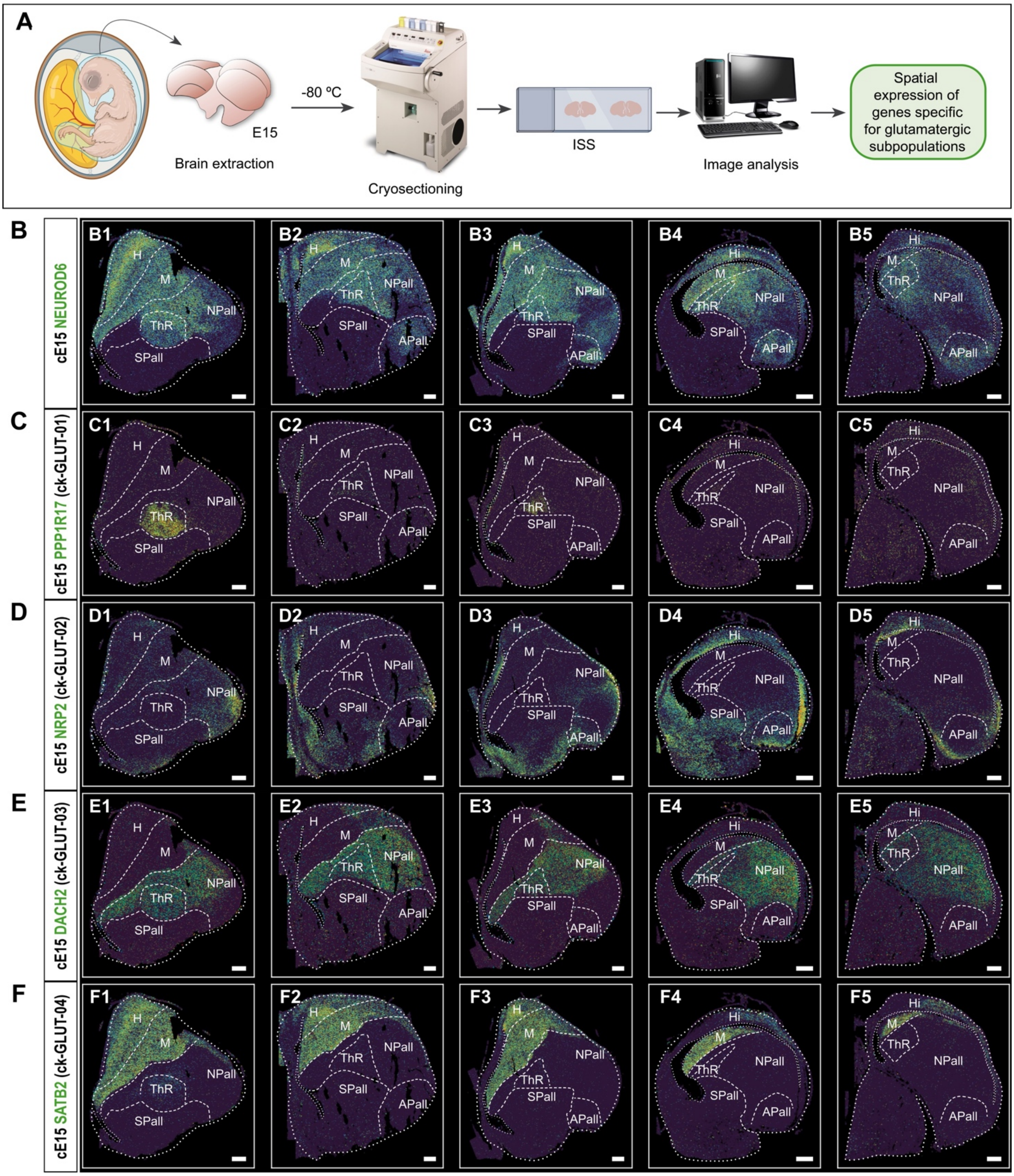


**Figure S19 – Expression pattern of selected glutamatergic gene markers by ISS on the chick E15 telencephalon.** (**A**) Schematic representation of the experimental design of ISS for the spatial visualization of genes specific for the glutamatergic neuron clusters defined in the scRNA-Seq analysis of early-born mature chick neurons. (**B-F**) Five brain coronal sections of E15 chick embryos from anterior to posterior with in situ staining for population-specific *NEUROD6* (**B**), *PPP1R17* (**C**), *NRP2* (D), *DACH2* (**E**) and *SATB2* (**F**) showing their expression. For nomenclature, refer to the list of abbreviations. Dashed white lines demarcate anatomical boundaries. Scale bars, 500 µm. Sections were obtained from 3 different embryos.

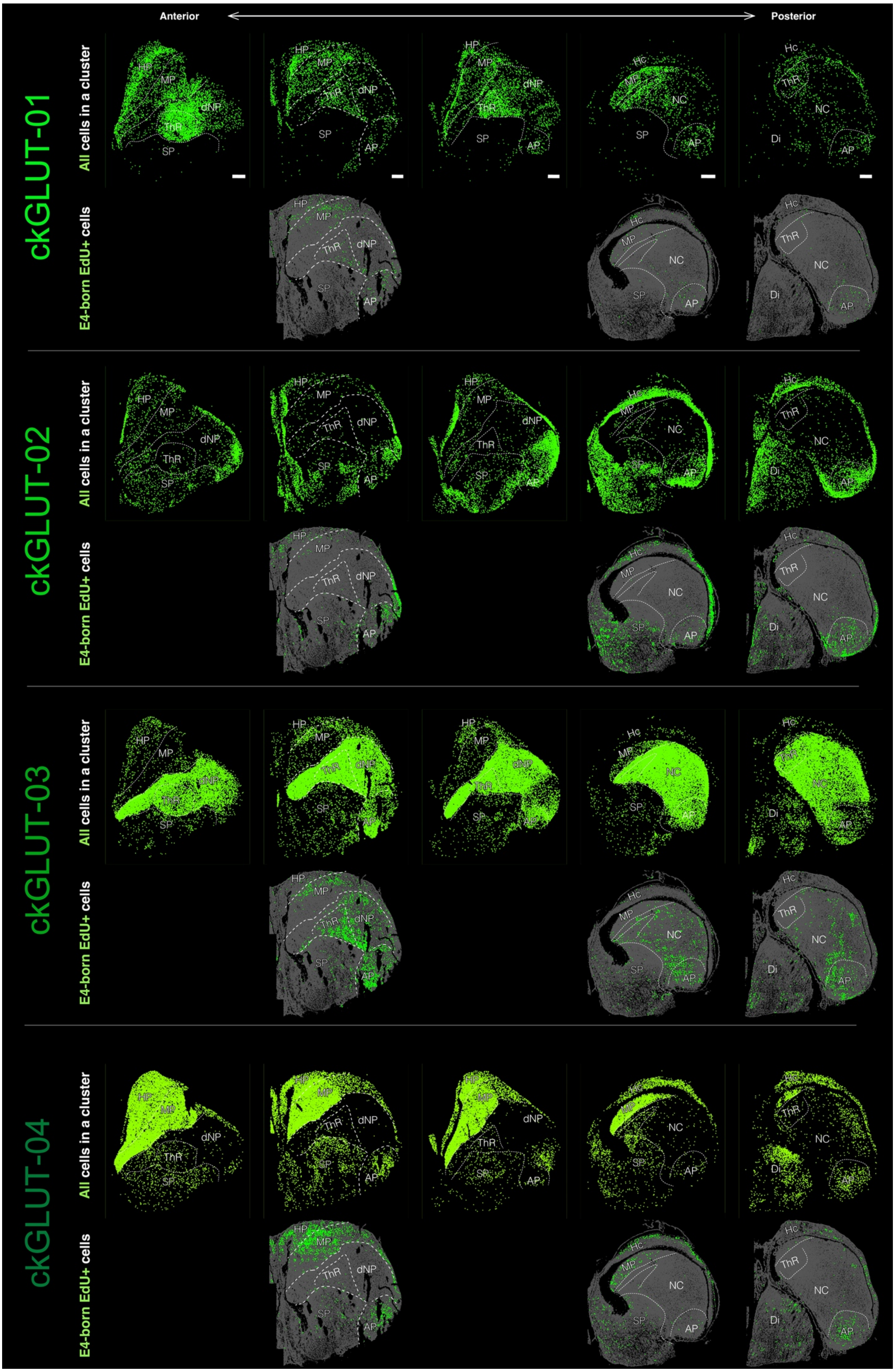
Anterior
Posterior
ckGLUT-01
ckGLUT-02
ckGLUT-03
ckGLUT-04
All cells in a cluster
E4-born EdU+ cells
HP
MP
dNP
ThR
SP
AP
Hc
NC
Di

**Figure S20 – Pallial distribution of early-born pallial glutamatergic neurons as revealed by *Neurogenes-ISS*.** For each of the four glutamatergic clusters identified by scRNAseq (ckGLUT01-04), the top row shows the distribution of cells assigned to that given clusters after ISS of a panel of 84 genes and probabilistic cell typing. Five coronal sections are shown covering the anterior (left) to posterior (right) levels of the pallium. The bottom row displays the Neurogenes-iss analysis of early-born glutamatergic neurons assigned to a given cluster after E4 birthdating with EdU. Three sections are shown corresponding to the animals that received the E4 EdU administration. Scale bars, 500 µm. Sections were obtained from 3 different embryos.

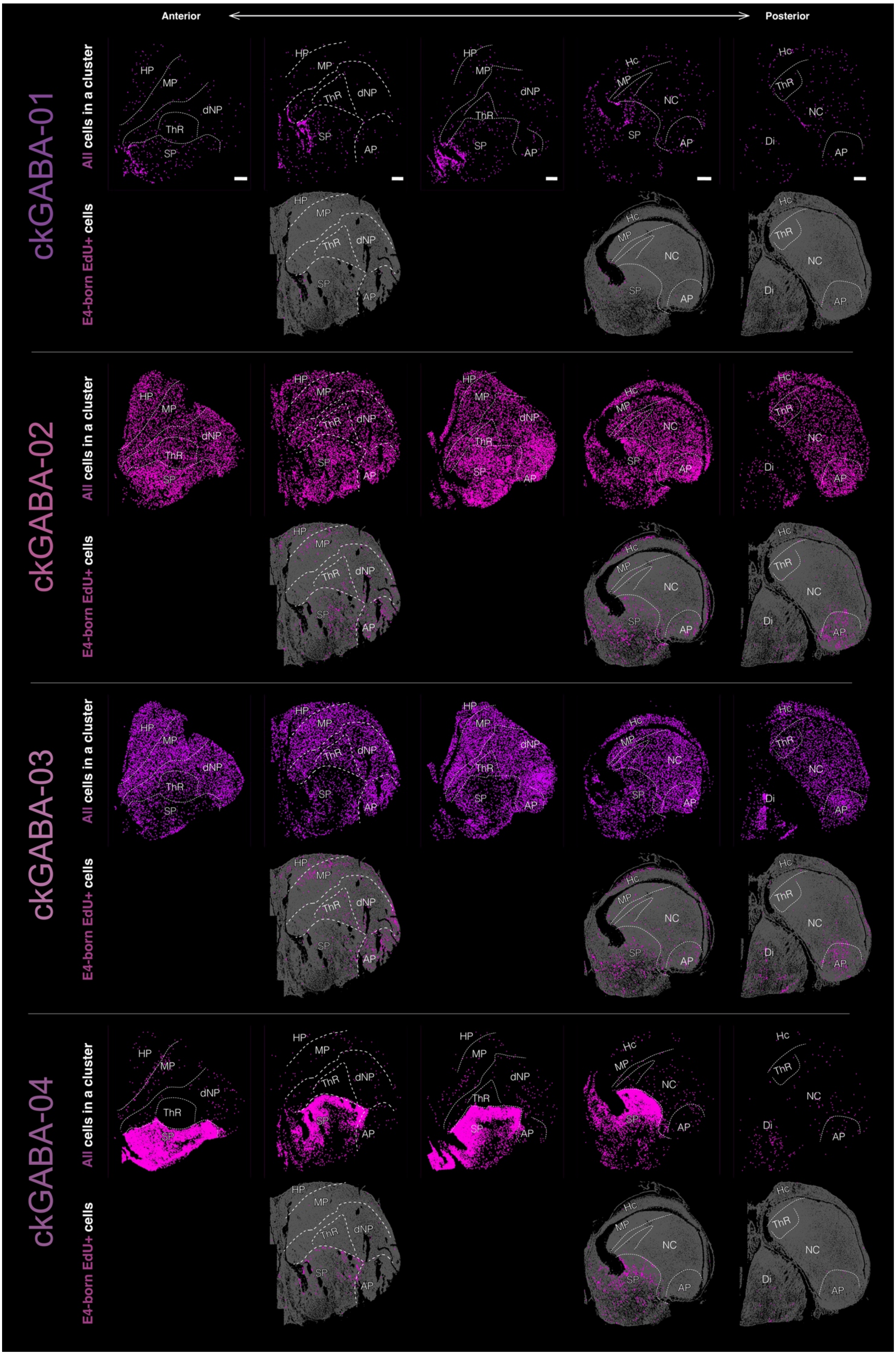
Anterior
Posterior
ckGABA-01
ckGABA-02
ckGABA-03
ckGABA-04
All cells in a cluster
E4-born EdU+ cells
HP
MP
dNP
ThR
SP
AP
Hc
NC
Di

**Figure S21 – Pallial distribution of early-born pallial GABAergic neurons as revealed by *Neurogenes-ISS*.** For each of the four GABAergic clusters identified by scRNAseq (ckGABA01-04), the top row shows the distribution of cells assigned to that given clusters after ISS of a panel of 84 genes and probabilistic cell typing. Five coronal sections are shown covering the anterior (left) to posterior (right) levels of the pallium. The bottom row displays the Neurogenes-iss analysis of early-born glutamatergic neurons assigned to a given cluster after E4 birthdating with EdU. Three sections are shown corresponding to the animals that received the E4 EdU administration. Scale bars, 500 µm. Sections were obtained from 3 different embryos.

---

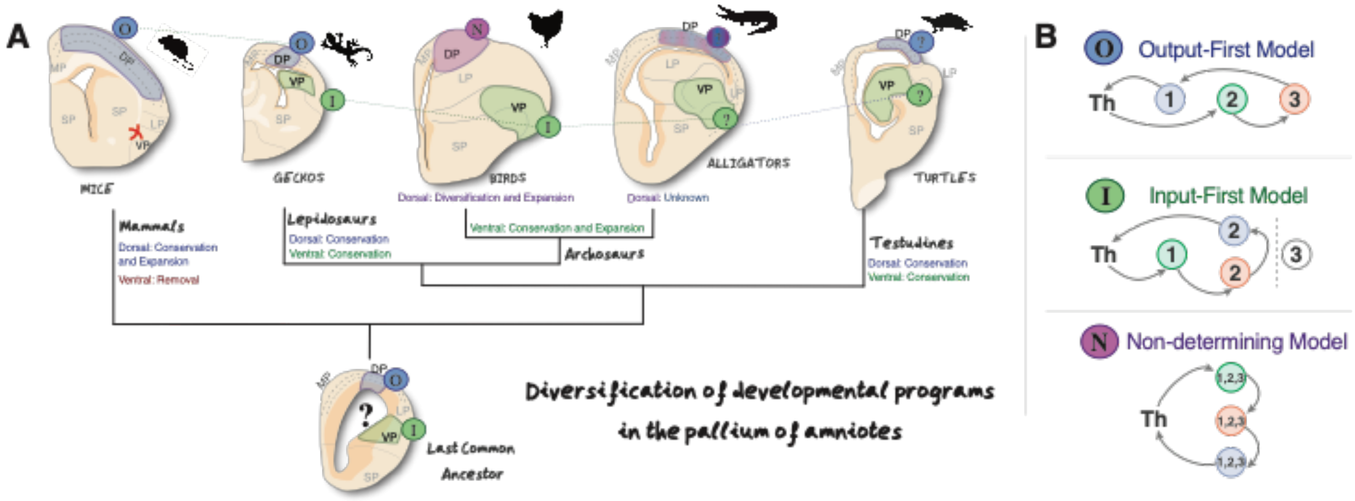


**Figure S22 – Diversification of developmental programs in the pallium of amniotes.** (**A**). Phylogenic diagram depicting the proposed evolutionary scenario of pallial sensory circuits in amniotes. The diagram represents a parsimonious scenario on which the last common ancestor of amniotes possessed two separated sensory circuits in the pallium, which were different and according to their developmental program. Lepidosaurian reptiles inherited these circuits. Avian archosaurs inherited and expanded the ventral pallial circuit, whereas highly evolved a new dorsal pallial circuit. On the other hand, mammals inherited and expanded the dorsal pallial circuitry, whereas completely ablated the ventral circuit. Non-avian archosaurs and testudine reptiles -not investigated in this research- are shown in the diagram with their speculative scenarios of pallial circuit development, according to available literature and the most parsimonious scenario. Amongst them, the most open scenario are non-avian archosaurs, that possess both a laminated cortex in DPall such as other reptiles and mammals, and an avian-like DVR in the VPall. (**B**) The sequential order of generation of each neuronal type in the sensory pallium. *O* from Output first, *I* for Input-first and *N* for Non-determining models are marked in the diagrams of (**A**).
