## Supplementary notes on the maths model for "Evolutionary convergence of sensory circuits in the pallium of amniotes"

### Supplementary Notes S1 for

#### **Evolutionary convergence of sensory circuits in the pallium of amniotes**

Eneritz Rueda-Alaña et al,

### 1 Characterization of neurogenesis

#### 1.1 Experimental rates of neurogenesis

Neurogenesis is characterized by the proportion of neurons per brain area that are generated at each development stage. In this framework, development stages are expressed by embryon stages (E#), sampled every two days. Neurogenesis of the circuits that we are interest in lasts about ten days in both the chick and the mouse, although in the latter, it starts at later stages. Tables 1 and 2 summarize the experimental data available by this study for the chick, and for the mouse by Hatanaka et al. (2016).

| *Region* | *E2* | *E4* | *E6* | *E8* |
|---|---|---|---|---|
| dNPall | 0% | 9.42% | 85.45% | 5.13% |
| EPall | 0% | 80.00% | 16.78% | 3.22% |
| APall | 0% | 50.64% | 41.04% | 8.32% |

**Table 1.** Proportion of neurons generated at each development stage in each circuit relay in the chick. Data from original experiments.

| *Region* | *E9.5* | *E11.5* | *E13.5* | *E15.5* |
|---|---|---|---|---|
| L2/3 | 0% | 1.6 % | 33.3 % | 65.1 % |
| L4 | 0% | 5.5 % | 91.2 % | 3.3 % |
| L5/6 | 0% | 73.23% | 25.24% | 1.53% |

**Table 2.** Proportion of neurons generated at each development stage in each circuit relay in the mouse. Data by Hatanaka et al. (2016).

The proportions of neurons per region sum up to 100% by end (i.e. completion of neurogenesis). Hence the data in the tables can be conveniently plotted (Figure 1) in terms of percentage of completion ($N\%$) of neurogenesis as a function of the development stage, where the percentage of completion of neurogenesis at the $i$-th stage is simply $N\%_i = \sum_{0 \leq j < i} N\%_j$. We note that the percentages are meant to match the time instant at the upper bound of each embryonic stage. In other words, $N\%_i$ corresponds to the time instant immediately preceding $E_{i+i}$.

#### 1.2 Model of neurogenesis

Dividing the percentage of neurogenesis completion by 100 provides the region-specific *fraction* of the relay circuit developed up to some stage, in terms of fraction of total neurons present in that circuit at a given point in time. On the other hand, such fraction can also be interpreted as

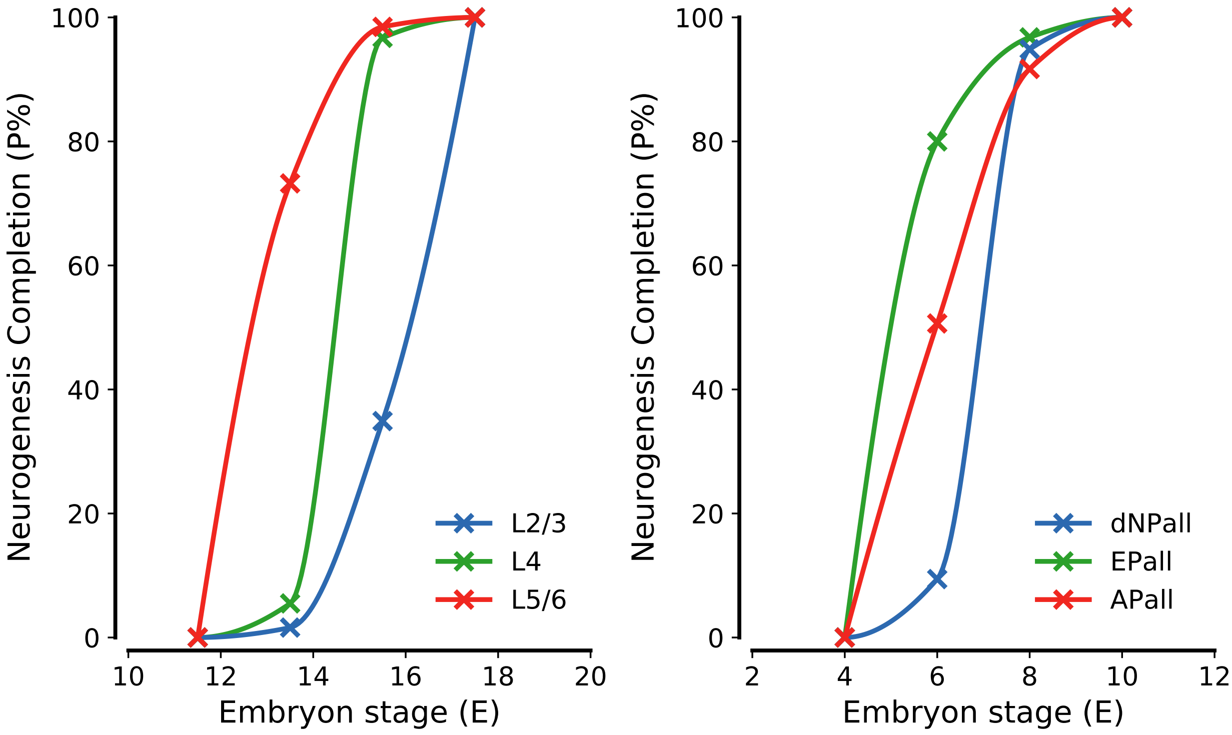

**Figure 1.** Neurogenesis in the chick and the mouse. Experimental data ($\times$) from Tables 1 and 2 are interpolated by a piecewise cubic hermite polynomial *solid curve* to illustrate the likely dynamics of neurogenesis. This latter curve will be used as the basis to deduce analytic expressions later on.

| *Region* | $t = 3.99$ | $t = 5.99$ | $t = 7.99$ | $t = 9.99$ |
|---|---|---|---|---|
| dNPall | 0.0 % | 9.4 % | 84.9 % | 100.0 % |
| EPall | 0.0 % | 80.0 % | 96.8 % | 100.0 % |
| APall | 0.0 % | 50.6 % | 91.7 % | 100.0 % |

**Table 3.** Cumulative percentiles of neurogenesis at the end of each development stage in each circuit relay in the chick. Data obtained by original experiments from Table 1. Time ($t$) units are embryon days (E#).

the probability of completion of the circuit at a given time. Such "probability of completion" can be regarded as the cumulative probability function (CDF) for neurogenesis, which we denote by $\Phi$. To seek an analytical expression for $\Phi$ from experimental data, we deploy Metalogistic (a.k.a. Metalog) fitting, motivating our approach by the unique shape and bounds flexibility of Metalog distributions among continuous distributions (Keelin, 2016).

We start from the consideration of the Piecewise Cubic Hermite Interpolant Polynomial (PCHIP) curve of our experimental data, which we denote by $N\%(t)$, then the experimental cumulative

| *Region* | $t = 11.49$ | $t = 13.49$ | $t = 15.49$ | $t = 17.49$ |
|---|---|---|---|---|
| L2/3 | 0.0 % | 1.6 % | 34.9 % | 100.0 % |
| L4 | 0.0 % | 5.5 % | 96.7 % | 100.0 % |
| L5/6 | 0.0 % | 73.2 % | 98.3 % | 100.0 % |

**Table 4.** Cumulative percentiles of neurogenesis at the end of each development stage in each circuit relay in the chick. Data obtained by original experiments. Data inferred from Table 2.

probability function of neurogenesis is given by

$$y = \Phi(t) = N\%(t)/100 \tag{1}$$

We aim to fit $\Phi(t)$ by a Metalog distribution. We do so, considering the quantile function $t = Q(y) = \Phi^{-1}(y)$ Specifically, we pick three quantile values that are symmetric with respect to $y = \Phi = 0.5$, namely $(t_\alpha, \alpha)$, $(t_{0.5}, 0.5)$, $(t_{1-\alpha}, 1-\alpha)$ with $0 < \alpha < 1$. In this way, we simplify our framework looking to fit our experimental data by a symmetric-percentile triple (SPT) Metalog distribution, whose quantile function reads (Keelin, 2016)

$$\hat{Q}(y) = \begin{cases} \dfrac{b_l + b_u \exp\left(M(y)\right)}{1 + \exp\left(M(y)\right)} & \text{for } 0 < y < 1 \\ b_l & \text{for } y = 0 \\ b_u & \text{for } y = 1 \end{cases} \tag{2}$$

with

$$M(y) = a_1 + (a_2 + a_3(y - 0.5)) \ln\left(\frac{y}{1-y}\right) \tag{3}$$

The advantage of this formulation, is that the SPT Metalog PDF is expressed in terms of quantiles, being

$$\hat{n}(y) = \begin{cases} m(y) \dfrac{\left(1 + \exp\left(M(y)\right)\right)^2}{b_u - b_l \exp\left(M(y)\right)} & \text{for } 0 < y < 1 \\ 0 & \text{for } y = 0, y = 1 \end{cases} \tag{4}$$

with

$$m(y) = \left(\frac{a_2}{y(1-y)} + a_3\left(\frac{y - 0.5}{y(1-y)} + \ln\left(\frac{y}{1-y}\right)\right)\right)^{-1} \tag{5}$$

In the previous expressions, $(b_l, b_u)$ define the time interval for neurogenesis, and respectively are $(3.99, 9.99)$ in the chick and $(11.49, 17.49)$ in the mouse (rounded to the second digit). The coefficients $a_1$, $a_2$, $a_3$ can instead be expressed as a function of the quantile triplet (Keelin, 2016):

$$a_1 = \ln \gamma_{0.5} \qquad a_2 = \frac{\ln\left(\frac{\gamma_{1-\alpha}}{\gamma_\alpha}\right)}{2\ln\left(\frac{1-\alpha}{\alpha}\right)} \qquad a_3 = \frac{\ln\left(\frac{\gamma_{1-\alpha}\gamma_\alpha}{\gamma_{0.5}}\right)}{(1-2\alpha)\ln\left(\frac{1-\alpha}{\alpha}\right)} \tag{6}$$

where $\gamma_\beta = \frac{t_\beta - b_l}{b_u - t_\beta}$ for $\beta = \alpha, 0.5, 1-\alpha$. The fitting procedure consists of estimating the coefficients $a_1$, $a_2$, $a_3$ from data by linear least squares based on the procedure described in (Keelin, 2016). This is practically implemented by the Python package `pymetalog` (v0.2.2, `https://github.com/tjefferies/pymetalog`). Figures 2 and 3 show the result of this fitting for the different region-specific CDFs (as $y$ vs. $\hat{Q}(y)$) and the corresponding PDFs (as $\hat{n}(y)$ vs. $\hat{Q}(y)$) for the chick and the mouse, respectively.

| *Region* | $a_1$ | $a_2$ | $a_3$ |
|---|---|---|---|
| dNPall | $-0.034$ | $0.262$ | $-0.051$ |
| EPall | $-1.623$ | $0.769$ | $-0.178$ |
| APall | $-0.714$ | $0.749$ | $-0.425$ |

**Table 5.** Metalog fitting coefficients for the chick neurogenesis data.

| *Region* | $a_1$ | $a_2$ | $a_3$ |
|---|---|---|---|
| L2/3 | $1.179$ | $0.738$ | $0.374$ |
| L4 | $-0.016$ | $0.223$ | $-0.013$ |
| L5/6 | $-1.428$ | $0.754$ | $-0.271$ |

**Table 6.** Metalog fitting coefficients for the mouse neurogenesis data.

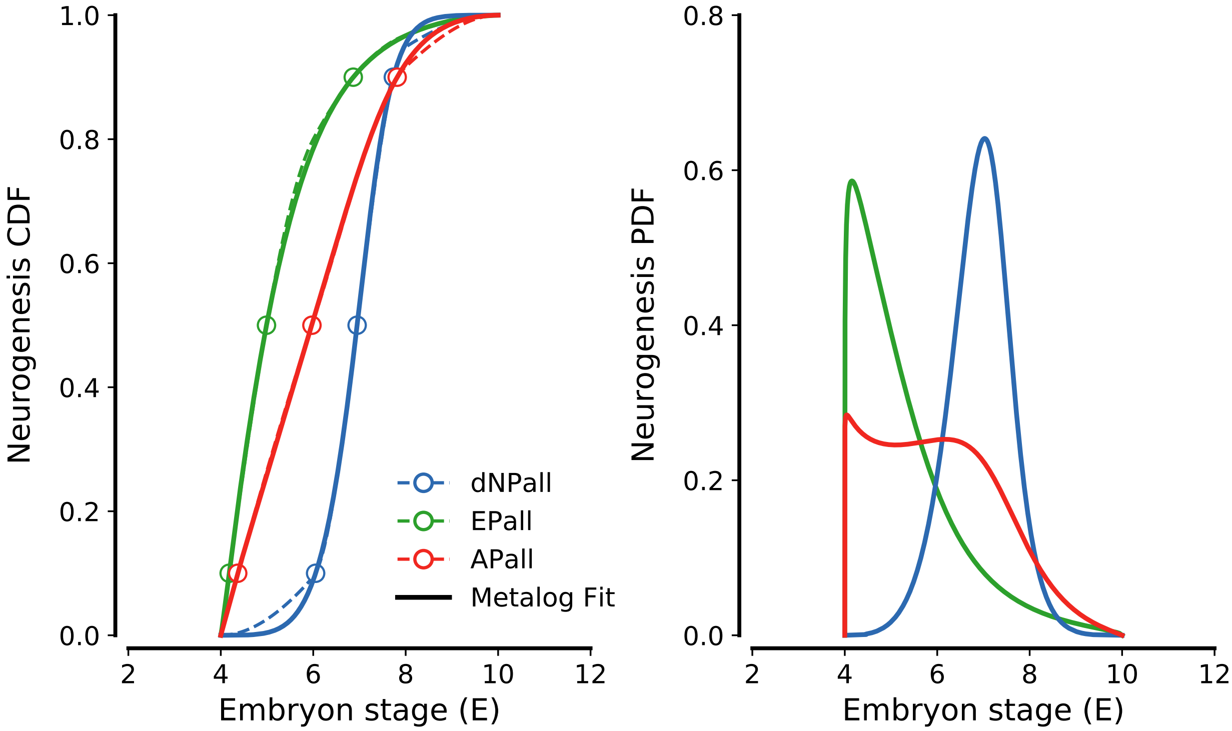

**Figure 2.** Neurogenesis model in the chick. (*Left panel*) Experimentally-derived region-specific CDFs (*dashed curves*) are fitted by SPT Metalog distributions (*solid curves*). (*Right panel*) Corresponding PDFs denoting the probability of neurogenesis within the development period are analytically constructed based on the prescribed procedure.

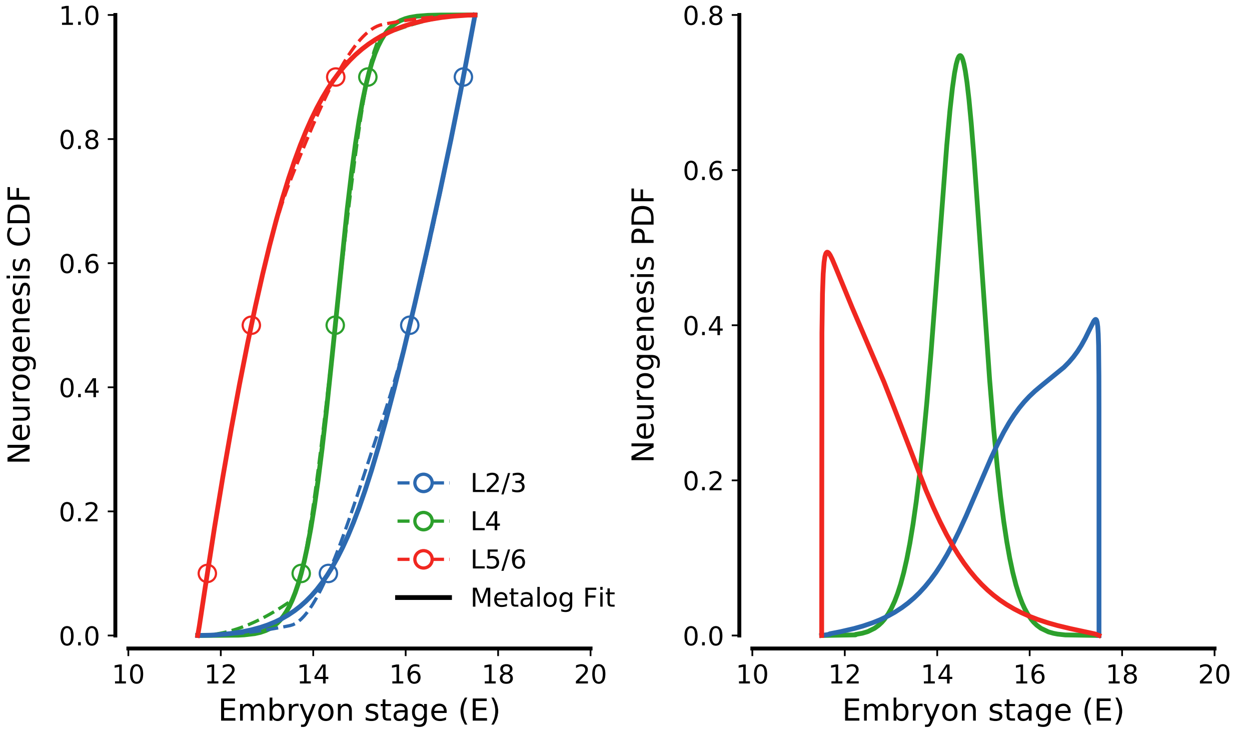

**Figure 3.** Neurogenesis model in the mouse. Legend as in Figure 2

### 2 Network model

#### 2.1 General framework

We consider a network of $N$ identical neurons. The activation $x_i(t)$ of the $i$th neuron determines its activity rate $\phi_i(t)$ by a nonlinear function $\phi_i(t) = \tanh(x_i)$, and depends on the activity of all presynaptic neurons connected to the neuron $i$ by the connection matrix $\mathbf{J}$, i.e.

$$\frac{\mathrm{d}}{\mathrm{d}t} x_i(t) = -x_i(t) + \sum_{j=1}^{N} J_{ij}\phi_j(t) \tag{7}$$

where $\phi_j(t) = \tanh(x_j)$. In our framework, neuronal activity can be equivalently interpreted either as the firing rate of the neuron, or as the elevation above baseline of the neuron's intracellular calcium, both of which may be representative readouts of embryonic activity (Spitzer, 2006).

The matrix $\mathbf{J}$ can be decomposed by the element-wise (or Hadamard) product of the network's adjacency matrix $\mathbf{A}$, whose binary elements $A_{ij}$ equal 1 only if cell $i$ is connected with $j$, and the weight matrix $\mathbf{G}$, whose elements are drawn from plausible weight distributions for $i$-to-$j$ connections. i.e.

$$\mathbf{J} = \mathbf{A} \circ \mathbf{G} \tag{8}$$

where '$\circ$' denotes the Hadamard product. This decomposition reflects the fact that synaptogenesis and synaptic maturation can be regarded, as least to a first order approximation, as separate processes (Blankenship et al., 2010; Morrison et al., 2008). In more general terms, in the embryo, establishment of a connection between two cells as a mean of coupling (i.e. correlating) their activities, can be mediated by volume signaling and gap junction coupling well before the actual maturation of a chemical synapse between those cells (Blankenship et al., 2010). Indeed, evidence in mouse embryos pinpoint to synaptic maturation to occur prominently postnatally (Blankenship et al., 2010; Martini et al., 2021). These arguments translate into the following modelling choices:

1. The existence of electrical rather than coupling between neurons in the embryo, accounts for undirected connections, namely $\mathbf{J}$, $\mathbf{A}$, and $\mathbf{G}$ are symmetric.

2. In the most general framework, establishment of a connection between two cells $i$ and $j$ may be thought to occur randomly, and be mediated by extracellular (volume) signal. Here we assume that the elements of $\mathbf{A}$ are binary values such that $A_{ij} \sim \mathrm{Bernoulli}(p)$, where $p$ is the probability of establishing a connection between cells $i$ and $j$. Generally, we assume that the connection probability within a pallial (respectively, cortical) area, that is the probability of recurrent connection $p_r$, is different from the probability of connection between different areas, i.e. $p_x$.

3. Electrical transmission can depolarize (excite) or hyperpolarize (inhibit) target cells, depending on intra- vs. extra-cellular chemical gradients. Accordingly, we consider $G_{ij} \sim \sigma W_{ij}$, where $\sigma$ is the weight standard deviation from zero, and $W_{ij} \sim \mathrm{Normal}(0, 1)$. Generally, we assume couplings within pallial (cortical) areas vs. coupling between different areas, could be generally different, and for this reason we consider weight deviations $\sigma_r$ for recurrent

| *Region* | Size ($f_\alpha$) |
|---|---|
| dNPall | 34.4 % |
| EPall | 22.2 % |
| APall | 43.4 % |

**Table 7.** Percentages of neurons in pallial areas at $t = 10$. Original data.

| *Region* | Size ($f_\alpha$) |
|---|---|
| L2/3 | 32.1 % |
| L4 | 24.7 % |
| L5/6 | 43.2 % |

**Table 8.** Percentages of neurons in cortical layers at $t = 17.5$. Data by [Meyer2010].

connections, and $\sigma_x$ for non-recurrent ones, respectively. Specifically, we make $\sigma$ scale as $g/\sqrt{N}$ to constrain the eigenvalue spectrum within a real axis interval of $O(1)$ (Ahmadian et al., 2015; Sommers et al., 1988; Sompolinsky et al., 1988).

To account for cell heterogeneity by area, we can group cells from the same area together, and envisage $\mathbf{J}$, $\mathbf{A}$, and $\mathbf{G}$ as block structured matrices of 4-by-4 blocks. In this fashion, we rewrite $\mathbf{J}$ in the form:

$$\mathbf{J} = \begin{pmatrix} \mathbf{J}_{11} & \mathbf{J}_{12} & \mathbf{J}_{13} & \mathbf{J}_{14} \\ \mathbf{J}_{11} & \mathbf{J}_{22} & \dots & \dots \\ \dots & \dots & \dots & \dots \\ \dots & \dots & \dots & \mathbf{J}_{44} \end{pmatrix} = \begin{pmatrix} \mathbf{A}_{11} \circ \mathbf{G}_{11} & \mathbf{A}_{12} \circ \mathbf{G}_{12} & \mathbf{A}_{13} \circ \mathbf{G}_{13} & \mathbf{A}_{14} \circ \mathbf{G}_{14} \\ \mathbf{A}_{11} \circ \mathbf{G}_{11} & \mathbf{A}_{22} \circ \mathbf{G}_{22} & \dots & \dots \\ \dots & \dots & \dots & \dots \\ \dots & \dots & \dots & \mathbf{A}_{44} \circ \mathbf{G}_{44} \end{pmatrix} \tag{9}$$

where the block matrices $\mathbf{A}_{kl}, \mathbf{G}_{kl}$ at the end of synaptogenesis are of size $f_k N \times f_l N$, where $f_\alpha$ represents the size of a region in terms of neuron counts with respect to total number of neurons in the pallium (respectively, the cortex).

#### 2.2 Organism-specific circuits

We look at thalamo-pallial and thalamo-cortical circuits from the mature brain of chicks and mice, respectively, to infer some constraints on the structure of $\mathbf{J}$ and its constituting block matrices $\mathbf{J}_{kl}$. We argue that, as connections develop between neurons in the embryo, they do so based on the circuits in Figures 4 and 5. In other words, based on the knowledge of avian thalamo-pallial and mouse thalamo-cortical in the mature brain, we set what connections can develop and which ones are prohibited instead. Accordingly, the connection matrices for the directed ($\rightarrow$) mature circuits are:

- in the avian brain:

$$\mathbf{J} = \begin{pmatrix} \mathbf{J}_{\mathrm{DT}\rightarrow\mathrm{DT}} & \mathbf{0} & \mathbf{J}_{\mathrm{DT}\rightarrow\mathrm{EP}} & \mathbf{0} \\ \mathbf{J}_{\mathrm{AP}\rightarrow\mathrm{DT}} & \mathbf{J}_{\mathrm{AP}\rightarrow\mathrm{AP}} & \mathbf{0} & \mathbf{J}_{\mathrm{AP}\rightarrow\mathrm{dNP}} \\ \mathbf{0} & \mathbf{0} & \mathbf{J}_{\mathrm{EP}\rightarrow\mathrm{EP}} & \mathbf{J}_{\mathrm{EP}\rightarrow\mathrm{dNP}} \\ \mathbf{0} & \mathbf{J}_{\mathrm{dNP}\rightarrow\mathrm{AP}} & \mathbf{J}_{\mathrm{dNP}\rightarrow\mathrm{EP}} & \mathbf{J}_{\mathrm{dNP}\rightarrow\mathrm{dNP}} \end{pmatrix} \tag{10}$$

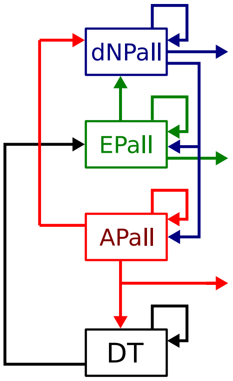

**Figure 4.** Thalamo-pallial circuit in the mature chick brain (inferred from Stacho et al., 2020).

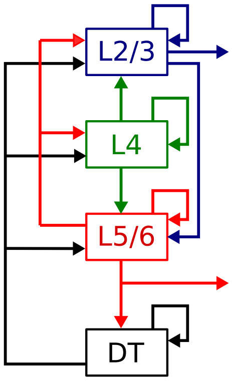

**Figure 5.** Thalamo-cortical circuit in the mature mouse brain (inferred from Thomson et al., 2003).

- in the mouse brain:

$$\mathbf{J} = \begin{pmatrix} \mathbf{J}_{\mathrm{DT}\to\mathrm{DT}} & \mathbf{J}_{\mathrm{DT}\to\mathrm{L5/6}} & \mathbf{J}_{\mathrm{DT}\to\mathrm{L4}} & \mathbf{J}_{\mathrm{DT}\to\mathrm{L2/3}} \\ \mathbf{J}_{\mathrm{L5/6}\to\mathrm{DT}} & \mathbf{J}_{\mathrm{L5/6}\to\mathrm{L5/6}} & \mathbf{J}_{\mathrm{L5/6}\to\mathrm{L4}} & \mathbf{J}_{\mathrm{L5/6}\to\mathrm{L2/3}} \\ \mathbf{0} & \mathbf{J}_{\mathrm{L4}\to\mathrm{L5/6}} & \mathbf{J}_{\mathrm{L4}\to\mathrm{L4}} & \mathbf{J}_{\mathrm{L4}\to\mathrm{L2/3}} \\ \mathbf{0} & \mathbf{J}_{\mathrm{L2/3}\to\mathrm{L5/6}} & \mathbf{0} & \mathbf{J}_{\mathrm{L2/3}\to\mathrm{L2/3}} \end{pmatrix} \tag{11}$$

It thus follow that, in the presence of undirected ($\leftrightarrow$) connections, the above matrices can be rewritten as

- in the avian brain:

$$\mathbf{J} = \begin{pmatrix} \mathbf{J}_{\mathrm{DT}\leftrightarrow\mathrm{DT}} & \mathbf{J}_{\mathrm{DT}\leftrightarrow\mathrm{AP}} & \mathbf{J}_{\mathrm{DT}\leftrightarrow\mathrm{EP}} & \mathbf{0} \\ \mathbf{J}^T_{\mathrm{DT}\leftrightarrow\mathrm{AP}} & \mathbf{J}_{\mathrm{AP}\leftrightarrow\mathrm{AP}} & \mathbf{0} & \mathbf{J}_{\mathrm{AP}\leftrightarrow\mathrm{dNP}} \\ \mathbf{J}^T_{\mathrm{DT}\leftrightarrow\mathrm{EP}} & \mathbf{0} & \mathbf{J}_{\mathrm{EP}\leftrightarrow\mathrm{EP}} & \mathbf{J}_{\mathrm{EP}\leftrightarrow\mathrm{dNP}} \\ \mathbf{0} & \mathbf{J}^T_{\mathrm{AP}\leftrightarrow\mathrm{dNP}} & \mathbf{J}^T_{\mathrm{EP}\leftrightarrow\mathrm{dNP}} & \mathbf{J}_{\mathrm{dNP}\leftrightarrow\mathrm{dNP}} \end{pmatrix} \tag{12}$$

- in the mouse brain:

$$\mathbf{J} = \begin{pmatrix} \mathbf{J}_{\mathrm{DT}\leftrightarrow\mathrm{DT}} & \mathbf{J}_{\mathrm{DT}\leftrightarrow\mathrm{L5/6}} & \mathbf{J}_{\mathrm{DT}\leftrightarrow\mathrm{L4}} & \mathbf{J}_{\mathrm{DT}\leftrightarrow\mathrm{L2/3}} \\ \mathbf{J}^T_{\mathrm{DT}\leftrightarrow\mathrm{L5/6}} & \mathbf{J}_{\mathrm{L5/6}\leftrightarrow\mathrm{L5/6}} & \mathbf{J}_{\mathrm{L5/6}\leftrightarrow\mathrm{L4}} & \mathbf{J}_{\mathrm{L5/6}\leftrightarrow\mathrm{L2/3}} \\ \mathbf{J}^T_{\mathrm{DT}\leftrightarrow\mathrm{L4}} & \mathbf{J}^T_{\mathrm{L5/6}\leftrightarrow\mathrm{L4}} & \mathbf{J}_{\mathrm{L4}\leftrightarrow\mathrm{L4}} & \mathbf{J}_{\mathrm{L4}\leftrightarrow\mathrm{L2/3}} \\ \mathbf{J}^T_{\mathrm{DT}\leftrightarrow\mathrm{L2/3}} & \mathbf{J}^T_{\mathrm{L5/6}\leftrightarrow\mathrm{L2/3}} & \mathbf{J}^T_{\mathrm{L2/3}\leftrightarrow\mathrm{L4}} & \mathbf{J}_{\mathrm{L2/3}\leftrightarrow\mathrm{L2/3}} \end{pmatrix} \tag{13}$$

#### 2.3 Simulating synaptogenesis

The mechanisms of embryonic synaptogenesis are little known, although emerging data suggest that it likely depends on neurogenesis (Blankenship et al., 2010). We mimic this possibility, considering the network immediately before birth (i.e. $t = t_\infty$), and randomly deleting neurons and their associated connections proceeding backward in time to earlier embryonic stages. Accordingly, we consider a succession of time points $t_0 < t_1 < \ldots < t_\infty$, at $t_i < t_\infty$, and extrapolate the fraction of neurons generated in each region $\alpha$ by fitting $\Phi_\alpha(t_i)$ (Section 1). As we proceed backward in time, at any instant $t_i$ we randomly pick and remove $\Delta N_\alpha(t_i)$ neurons and their connections, such that

$$\Delta N_\alpha(t_i) = N \cdot f_\alpha \cdot (\Phi_\alpha(t_{i-1}) - \Phi_\alpha(t_i)) = N \cdot r_\alpha(t_i) \tag{14}$$

where $r_\alpha(t_i) = f_\alpha \cdot (\Phi_\alpha(t_{i-1}) - \Phi_\alpha(t_i))$ denotes the time-dependent fraction of neurons generated for region $\alpha$. Although the random neuron selection can only account for specific realizations of the synaptogenic process, and to estimate the actual dynamics of synaptogenesis we should repeat our procedure over many realizations of the network at $t_\infty$, $r_\alpha$ is known regardless of the underlying distribution of connections. As discussed in the next section, this is all we need to estimate neural activity in our networks during embryogenesis.

#### 2.4 Analysis of neural dynamics

We consider the dynamic mean-field approach by (Aljadeff, Renfrew, et al., 2016; Aljadeff, Stern, et al., 2015) to study neural activity in our networks in the large $N$ limit. In this fashion, averaging equation 7 over the ensemble from which $\mathbf{J}$ is drawn implies that only neurons that belong to the same group are statistically identical. Thus, we can conveniently look at the activities $\xi_k(t)$ of four representatives neurons (each from one of the four regions in our circuits), and their inputs $\eta_k(t)$. In this way, the stochastic mean-field variables $\eta$ and $\eta$ can be adopted to approximate the activities and inputs in the full network, provided that they satisfy the dynamic equation

$$\frac{\mathrm{d}}{\mathrm{d}t}\xi_k(t) = -\xi_k(t) + \eta_k(t) \tag{15}$$

where $\eta_k(t)$ is a random variable with zero mean and cross-covariance (Aljadeff, Stern, et al., 2015)

$$\langle \eta_k(t)\eta_l(t+\tau)\rangle = \delta_{kl} \sum_m f_m G_{km}^2 C_m(\tau) \tag{16}$$

with $\delta_{kl}$ denoting the Kroenecker delta, and the indexes $k, l$, and $m$ running over $\mathrm{DT}, \mathrm{AP}, \mathrm{EP}, \mathrm{dNP}$ for the chick, and $\mathrm{DT}, \mathrm{L5/6}, \mathrm{L4}, \mathrm{L2/3}$ for the mouse. The average firing rate correlation matrix $\mathbf{C}(\tau)$ has components:

$$C_m(\tau) = \langle \phi_m(t)\phi_m(t+\tau)\rangle \tag{17}$$

and is linked with the cross-covariance by the equation (Aljadeff, Stern, et al., 2015)

$$\mathbf{H}(\tau) = \mathrm{diag}\left(\langle \eta_k(t)\eta_l(t+\tau)\rangle\right) = \mathbf{MC}(\tau) \tag{18}$$

where $M_{kl} = f_l G_{kl}^2$. In this fashion it can be shown that the eigenvalue spectrum of the $\mathbf{J}$ is limited by the Perron-Frobenious eigenvalue of the $\mathbf{M}$, that is its largest eigenvalue, which we denote by $\Lambda_1$ (Aljadeff, Renfrew, et al., 2016; Aljadeff, Stern, et al., 2015). As we set the connection gains to the constant values $g_r$ and $g_x$ for recurrent connections vs. non-recurrent ones, we study the developing network dynamics, considering $\Lambda_1(t)$ as $\mathbf{M}$ evolves during embryogenesis according to the formula

$$M_{kl}(t) = \varphi_l(t) G_{kl}^2 \tag{19}$$

where

$$\varphi_l(t) = \frac{f_l \Phi_l(t)}{\sum_m f_m \Phi_m(t)} \tag{20}$$

#### 2.5 Functional convergence

Based on the analysis of the eigenvalue spectrum, we look for functional convergence between the chick's thalamo-pallial circuit and the mouse thalamo-cortical one, as follows:

1. Comparing the time evolution of $\Lambda_1(t)$ in the two circuits, under the hypothesis that the thalamus size and synaptic connections are already in place at the onset of neurogenesis. In this respect, we take into account the observation that thalamic neural activity transitions from scant uncorrelated events to correlated dynamics at $t_\theta$ around E14.5 in the mouse (Martini et al., 2021) (and thus equivalently, at E7 in the chick), choosing $g_r/g_x$ such $\Lambda_1(t_\theta) \to 1$.

2. Comparing network dynamics as reflected by the loading matrix $\mathbf{L}(t) = \mathbf{\Lambda}(t)\mathbf{U}(t)$ where $\mathbf{\Lambda}, \mathbf{U}$ are the sorted eigenvalue-eigenvector pairs of $\mathbf{M}$ (Krzanowski, 1979). Procrustes analysis is then performed to evaluate how network dynamics in terms of eigenvectors weighted by their associatedd variance evolves during embryogenesis, towards the prenatal stage, i.e. as $t \to t_\infty$.
